# Evolution Shapes Cortical Interneuron Migration

**DOI:** 10.64898/2026.11.02.763736

**Authors:** Antonela Bonafina, Anaïs Boutsen, Haw-Yuan Chen, Miriam Javier-Torrent, Bernard Coumans, Ekaterina Epifanova, Adrian Cardenas, Romann Close, Hannah Bruce, Natali Rasetto, Sylvia Tielens, Christopher Kune, Isabel Maria Gallego-Lopez, Ryohei Iwata, Romain Le Bail, Sophie Perrier d’Hauterive, Anne Firquet, Silvia Cappello, Pierre Vanderhaeghen, Gabriel Mazzucchelli, Carsten Janke, Bernard Lakaye, Jean-Yves Raty, Victor Borrell, Laurent Nguyen

## Abstract

The cerebral cortex comprises excitatory neurons and inhibitory interneurons originating from distinct forebrain regions. During migration, cortical interneurons (cINs) interact dynamically with neighbouring cells and their microenvironment, contributing to cortical architecture. However, how evolutionary changes in cIN migration shape species-specific cortical development remains unclear. Comparative analyses in mouse, ferret, and human models revealed a conserved migratory sequence but distinct behaviours: human cINs migrate slower, display increased leading-process branching, and exhibit more frequent homotypic repulsive interactions than their mouse and ferret counterparts. These features promote dispersed cortical invasion of cINs in humans and ferrets, contrasting with the stream-like migration observed in mice. Mechanistically, elevated expression of the cytosolic carboxypeptidase CCP1 in human cINs reduces microtubule glutamylation and doublecortin recruitment, thereby facilitating leading-process branching. These findings identify a CCP1-dependent mechanism regulating cIN dispersion and suggest that evolutionary changes in cIN migration contribute to divergent patterns of cortical morphogenesis across mammals.

## Introduction

The cerebral cortex computes higher-order cognitive functions through complex neuronal circuits that are progressively assembled and refined after birth. These circuits comprise two major neuronal populations, the excitatory projection neurons (PNs), born from dorsal progenitors, and the inhibitory cortical interneurons (cINs). cINs are predominantly generated by ventral forebrain progenitors in the ganglionic eminences (GEs) from which they migrate tangentially to reach the developing cortex (*1*). They are subdivided into “cardinal” classes that are progressively refined into adult supertypes, accounting for the diversity of cINs in the adult cortex (*2*). Despite subtle differences in abundance, laminar distribution, and morphology, cIN diversity is overall conserved between mouse and human (*3*). However, while mouse cINs are generated exclusively in the ventral forebrain, some late born cINs arise from human pallial progenitors (*4–7*). These species-specific differences highlight a potential divergence in the developmental mechanisms underlying cortical circuit assembly between rodents and humans. Additional “evo-devo” differences may arise from species-specific features in the spatiotemporal establishment and integration of cINs within the cortical plate. The timing of cIN generation, migration, and integration is critical for circuit assembly and function, while differences in the number and pace of cINs assimilating into networks may have profound consequences on cortical morphogenesis such as gyrification as well as cerebral cortex connectivity and subsequent functional properties (*8*). Consistent with this view, emerging evidence suggests that cINs have increased in number and acquired greater integrative connectivity within human cortical circuits compared with those in the mouse (*9*).

Here, we tested the hypothesis that species differences in cortical network organization may be driven, at least in part, by evolutionary divergence in the migratory dynamics of cINs. We found that, while retaining the saltatory mode of migration observed in rodents (*10*), human cINs move slower and exhibit distinct pattern of nucleokinesis alongside more complex leading-process dynamics, characterized by increased branching and more frequent repulsive cell-cell interactions than their mouse counterparts. These differences in migratory dynamics prevent human cINs from organizing into the sharp migratory streams observed in rodents, instead resulting in a more dispersed pattern of cortical migration. Ferret cINs display an intermediate phenotype, consistent with an evolutionary transition toward increased dispersion. At the molecular level, human migrating cINs express higher levels of the cytosolic carboxypeptidase CCP1 than mouse cINs, resulting in reduced microtubule (MT) glutamylation. This reduction weakens the association of the MT-associated protein doublecortin (DCX) with MTs, thereby modulating leading-process stability and promoting branching. Consistent with this interspecies gradient, ferret cINs exhibit an intermediate migratory phenotype between mouse and human cINs, suggesting progressive differences in the regulation of DCX-MT interactions and their effects on migratory dynamics. Supporting a causal role for this mechanism, CCP1 overexpression in mouse cINs increases leading-process branching and apicobasal dispersion within the developing cortical wall, partially recapitulating the migratory behavior observed in human cINs.

Together, our findings identify a cell-intrinsic evolutionary mechanism linking CCP1-dependent regulation of MT dynamics to species-specific patterns of cIN invasion during cortical migration, providing a cellular and molecular basis for the diversification of cortical development and morphogenesis across mammals.

## Results

### Interneurons invade the cerebral cortex with species-specific migration dynamics

To enable a cross-species comparison of cIN migration dynamics, we first integrated distinct publicly available single-cell RNA-sequencing (scRNAseq) datasets, using corresponding developmental stages for embryonic mouse (embryonic day (E) 13.5-E14.5) and fetal human (gestational week (GW) 11-13) subpallial tissues (*11–15*) (Fig. 1A-1C). We focused the analysis on DLX5/6^+^ GABAergic neurons, which encompass post-mitotic cINs undergoing tangential migration to populate the cortical wall (*16*). DLX5/6 GABAergic neurons from different species showed overlapping transcriptional profiles with molecular signatures of their ganglionic eminence (GE) of origin. Based on regional GE signatures, we identified neural populations enriched with medial (MGE), caudal (CGE), and lateral ganglionic eminence (LGE) origins (Fig. 1D-1E; S1A). We next analyzed and compared the migration dynamics of DLX5/6 cINs as they populate the developing mouse or human cerebral cortex, focusing on the presumptive somatosensory cortex. To this end, we performed time-lapse imaging (16-24 h) of forebrain organotypic slices obtained from E13.5 mouse embryos or GW11-GW13 human fetuses. Slices were transduced with lentiviral reporters driven by the Dlx5/6 regulatory elements and expressing eGFP (Fig. 1F-1J; S1B-S1C). In both species, eGFP^+^ cINs were observed in the developing cortex, where they underwent saltatory migration, with human cINs showing lower average migration speed (Fig. 1G; Movies S1-S2) driven by less frequent nuclear translocation of comparable amplitude (nucleokinesis; Fig. 1H-1J) and longer nuclear pausing periods (Fig. 1I). To resolve fine-scale dynamics of leading-process branching that cannot be captured in organotypic slices, we imaged cINs migrating from mouse and human MGE explants (*17*) (Fig. 1K-1T; Movies S3-S4). Differences in average migration speed and nucleokinesis between species were consistent with those observed in forebrain organotypic slices (Fig. 1M-1P; S1E). Morphological analyses revealed that human cINs displayed greater maximal branching complexity than mouse cINs (Fig. 1Q; S1D), accompanied by increased growth-cone splitting (Fig. 1R) and a higher number of secondary and tertiary branches (Fig. 1S; S1F). These findings suggest enhanced branching activity in human cINs, despite a slight reduction of the maximal extension of the leading process (Fig. 1T) during migration compared with their mouse counterparts. Altogether, these analyses indicate that the overall migratory sequence of cINs is conserved across species, characterized by the coordinated alternation of leading-process extension and branching with nucleokinesis. However, human cINs exhibit distinct migratory features, including slower migration, and increased leading-process branching activity compared with mouse cINs.

**Figure 1.**
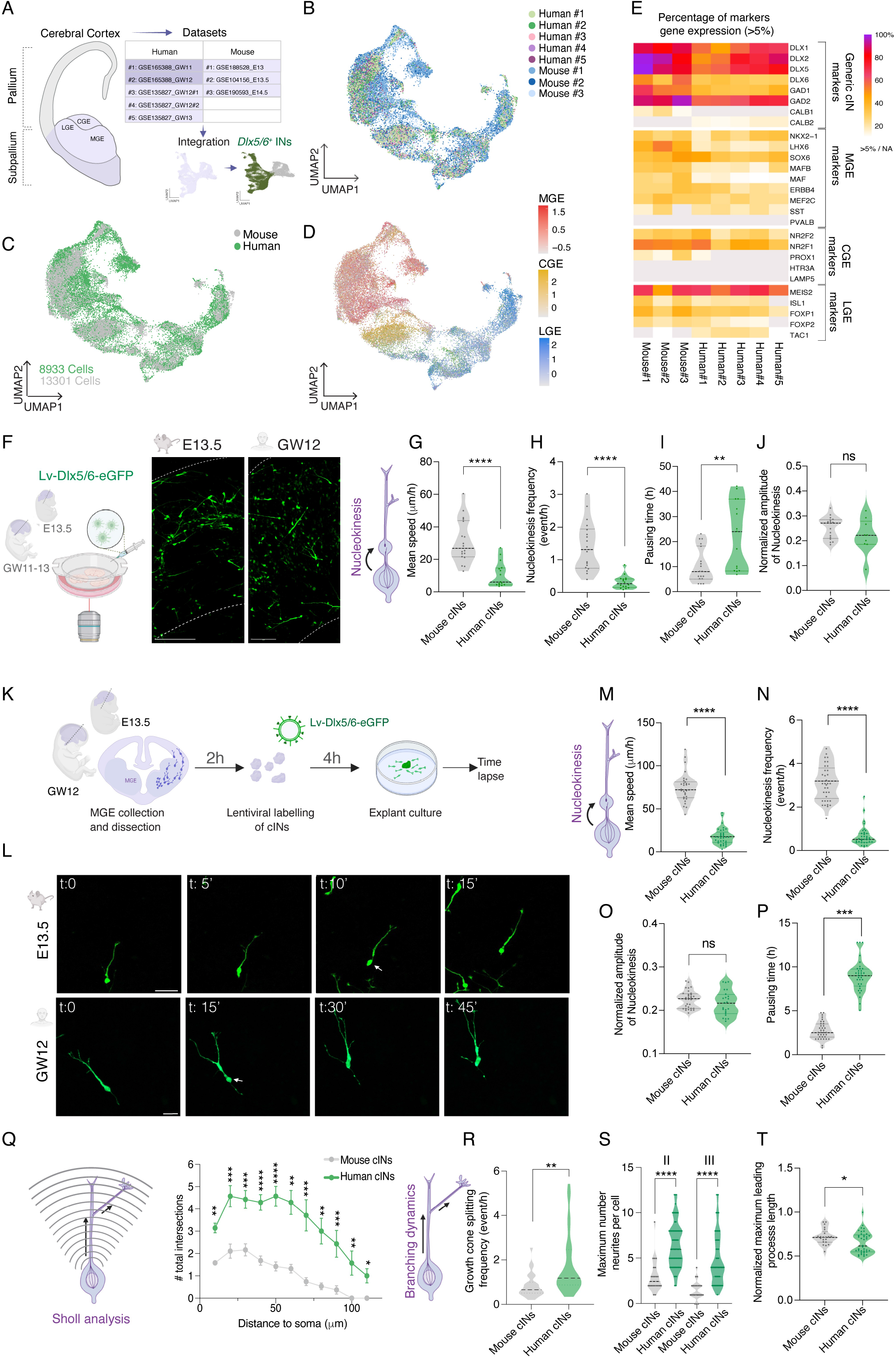
Human and mouse cINs exhibit distinct migration dynamics during their journey to the developing cerebral cortex. **A.** Single-cell RNA-sequencing workflow. Integration of 5 human and 3 mouse datasets spanning gestational weeks (GW) 11-13 (human) and embryonic days (E) 13-14.5 (mouse), respectively. *DLX5/6* interneurons (INs) cluster are isolated (dark green). MGE, medial ganglionic eminence ; LGE, lateral ganglionic eminence ; CGE, caudal ganglionic eminence. **B**-**D**. UMAP representations of integrated *DLX5/6* INs, annotated by sample identity (B), species (C), and expression level of ganglionic eminence (GE) specific markers (D). UMAP, uniform manifold approximation and projection. **E**. Heatmap of *DLX5/6* INs expressing selected genes associated with defined GE progenitors or IN subtypes, across datasets. Gray indicates expression below 5% threshold. **F**. Experimental setup for assessing cortical interneuron (cIN) migration within organotypic slices. Representative images of the cortical wall on forebrain organotypic slices from mouse (E13.5) and human (GW12) (right), labeled with Lv-Dlx5/6-eGFP. Scale bar, 50 μm. **G**-**I**. Quantification of nucleokinesis parameters in cINs migrating in mouse (E13.5) and human (GW11-13) organotypic slices after 2 days of culture in vitro (DIV) : average speed (G), nucleokinesis frequency (H), and nuclear pause duration (I). **J**. Amplitude of nucleokinesis normalized by cell size. Data represent between 15-18 cells from distinct mouse embryos (5) and human fetus (4 to 5) : ** p<0.01, ****p< 0.0001, ns not significant, unpaired t test. **K**. Experimental pipeline for time lapse recording of human and mouse cIN migrating from MGE explants. **L**. Time-lapse sequence illustrating the migration of a representative cIN in mouse and human MGE explant cultures. Scale bar, 50 μm. **M**-**P**. Quantification of nucleokinesis parameters of migrating cIN in MGE explant cultures. Distributions of mean migration speed (M), nucleokinesis frequency (N), normalized nucleokinesis amplitude (O), and nuclear pause duration (P). **Q**. Sholl analysis of migrating cIN at maximal branching complexity in mouse (E13.5) and human (GW12) explants (data represent 15 to 30 cells from distinct mouse embryos and human fetus (4 to 5)). **R**-**T**. Branching dynamics of migrating cINs. Violin plots show the frequency of growth cone splitting (**R**), the average maximum number of secondary (II) and tertiary (III) neurites (**S**), the normalized maximum leading process length (**T**). Data represent 15 to 35 cells from distinct mouse embryos (4–5) and human fetus (5 to 12) : ** p<0.01, ***p< 0.001, ns not significant, unpaired t test.

### CCP1 is a regulator of species-specific cortical interneuron migration dynamics

To identify molecular candidates underlying the species-specific differences of cIN migratory behavior, we compared the transcriptional profiles of postmitotic MGE-enriched *DLX5/6* cINs isolated from mouse and fetal human subpallial tissues using published transcriptomic datasets ((*11–15*); see also Fig. 1A). This analysis determined 2,967 differentially expressed genes (DEGs) between species (Fig. 2A, 2B), including 1,545 genes downregulated and 1,422 genes upregulated in human relative to mouse cINs (purple and orange, respectively). These findings revealed extensive transcriptional divergence between species within an otherwise “comparable” cIN population. We next performed Ingenuity Pathway Analysis (IPA) (*18*) to determine whether the transcriptional differences between mouse and human cINs converged on biological pathways and processes relevant to migration. Within the Disease and Functions analysis, we focused on terms related to Nervous System Development and Function. This analysis identified several functions associated with neural cell morphology, including neuritogenesis, branching of neurites, and dendritic growth/branching (Fig. 2C), consistent with the significant differences in branching dynamics that we observed across species (Fig. 1Q-1T). We next ran an over-representation analysis (ORA) of genes contributing to these IPA categories to determine whether specific biological pathways were enriched among the differentially expressed genes relative to chance expectation. This analysis highlighted cytoskeleton-related pathways, with multiple terms associated with MTs and actin organization & regulation among the most enriched categories (Fig. 2D and Fig. S2A). Given the central roles of MTs and actomyosin in nucleokinesis and leading-process branching during cIN migration (*19*), we focused subsequent analyses on molecules that coordinate such dynamics. Among them, cytosolic carboxypeptidase 1 (CCP1) encoded by *AGTPBP1*/*CCP1* (hereafter *CCP1*), a member of the CCP/AGBL family of deglutamylases, was compelling, as its activity regulates both nucleokinesis and leading-process branching in migrating mouse cINs (*20*) and this enzyme is immunodetected in human migrating cINs (Fig. S2B). Analysis of the selected scRNAseq datasets showed that *CCP1* was indeed the most broadly expressed member of its family in DLX5/6 cINs in both species. In addition, it showed markedly higher expression in human than in mouse cINs (Fig. 2E). Consistent with the scRNAseq analysis, human fetal MGE tissue extracts exhibited higher *CCP1* mRNA and protein levels (Fig. 2F-2H), accompanied by reduced levels of polyglutamylated MTs (PolyE), the main target of CCP1, as compared to mouse MGE extracts (Fig. 2G, 2I). Comparable differences were observed in individual cINs migrating in vitro (Fig. S2C-S2D) and in embryonic forebrain sections (human GW12 and mouse E13.5), where lower CCP1 expression was detected in mouse LHX6 cINs, both migrating from the GEs and those invading the developing cerebral cortex (Fig. 2J-2K). MT glutamylation is a reversible posttranslational modification depending on a balance between the activity of glutamylases (tubulin tyrosine-like ligases, TTLLs) and deglutamylases (CCPs). In addition to differences in *CCP1* expression the glutamylases encoded by *TTL1* and *TTL7* exhibit distinct species-specific expression patterns, which may contribute to interspecies differences in MT glutamylation levels (Fig. 2F; Fig. S2E-S2F).

**Figure 2.**
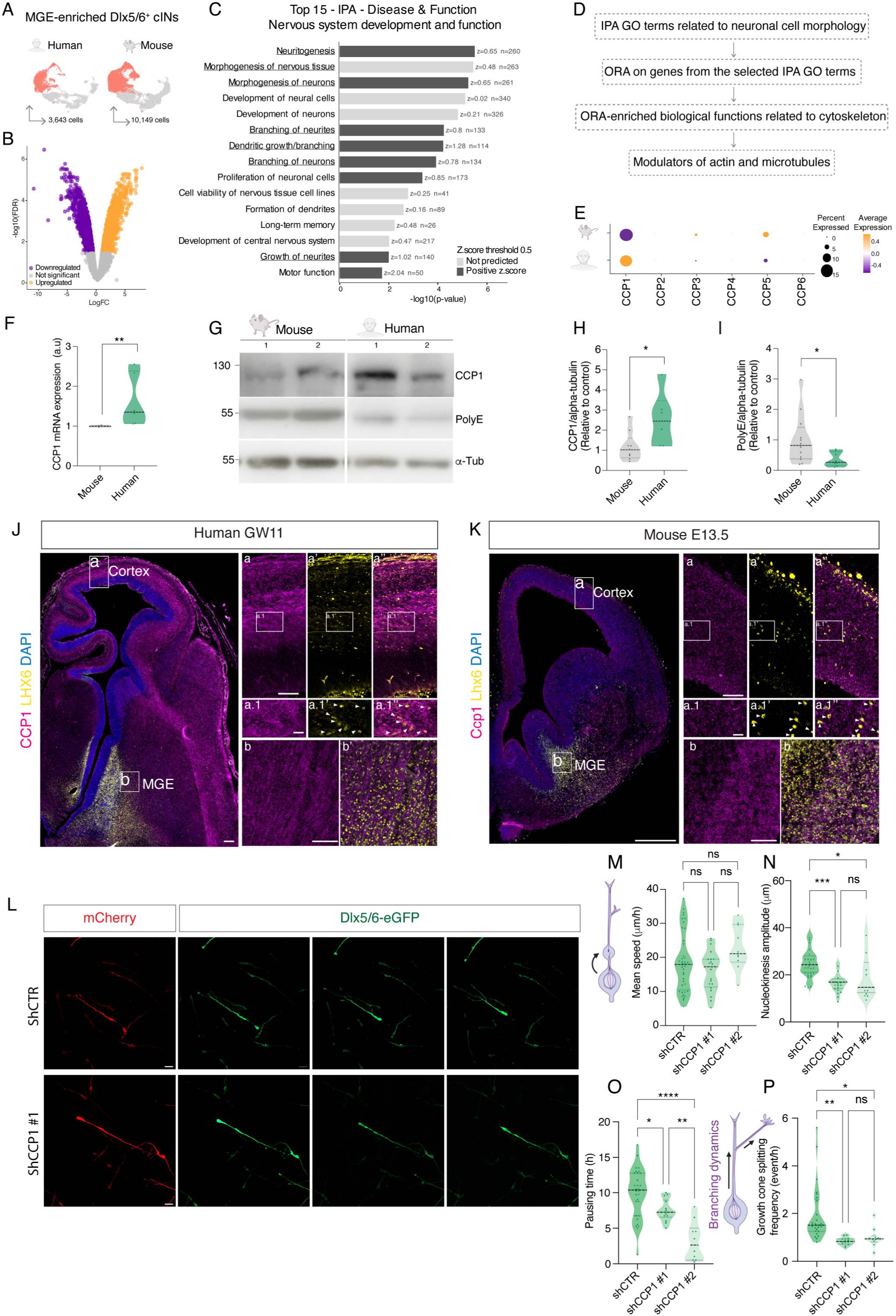
Elevated CCP1 expression drives human-specific cortical interneuron migration. **A**. UMAP representations of MGE-enriched *DLX5/6* cINs from human (left) and mouse (right) forebrains. MGE, medial ganglionic eminence ; UMAP, uniform manifold approximation and projection. **B**. Volcano plot of differentially expressed genes (DEGs) between human and mouse MGE-enriched *DLX5/6* cINs. Human genes significantly upregulated (orange), downregulated (purple) or not significantly different, as compared to mouse forebrain cINs (gray, FDR < 0.05; |log FC| > 0.25). **C**. Ingenuity Pathway Analysis (IPA) for Disease and Function enrichment of DEGs, filtered for *Nervous System Development and Function* categories. Bar color indicates the predicted activation state based on the IPA z-score (dark gray, positive z-score ; light gray, |z-score| < 0.5). *n* indicates the number of DEGs contributing to each term. Underlined terms were selected for downstream pathway analysis in (D). **D**. Workflow for the identification of candidate regulators of cIN migration. Over-representation analysis (ORA) was performed on genes contributing to the IPA related to cell morphology pathways. Gene Ontology (GO) terms associated with microtubule regulation are highlighted. **E**. Dot plot of expression of CCP/AGBL family deglutamylases in postmitotic MGE-enriched *Dlx5/6* cINs from mouse and human forebrains. Dot size indicates the percentage of expressing cells, and color represents scaled average expression. **F**. qRT-PCR analysis of *CCP1* expression in human (GW11-12 ; n=5) and mouse (E13.5; n=5) MGE samples, normalized to mouse expression (set to 1). *p<0.05, unpaired t test. **G**. Western blot of MGE protein lysates from human GW12 and mouse E13.5 embryos showing CCP1, polyglutamylated tubulin (PolyE), and α-tubulin (α-Tub) as a loading control. **H**, **I**. Quantification of CCP1 (H) and PolyE (I) immunoreactivity normalized to α-tubulin and expressed relative to mouse level in cINs (set to 1) (n=5 to 7 samples per group ; *p<0.05, unpaired t test). **J**, **K**. Immunolabeling of LHX6 (yellow), CCP1 (magenta), and DAPI (blue) in hemisection of GW11 human (J) and E13.5 mouse (K) forebrains. Insets show closer views on corresponding large image. Arrowheads point CCP1 /LHX6 cINs. Scale bars, 500 μm (main images), 100 μm (insets), and 20 μm (high-magnification views). **L.** Representative time-lapse sequences of GW12 human migrating cINs out of MGE explants and co-transduced with a Dlx5/6-eGFP lentivirus and an EF1α-dTomato lentivirus expressing either a control shRNA (shCTR; top) or a CCP1-targeting shRNA (shCCP1#1 ; bottom). Scale bar, 10 μm. **M**-**O**. Quantification of nucleokinesis parameters of migrating cINs in explant cultures transduced with control shRNA (shCTR) or CCP1-targeting shRNAs (shCCP1#1 and shCCP1#2). Violin plots show average migration speed (M), nucleokinesis amplitude (N), and nuclear pause duration (O) (n=15-30 cells from 4-5 GW11-12 human embryos, GW11-12) : *p<0.05, **p<0.01, ****p< 0.0001, One-Way ANOVA Tukey’s post-test. **P.** Quantification of branching dynamics in migrating human cINs. Violin plots show growth cone splitting frequency (n=15-30 cells from 3-4 GW11-12 human embryos; *p<0.05, **p<0.01, One-Way ANOVA Tukey’s post test.).

We next investigated the function of CCP1 in human cINs by expressing either control (shCTR) or two independent *CCP1*-targeting shRNAs (shCCP1#1 and shCCP1#2) (*21, 22*) in cINs migrating from GW11-12 MGE explants (Fig. S2G; 2L). CCP1 knockdown did not alter the mean migration speed but, it reduced nucleokinesis amplitude and shortened the nuclear pausing time. In parallel, CCP1 depletion in cINs reduced leading process branching dynamics, including growth cone splitting and neurite complexity (Fig. 2M-2P; S2H-S2I; Movies S5-S6). Given the role of MT glutamylation in regulating MT properties and interactions with MT-associated proteins (MAPs), the concomitant increase in CCP1 expression and reduction in PolyE levels in human cINs suggested the altered MT regulation is a key determinant of their distinctive migratory dynamics.

### Human forebrain assembloids as a model of CCP1-dependent cIN migration

Having identified CCP1 as a molecular candidate that drives cIN species-specific migratory behavior, we next sought to determine how CCP1 loss of function (LOF) affects cIN migration using human forebrain assembloids (hFAs) by dissecting the underlying molecular mechanisms. For this purpose, we generated human ventral forebrain organoids (hvFOs) (Fig. 3A) whose regional identity was validated by RT-qPCR (Fig. S3A-S3B). The protocol was applied to a panel of patient-derived CCP1 LOF and control human pluripotent stem cell lines (*23*) (Table S1), comprising three CCP1 mutant lines (CCP1^Mut#1-#3^) and four controls, including one hESC line (CCP1^WT#1^), two independent hiPSC lines (CCP1^WT#2-#3^), and an isogenic control of CCP1^Mut#2^ (CCP1^Iso#2^) (Table S1; Fig. 3B-3C; S3G). Most experiments compared CCP1^WT#2^, CCP1^Mut#2^, and CCP1^Iso#2^ lines, with the remaining lines used to confirm phenotype reproducibility. We next characterized the cellular composition and developmental trajectories of these hvFOs by scRNAseq (Fig. 3D-3H; Fig. S3C-S3F). Across cell lines, cells followed comparable developmental trajectories and gave rise to the major populations expected in hvFOs, including progenitors, dividing progenitors, oligodendrocyte precursor cells (OPCs), maturing interneurons, cortical-like interneurons (clINs), hindbrain-like interneurons (hilINs), and hypothalamic-like interneurons (hylINs) and a small pool of mature clINs (Fig. 3F-3G). Postmitotic DLX5/6 clINs were identified across all lines and selected for subsequent analyses (Fig. 3H). These cells exhibited cIN transcriptional identity, with comparable expression profiles across genotypes (Fig. 3D-3E, 3G; Fig. S3E). Mapping the identified DLX5/6 cell population onto a reference human forebrain dataset (human dataset on Fig. 1D) (*14, 15*) further confirmed their ventral forebrain identity, corresponding to MGE, LGE, and CGE populations (Fig. S3F). The conserved DLX5/6 clIN population across genotypes provided a robust cellular framework for investigating specific CCP1-dependent migratory phenotypes in a human model without evidence of cell fate alteration.

**Figure 3.**
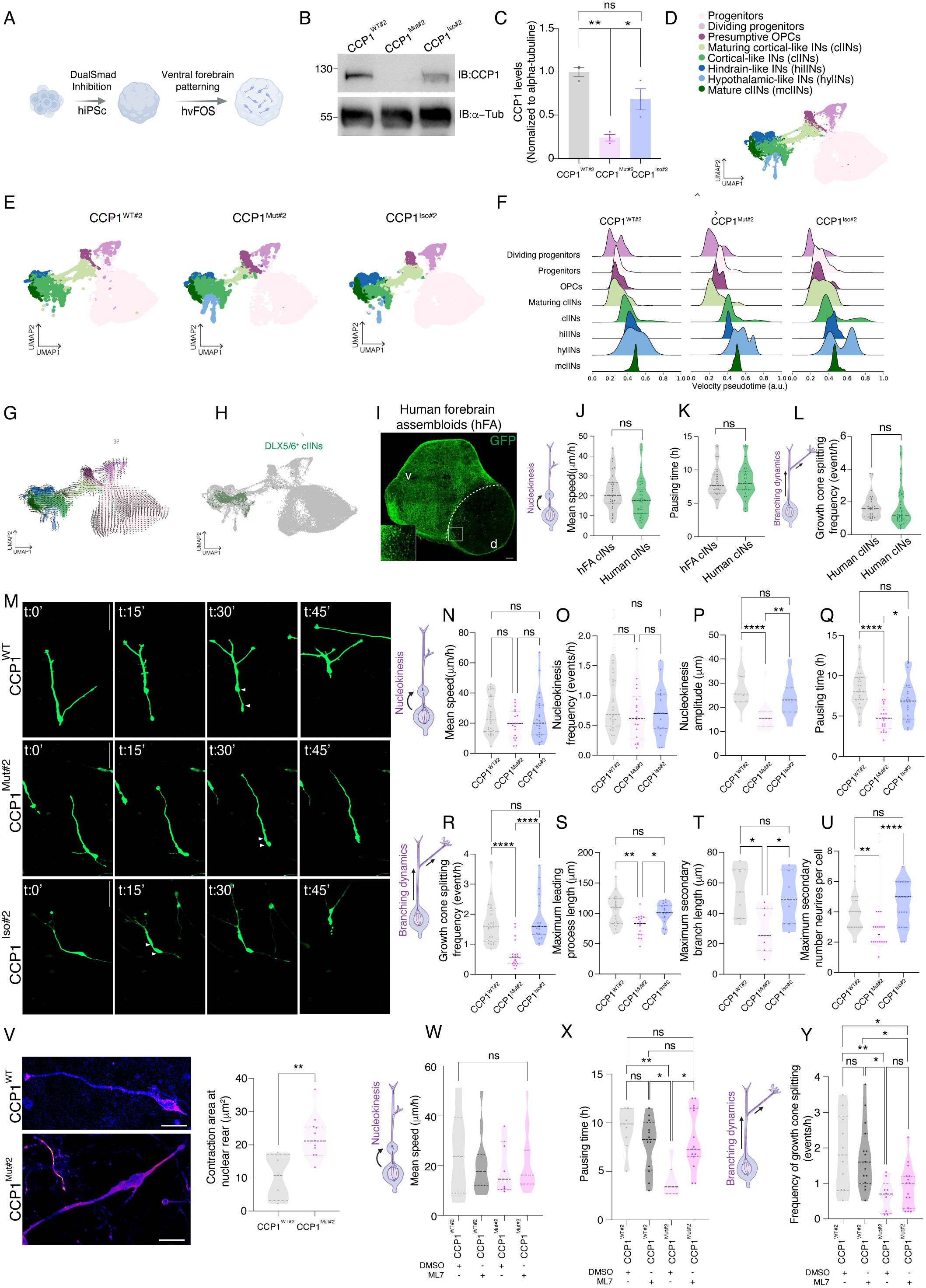
Human forebrain assembloids model the effects of CCP1 deficiency in cINs. **A**. Experimental workflow for the generation of human ventral forebrain organoids (hvFOs) from human induced pluripotent stem cells (hiPSCs). **B**-**C**. Representative western blot on 30 days hvFOs lysates showing CCP1 expression levels in CCP1^WT^, CCP1^Mut#2^, and CCP1^Iso#2^ lines. α-tubulin (α-Tub) serves as a loading control (B). Quantification of CCP1 protein levels normalized to α-tubulin and expressed relative to CCP1^WT#2^ level (set to 1) (n=3 independent experiments ; one-way ANOVA with Tukey’s multiple-comparison test, *p<0.05, **p<0.01) (C). **D-E**. Integrated UMAP from hvFOs, as annotated, showing cell population clusters, as reported in the figure (D). UMAP representations of scRNAseq data from 80-90 days hvFOs. Single cell transcriptomic contribution from CCP1^WT#2^ (control hiPSC line), CCP1^Mut#2^ (patient-derived hiPSC line carrying a loss-of-function mutation in CCP1), and CCP1^Iso#2^ (genome-edited isogenic line to CCP1^Mut#2^ hiPSCs) hvFOs to integrated UMAP (E). UMAP, uniform manifold approximation and projection ; OPCs, oligodendrocyte precursor cells ; INs, interneurons ; clINs, cortical-like interneurons ; hlINs, hindbrain-like interneurons ; hylINs, hypothalamic-like interneurons ; mclINs, mature cortical-like interneurons. **F-H**. Ridge plot comparing pseudotime distributions of identified cell populations between hvFOs lines, a.u., arbitrary units (F). RNA velocity analysis projected onto the integrated UMAP. Arrows indicate the predicted direction and magnitude of transcriptional state transitions (G). Dlx5/6 clINs are highlighted in green (H). **I**. Immunolabeling of a section of human forebrain assembloid cultured 90 days to highlight GFP^+^ clINs (green). Human forebrain assembloid (hFA) results from the fusion of a human dorsal forebrain organoid (hdFO, d) with a human ventral forebrain organoid (hvFO, v). Scale bar, 250 μm. **J**-**L**. Quantitative analysis of clIN migration in the dorsal region of hFA, compared with cIN migration in organotypic slices from human fetal cortices (see also Figure. 1F). Thirty days-old hvFOs were transduced with a Dlx5/6-eGFP lentiviral reporter before fusion and cultured for an additional 60 days prior to live imaging. Violin plots show mean migration speed (J), total pausing time (K), and growth cone splitting frequency (L). Data represent n=30-45 cells from 4-5 GW11-12 human embryos or 3-4 independent hFA differentiation assays (unpaired t test ; ns, not significant). **M**. Representative time-lapse sequences of migrating clINs generated in distinct hvFOs. clIN migration onto cortical feeder was monitored as cells exited the explant. Arrowheads indicate the soma and bulge positions prior to nucleokinesis. Scale bar, 50 μm. **N**-**Q**. Quantitative analysis of nucleokinesis dynamics of clINs during their migration out of hvFOs explants. Violin plots show mean migration speed (N), nucleokinesis frequency (O), nucleokinesis amplitude (P), and nuclear pause duration (Q). **R**-**U**. Quantitative analysis of branching dynamics of clINs during their migration from hvFOs explants. Violin plots show growth cone splitting frequency (R), maximum leading process length (S), maximum secondary neurites length (T), and maximum number of secondary neurites per cell (U) (n=23-30 cells from four independent experiments; one-way ANOVA, Tukey’s post-test ; ns, not significant; *p<0.05, **p<0.01, ***p<0.001). **V**. Representative images of migrating clINs derived from CCP1^WT#2^ and CCP1^mut#2^ hvFOs explants co-transduced with Dlx5/6-eGFP and Lv-PGK-LifeAct-Ruby (fire lookup table in ImageJ) (left). Scale bar, 20 μm. Quantification of maximum LifeAct-Ruby signal intensity at the nuclear rear during nucleokinesis (right, n=10-19 cells from three independent experiments; unpaired t test, **p<0.01). **W**-**Y**. Quantitative analyses of the migration dynamics of clINs derived from CCP1^WT^ and CCP1^Mut#2^ hvFOs treated with DMSO or ML-7, a MLCK inhibitor. Violin plots show mean migration speed (W), total pausing time (X), and growth cone splitting frequency (Y) (n=6-25 cells from at least three independent experiments; one-way ANOVA Tukey’s post-test ; ns, not significant ; *p<0.05; **p<0.01).

We next fused hvFOs with genotypically matched dorsal forebrain organoids (hdFOs) to check whether hFA-derived clINs recapitulate the migratory behavior of primary human cINs (Fig. 3I-3L; 1K-1T). We performed live imaging of clIN migration in organotypic hFA slices and found that their migratory behavior closely recapitulated that of GW11-GW12 primary human cINs migrating through the embryonic cortical wall (Fig. 3I-3L). Thus, hFA provided a tractable model to investigate the cellular consequences of CCP1 loss in clINs. Time lapse analyses on hFA organotypic slices showed that average clIN migration speed was unchanged upon CCP1 LOF, (Fig. S3H), however, the amplitude of nucleokinesis and nuclear pausing time were reduced (Fig. S3I, S3J). Such changes were specific to CCP1 loss of function as isogenic controls showed no defects (Fig. 3M-3Q). Analysis of hFA organotypic slices also showed impaired branching dynamics in CCP1-deficient clINs (Fig. S3K-S3M). To further examine leading process dynamics, we monitored Dlx5/6-eGFP clINs migrating from hvFO explants cultured on cortical feeder cells (Fig. 3M, 3R-3U). Compared with the control line, CCP1^Mut#2^ clINs exhibited a reduced frequency of growth cone splitting, shorter leading processes and secondary neurites, and fewer secondary neurites per cell. These phenotypes were not observed in the isogenic control of CCP1^Mut#2^ (Fig. 3M, 3R-U). Previous work identified the myosin light-chain kinase (MLCK) as a direct CCP1 target in mouse cINs, with its activity reliant on CCP1-dependant deglutamylation of its C-terminal polyglutamate chain. Loss of CCP1 in cINs impairs their nucleokinesis via increased actomyosin contractions, while the MLCK pharmacological inhibition with ML7 rescues this process in CCP1-deficient mouse cINs (*20*). We next asked whether CCP1 function in nucleokinesis regulation is conserved in human clINs. To this end, we performed time-lapse imaging and observed disrupted F-actin polarization, visualized using LifeAct-Ruby, at the rear of the nucleus, accompanied by reduced nuclear pausing time in CCP1^mut#2^ clINs compared with control clINs (Fig. 3V). MLCK inhibition with ML7 restored nuclear pausing time but did not rescue branching of the leading process (growth cone splitting) (Fig. 3W-3Y). These findings implicate MLCK in the nucleokinesis defects caused by loss of CCP1 activity, but not in the associated branching phenotype.

### CCP1 tunes human clINs dynamic branching through microtubule-DCX interactions

We next sought to define the molecular mechanism linking CCP1 loss to impaired leading-process branching in human clINs. To minimize hiPSC line-specific effects, we performed differential proteomic analysis on DLX5/6 clINs isolated from 90-day-old hFAs generated from CCP1^WT#2^ and pooled CCP1^Mut#1-#2^ lines (excluding CCP1^Mut#3^ which shows low *CCP1* expression) (*23*) (Fig. S3G, 4A). We identified 108 candidate proteins whose levels were significantly altered by CCP1 loss in clINs (53 down and 55 up) (Fig. 4B). IPA revealed two prominent pathways related to cell migration, encompassing cytoskeletal regulation and mitochondrial activity (Fig. S4A-S4C). Since there were no major mitochondrial dysfunctions or metabolic changes in clIN upon CCP1 LOF (Fig. S5A-S5N), we focused on cytoskeletal pathways given their likely contribution to the branching defects caused by CCP1 loss (S4B-S4C). Doublecortin (DCX), a MAP, was the most upregulated protein, not only amongst cytoskeletal proteins, but also amongst all upregulated proteins (Fig. 4C, S4A). The increased expression of DCX was further validated in CCP1 LOF hvFOs and was not attributable to changes in the transcript levels of its corresponding mRNA (Fig. S4D, S4E). Moreover, DCX expression was reduced upon isogenic correction of the CCP1 mutation (Fig. S4F). Given DCX’s role in MT regulation and neuronal morphogenesis (*24*), we asked whether its increased expression contributes to the branching defects resulting from CCP1 loss in clINs. Dlx5/6-eGFP clIN migration was analyzed by time-lapse microscopy after co-transduction of CCP1^WT#2^ and CCP1^Mut#2^ hvFO explants with a Dlx5/6-eGFP reporter lentivirus and a second lentiviral vector expressing shRNAs targeting DCX (shDCX#1 or shDCX#2) or a control shRNA (shCTR) (Fig. S4J; 4E-4F). DCX knockdown restored growth cone splitting frequency and branching complexity in CCP1^Mut#2^ clINs (Fig. 4F-4H). However, it did not rescue the altered nucleokinesis dynamics, including reduced nucleokinesis amplitude and shortened nuclear pausing time (Fig. 4I-4J). We therefore asked whether simultaneous modulation of DCX-and MLCK-dependent pathways could restore the full migratory phenotype. Combined DCX knockdown and MLCK inhibition with ML7 restored both branching and nucleokinesis in CCP1^Mut#2^ cINs, including nucleokinesis amplitude, nuclear pausing, and growth cone splitting frequency without affecting the mean speed (Fig. 4K-4M). These findings further identify distinct DCX- and MLCK-dependent pathways downstream of CCP1 that regulate leading-process branching and nucleokinesis, respectively.

**Figure 4.**
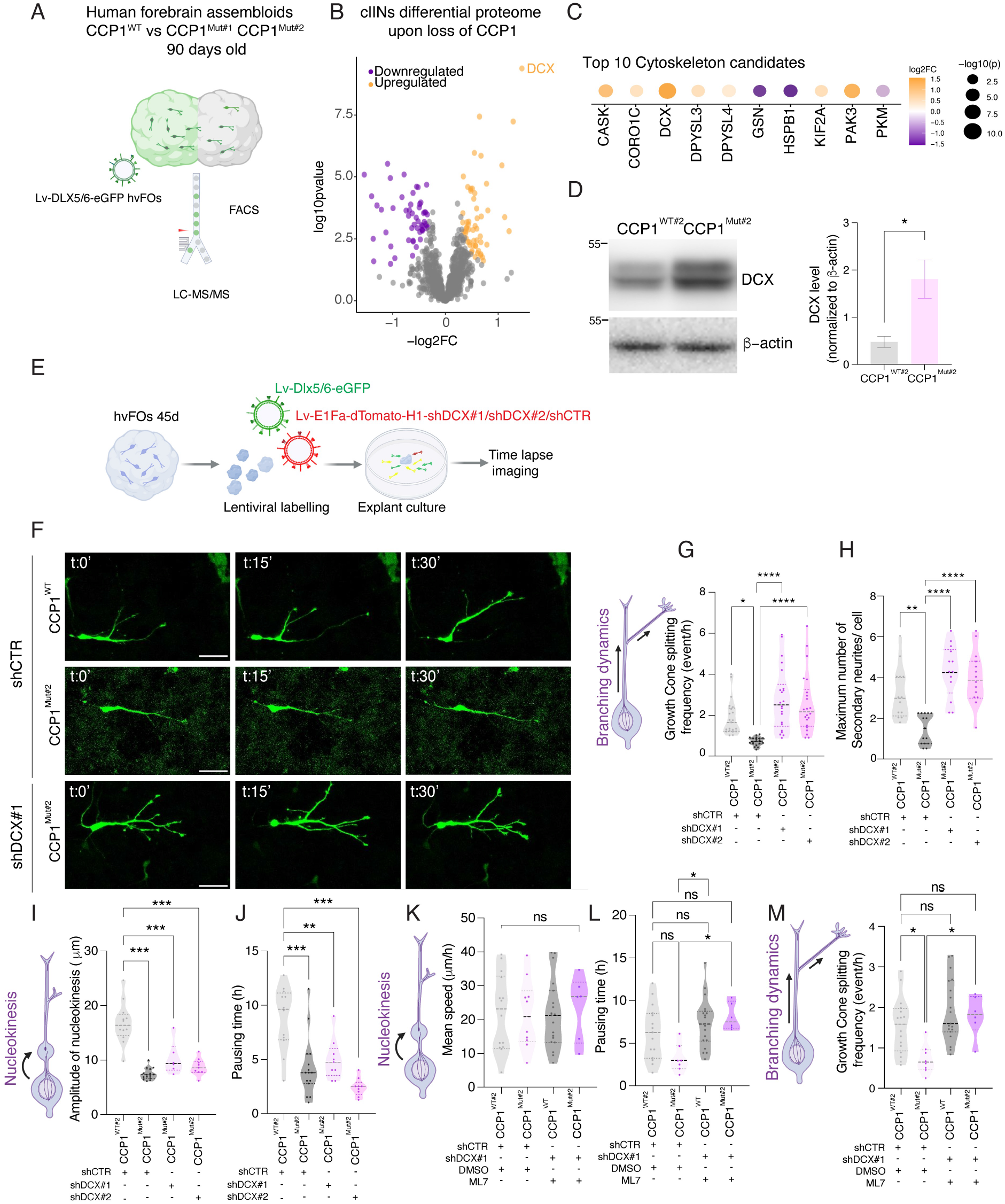
CCP1 drives cortical interneuron branching dynamics in part via DCX. **A**. Workflow for differential proteomic analysis of cortical-like interneurons (cINs) isolated from human forebrain assembloids (hFAs) generated using CCP1^WT#2^, CCP1^Mut#1^, and CCP1^Mut#2^ hiPSC lines. Human ventral forebrain organoids (hvFOs) were transduced with Dlx5/6-eGFP before fusion with human dorsal forebrain organoids (hdFOs). After 60 days, Dlx5/6-eGFP cINs were isolated by fluorescence-activated cell sorting (FACS) and analyzed by liquid chromatography-tandem mass spectrometry (LC-MS/MS). Differential protein expression was assessed by comparing CCP1^WT#2^ with pooled CCP1^Mut#1^ and CCP1^Mut#2^ samples. **B.** Volcano plot of differentially expressed proteins between CCP1^WT#2^ and pooled CCP1^Mut#1^ and CCP1^Mut#2^ samples. Proteins significantly upregulated in mutant clINs (orange), downregulated proteins (purple), and nonsignificant proteins (gray) are shown (*p< 0.05, |log FC| > 0.58). **C**. Dot plots of the top dysregulated proteins within the cytoskeleton category. Dot size indicates statistical significance (-log P), and color represents the magnitude and direction of differential expression (log fold change). **D**. Western blot showing DCX expression in lysates from 60 days CCP1^WT#2^ and CCP1^mut#2^ hvFOs (left). Quantification of DCX protein levels in the corresponding hvFOs extracts, normalized to β-actin levels (right). **E**. Experimental setup for live-cell imaging of hvFOs derived from the CCP1^WT#2^ and CCP1^Mut#2^ lines, co-transduced with a Dlx5/6-eGFP lentivirus and an EF1α-dTomato lentivirus expressing either a control shRNA (shCTR) or DCX-targeting shRNAs (shDCX#1 or shDCX#2). Time-lapse imaging was performed on yellow cortical-like interneurons (clINs) migrating out of the hvFOs explants. **F**. Representative time-lapse sequences of migrating GFP+ clINs (green), transduced with shRNA control or against DCX as indicated. Scale bar, 50 μm. **G**-**J**. Quantitative analysis of nucleokinesis and branching dynamics of migrating clINs derived from CCP1^WT#2^ and CCP1^Mut#2^ hvFOs transduced with either control shRNA (shCTR) or one of the two Dcx-targeting shRNA (shDCX#1). Violin plots show growth cone splitting frequency (G), maximum number of secondary neurites per cell (H), amplitude of nucleokinesis (I), and nuclear pause duration (J) (n=15-32 cells from three independent trials ; one-way ANOVA with Tukey’s multiple-comparison test ; ns, not significant; *p<0.05, **p<0.01, ****p< 0.0001). **K**-**M**. Quantitative analysis of the migration dynamics of clINs derived from CCP1^WT#2^ and CCP1^Mut#2^ hvFOs transduced with either control shRNA or shDCX and treated with the MLCK inhibitor ML7. Violin plots show mean migration speed (K), nuclear pause duration (L), and growth cone splitting frequency (M) (n=15-32 cells from three independent assays ; one-way ANOVA with Tukey’s multiple-comparison test ; ns, not significant; *p<0.05).

### CCP1 tunes doublecortin association to polyglutamylated microtubules

Having identified DCX as a downstream effector of CCP1-dependent leading process branching, we next asked how CCP1 regulates DCX function. DCX is a MAP that promotes MT polymerization and stability and supporting neuronal migration and morphogenesis (*25, 26*), including leading-process branching in migrating cINs (*24*). Recent work further linked DCX to tubulin polyglutamylation and neuronal branching, suggesting an interplay between DCX and MT post-translational modifications (*27*). Given CCP1’s role in regulating tubulin glutamylation, we therefore asked whether CCP1-dependent changes in MT glutamylation affect DCX-MT interactions. To test this, we performed MT cosedimentation assays using taxol stabilized extracts from 45 to 60-day-old CCP1^WT#2^, CCP1^Mut#2^, and CCP1^Iso#2^ hvFOs (Fig. 5A). DCX was detected in the polymerized MT fraction (P3, Fig. 5A-5B), with CCP1^Mut#2^ showing increased DCX association with polymerized MTs compared with CCP1^WT#2^. Such increase association between DCX and polymerized MTs was not observed in extracts from the isogenic CCP1^Iso#2^ line (Fig. 5B-5C). The elevated DCX levels detected upon CCP1 LOF may result from a stronger binding to hyperglutamylated MTs, thereby promoting a MT-stabilizing activity that reduces leading-process branching in CCP1 patient-derived clINs (Fig. S3G). While CCP1 is expressed in both mouse and human migrating cINs/clINs, their leading-process branching is more prominent in human than mouse (Fig. 1Q-1T). Moreover, since CCP1 LOF reduces branching in human cINs/clINs to mouse-like levels, we asked whether species differences in CCP1 expression might act as a rheostat to control leading-process branching via the MT recruitment of DCX. Our analyses showed that CCP1 expression is lower in mouse than human cINs (Fig. 2H, 2I). In addition, neither DCX protein nor mRNA levels are affected by CCP1 loss in mouse cINs (Fig. S4G-S4I). Along these lines, quantification of DCX fluorescence associated with βIII-tubulin revealed significantly higher MT-associated DCX levels in mouse than human cINs (Fig. 5D-5E). This was further confirmed by proximity ligation assay (PLA; Fig. 5F-5G) between DCX and βIII-tubulin, consistent with differential DCX recruitment to MTs in clINs/cINs, across species. To functionally test whether increased CCP1 expression alters DCX recruitment to MTs, we overexpressed either wild-type mouse CCP1 (CCP1^WT^) or the catalytically inactive H912S/E915Q mutant (CCP1^dead^) (*28*) in mouse cINs and showed increased DCX levels and association with MTs (Fig. 5H-5I). Together, these findings suggest that species differences in CCP1-dependent MT glutamylation are associated with differential DCX recruitment to MT, providing a molecular link between CCP1 expression levels and distinct branching behaviors of human and mouse clINs/cINs.

**Figure 5.**
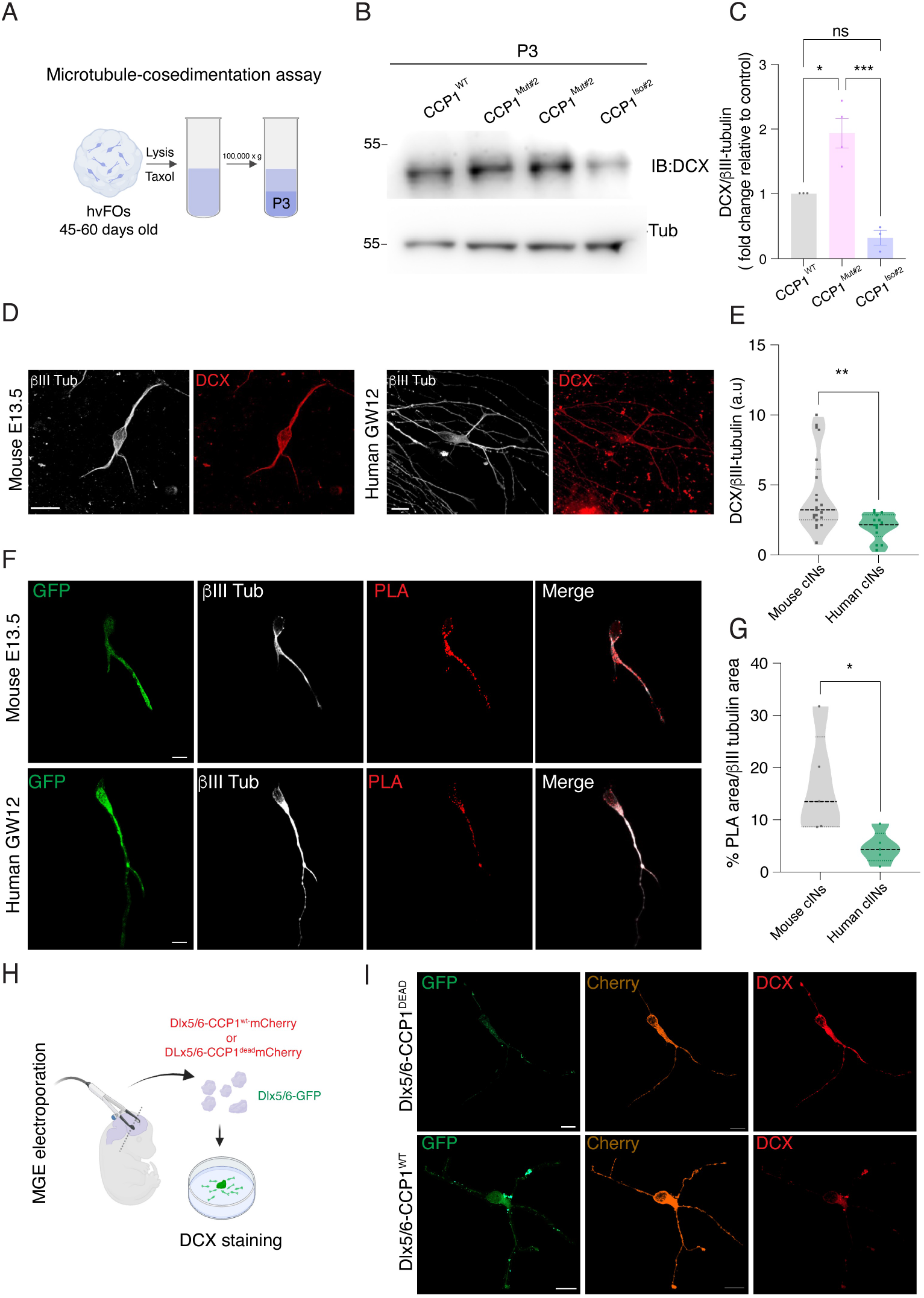
CCP1 tunes the recruitment of doublecortin to polygutamylated microtubules. **A.** Experimental set up for microtubule co-sedimentation assay using taxol-stabilized extracts from of CCP1^WT#2^, CCP1^Mut#2^ ventral forebrain organoid (hvFOs), and corresponding isogenic control (CCP1^Iso#2^) line. **B.** Western blotting for the detection of DCX and βIII-tubulin on lysates of CCP1^WT#2^, CCP1^Mut#2^, and corresponding isogenic control (CCP1^Iso#2^) hvFOs subjected to MT co-sedimentation. N = 3 to 4 independent organoid cultures. **C.** Western blot quantification of DCX levels normalized to βIII-tubulin in the polymerized microtubule fraction and expressed as fold change relative to CCP1^WT#2^ (*n* = 3-4 independent organoid cultures ; one-way ANOVA with Tukey’s multiple-comparison test ; *p*<*0.05). **D** Representative immunolabeling of DCX/Dcx (red) and βIII-tubulin (grey) in primary cINs that have migrated from E13.5 mouse or GW12 human MGE explants. **E**. Quantification of DCX immunofluorescence intensity normalized to βIII-tubulin in mouse and human cINs (unpaired t test ; *p< 0.05). **F**-**G**. Proximity ligation assay labeling (PLA, red) on Dlx5/6-eGFP-transduced primary cINs (green) that have migrated from E13.5 mouse or GW12 human MGE explants to detect proximity between βIII-tubulin (grey) and DCX (F). Quantification of PLA signal as ration between percentage of PLA area on βIII-tubulin area (unpaired t test ; *p< 0.05). **H-I**. Ex vivo electroporation of E13.5 mouse MGE with vectors (as indicated on the figure) and cultured as explants (H). Immunolabeling of cIN migrating out of MGE explant culture showing GFP (green), CCP1-Cherry (orange), and Dcx (red) (I).

### Species-specific cortical wall invasion pattern by interneurons relies on homotypic contacts

The migration routes of cINs into the cortical wall rely on physical and chemical cues sensed by their branching leading processes (*29, 30*). In mouse, cINs migrate tangentially together in organized streams. On the basis that leading-process branching is increased in human versus mouse cINs, we asked whether the pattern of cortical wall invasion is conserved across species. Live imaging of cINs migrating from MGE explants revealed more frequent homotypic contacts between human cINs as compared to mouse ones, with most contacts being mutually repulsive in human cINs but neutral in mouse cINs (18.5% vs. 66.7% repulsive contacts, respectively; Fig. 6A-6D; Movies S3-S4). As leading-process branching dynamics are regulated by CCP1 expression in human cINs (Fig. 3R-3U; Fig. S3K-S3M), and because mouse cINs express lower physiological level of CCP1 as compared to human ones (Fig. 2G-2H), we hypothesized that overexpression of CCP1 in mouse cINs may alter the frequency and nature of homotypic contacts. We generated lentiviruses expressing CCP1^WT^ or the catalytically inactive CCP1^dead^ that we co-transduced with Dlx5/6-eGFP reporter in mouse MGE explants (Fig. 6E; S6A-S6B). Live imaging revealed similar migration speeds and nucleokinesis frequencies across conditions, whereas CCP1^WT^ cINs showed longer nuclear pauses and increased growth cone splitting with higher branching complexity compared to CCP1^dead^ cINs (Fig. 6F; S6C-S6H). Notably, CCP1^WT^ cINs exhibited more frequent homotypic contacts and a higher proportion of repulsive responses than CCP1^dead^ cINs (39% vs. 15%; Fig. 6G-6H). While increased CCP1 activity in mouse cINs shifted branching dynamics and homotypic interactions toward the human phenotype, mutual repulsive contacts remained less frequent than observed between human cINs despite comparable cell densities across species (Fig. 6C, 6H). This suggests that mutual repulsion relies not only on branching and homotypic contact frequencies, but also on molecular mechanisms that may be human-specific. We performed CellChat analysis on cINs identified from publicly available scRNAseq datasets from the developing mouse and human cortical wall, integrating 16 mouse datasets spanning E13, E13.5, and E14.5 (*31–36*) and three human datasets at GW13 (*37, 38*), focusing our analysis on cIN-cIN communication (Fig. S7A-S7C). We identified 32 shared cell-cell interactions, along with 7 mouse-specific and 111 human-specific interactions (Fig. S7D-S7F), nominating putative candidate molecular signals underlying species-specific differences in homotypic cIN interactions during cortical migration.

**Figure 6.**
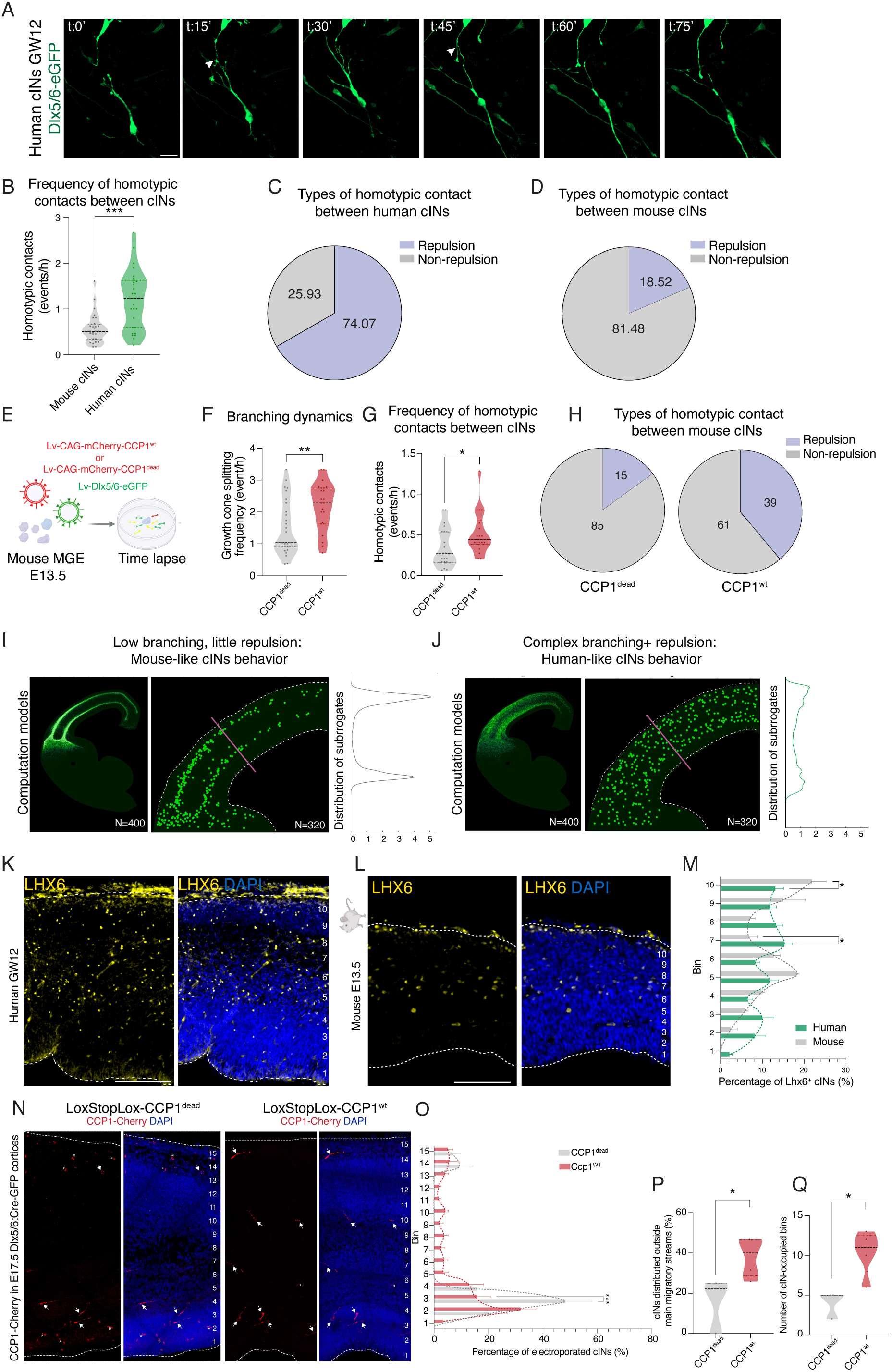
Species-specific cortical wall invasion patterns of interneurons rely on homotypic contacts. **A**. Representative time-lapse sequence of a homotypic contact between migrating human cortical interneurons (cINs) derived from a GW12 MGE explant. The arrowhead indicates the site of contact between two cINs. Scale bar, 20 μm. **B.** Frequency of homotypic contacts between migrating cINs per hour, quantified from time-lapse recordings of E13.5 mouse and GW11-12 human MGE explants (unpaired t test ; *p< 0.05; n = 25-30 cells from three independent experiments). **C-D.** Types of homotypic contacts between migrating cINs. Pie charts show the percentage of repulsive and neutral (non-repulsive) contacts in human cINs (D ; n = 27 cells from three independent experiments) and mouse cINs (E ; n = 27 cells from three independent experiments). **E.** Experimental pipeline for time lapse recording of migrating cINs from MGE explants after their transduction with Lv-Dlx5/6-eGFP and either Lv-CAG-Cherry-mCCP1 (CCP1^wt^) or Lv-CAG-Cherry-mCCP1dead (CCP1^dead^). **F.** Quantification of growth cone splitting frequency in rodent migrating CCP1^WT^ and CCP1^dead^ cINs (unpaired t test ; *p < 0.05; n = 35-45 cells). **G.** Frequency of homotypic contacts between cINs per hour (time lapse recording) in mouse CCP1^wt^ or CCP1^dead^ cINs. Unpaired t test ; *p<0.05 (n=25/30 cells). **H**. Pie charts show the percentage of repulsive and neutral (non-repulsive) contacts between transduced (condition, as indicated) mouse cINs (n = 22 cells). **I-J.** Dynamical simulation for Human and Mouse models using homemade code (Reference DOI: 10.5281/zenodo.22869350). Surrogates are generated in the sub-pallium and let to diffuse to the cortical wall in 2D. Left panels: trajectory points for the first 400 configurations observed after 65% of surrogates have crossed the basis line of the CP. Lighter green corresponds to higher probability density; same color scale for both models. Right panels: Single configuration observed in the CP after almost complete diffusion of surrogates. Side panels show the normalized (5px smoothing) apico-basal distribution (purple line) of the probability of presence of surrogates cINs as a function of the cortcial wall depth. **K-L.** Immunolabeling of the distribution of LHX6 cINs in embryonic cortical sections from human (GW12) (K) and mouse (E13.5) (L). Scale bars, 50 μm. **M.** Quantification of cIN distribution in embryonic mouse (grey) and human (green) cortical sections (two-way ANOVA with Tukey’s multiple-comparison test ; *p < 0.05; n = 3 to 4 embryos per conditions). **N-Q.** In utero electroporation of the ventral forebrain of E13.5Dlx5/6 :Cre-GFP with pCAG-LoxSTOPlox-Cherry-CCP1^WT^ or pCAG-LoxSTOPlox-Cherry-CCP1^Dead^ to target newborn cINs followed by brain collection at E17.5 to analyse the cortical distribution of transduced cINs. Arrows point to cINs and asterisks show non-specific staining of blood vessels. Scale bars, 50μm (N). Cortical bin distribution at E17.5 of transduced cINs with pCMV-LoxSTOPlox-CCP1^WT^ or pCMV-LoxSTOPlox-CCP1^Dead^ at E13.5 (O). Violins plots showing transduced cINs distributed outside the main migratory streams, as defined in mouse forebrain (P), and the number of cIN-occupied bins (Q).

We next asked whether species-specific differences in cIN branching and homotypic interactions influence the cortical invasion of cINs. We hypothesized that increased leading-process complexity and repulsive homotypic interactions could alter the distribution of migrating cINs within the developing cortex. To test this hypothesis, we used a computational model that placed migrating cIN surrogates (dynamically branching objects) within a cortical-like 2D space defined by a predetermined affinity field (Fig. 6I). The model incorporated experimentally measured migration parameters (Table S2), including leading-process branching and homotypic repulsion, which were tuned to reproduce mouse or human cIN behavior while keeping cortical geometry and the affinity landscape constant. cIN surrogates with mouse-like parameters formed two compact tangential streams, consistent with cIN migration streams in the marginal zone (MZ) and at the border of the subventricular and intermediate zones (SVZ/IZ) as described in the mouse cortex (*1, 39*). In contrast, increasing leading-process branching and homotypic repulsion modelling to mimic human cIN behavior disrupted stream formation, promoting a tangential dispersion of migrating surrogates across the cortical-like space (Fig. 6J; Movie S7). This prediction prompted us to examine the migration of cIN populations in stage-matched mouse (E13.5) and human (GW11-12) primary cortical samples. Human LHX6 cINs displayed a broader spatial distribution across the cortical wall than their mouse counterparts (Fig. 6K-6M), consistent with the model’s prediction. Together, these findings suggest that species-specific differences in leading-process branching and homotypic interactions may cell-autonomously shape the spatial distribution of migrating cINs during cortical invasion. The organization of tangential streams of migrating cINs in the mouse cortex is also sculpted by gradients of molecular guidance cues. Among these, CXCL12/Cxcl12 acts as a key chemoattractant, regulating the positioning of cINs within migratory streams and their subsequent invasion of the cortical plate (*30, 40*). We therefore compared CXCL12/Cxcl12 expression in the E13.5 mouse and GW11-12 human cortex. The overall pattern of CXCL12/Cxcl12 expression was comparable between species (Fig. S6N-S6O). However, whereas mouse LHX6 cINs were organized into distinct MZ and SVZ/IZ migratory streams, human LHX6 cINs displayed a broader distribution that extended beyond both domains. These findings suggest that the broader distribution of human migrating cINs is unlikely to reflect differences in CXCL12/Cxcl12 expression, supporting a role for cell-cell interactions in shaping species-specific cIN distribution during cortical invasion.

Our computational model predicts that increased branching and homotypic repulsion in mouse cINs would impair their cortical invasion by disrupting the organization of the main migratory streams (Fig. S6I). To test this prediction, we performed in utero electroporation (IUE) of Cre-dependent CCP1^WT^ or CCP1^Dead^ expression constructs (Fig. S6J-S6L) into the MGE of Dlx5/6:Cre-GFP E13.5 mouse embryos and collected their brains at E17.5. Analysis of the laminar distribution of transduced cINs across the cortical wall revealed a shift in their distribution with greater dispersion between the main migratory streams in CCP1^WT^- expressing cINs, but not in CCP1^Dead^-expressing cINs (Fig. 6N-6Q). Laminar distributions of corresponding non-transduced GFP^+^ cINs were not affected (Fig. S6M). Most CCP1^Dead^-transduced cINs entered the SVZ/IZ stream, as expected for a cohort of cINs born at mid-corticogenesis (*41*). These findings suggest that increasing branching and homotypic repulsion by raising CCP1 expression partially shifts cortical invasion towards the human cIN laminar distribution pattern.

We next asked whether mouse-human differences in cIN behavior and cortical dispersion extend to ferret, a gyrencephalic species with cortical developmental features intermediate between rodents and primates (*42, 43*). E34-E36 ferret MGE-derived cINs exhibited intermediate migration speed, branching complexity, and homotypic contact frequency relative to mouse and human cINs (Fig. S8A-S8H; Movie S8). Repulsive contacts were likewise intermediate, accounting for 44.4% of ferret contacts versus 18.5% in mouse and 66.7% in human cINs (Fig. S8I). Consistent with these cellular phenotypes, the tangential distribution of LHX6 cINs across the cortical wall in ferret was also intermediate between observations in mouse and human (Fig. S8J-S8K). At the molecular level, E36 ferret MGE extracts showed higher CCP1 levels than age-matched mouse E13.5 extracts, accompanied by reduced DCX expression (Fig. S8L-S8O). Together, these findings suggest that interspecies variation in CCP1 levels may contribute to differences in cIN branching and homotypic interactions which, ultimately, influences how cINs distribute across the developing cortical wall.

## Discussion

Here, we present an evo-devo comparison of the tangential migration of ventrally derived cINs in the developing cerebral cortex of different mammalian species. While these cells exhibit a conserved mode of migration, characterized by alternating phases of nuclear translocation and dynamic branching of the leading process, we identify species-specific differences in cIN migratory dynamics and homotypic contacts that shape their tangential trajectories and spatial organization within the cortical wall. One notable difference is that human cINs migrate slower than their mouse counterparts, despite the greater distance that they must travel to reach the cortex in humans. This difference is associated with an increased basal level of nuclear pausing prior to nucleokinesis in human cINs. Further, we found that this process is sensitive to CCP1 expression level, which regulates actomyosin contractility posterior to the nucleus and thereby influence nuclear translocation. In addition, evolutionary changes in CCP1 expression in cINs modulate the recruitment of DCX to MTs by altering their glutamylation profile, thus influencing leading-process branching dynamics via MT stabilization. DCX is a MAP that plays a critical role in neuronal migration during cortical development. Pathogenic variants in DCX disrupt this process and are associated with distinct cortical malformations, typically manifesting as classic lissencephaly in males and subcortical band heterotopia (SBH) in females because of projection neuron (PN) migration defects (*44, 45*). Beyond its established role in the migration of PNs, DCX also contributes to the tangential migration of cINs. Experimental studies have shown that DCX deficiency disrupts cIN branching and nucleokinesis and can delay migration (*24, 46*).

The enhanced branching activity observed in human cINs correlates with lower level of glutamylated MTs and is accompanied by more frequent repulsive contacts between neighboring cINs. This contributes to the dispersed pattern of tangential migration for human cINs as compared with that seen in mouse and ferret. The evolutionary changes in CCP1 expression observed in cINs across species concurrently modulate both nuclear pausing duration and leading-process branching activity during cIN migration. This finding suggests that CCP1 may act as a molecular coordinator that fine-tunes the balance between these two key migratory processes, thereby coordinating the dynamic interplay between leading-process branching and nuclear translocation during cIN migration. This hypothesis is further supported by gain-of-function experiments, in which the artificial elevation of CCP1 expression in mouse cINs modulates their migratory behavior, suggesting that CCP1 levels influence the regulation and coordination of these cellular dynamics. Specifically, elevating CCP1 expression in mouse cINs increased the duration of nuclear pausing, enhanced leading-process branching dynamics, and increased the frequency of repulsive interactions between neighboring cINs. The cINs transduced at E13.5 with inactive CCP1 (CCP1^dead^) were predominantly detected migrating through the lower SVZ/IZ stream at E17.5, consistent with the expected migratory pattern of cINs born during mid-corticogenesis (*41*). Such stream enrichment may reflect intrinsic differences in migratory behavior among cINs that are born at different developmental stages (*47, 48*). Strikingly, expression of active CCP1^wt^ in E13.5 cINs redirected their tangential migration away from cortical migratory streams at E17.5. These changes indicate that increased CCP1 expression can modulate multiple aspects of cIN migratory behavior, partially recapitulating the cellular dynamics observed in human cINs. However, the effects of CCP1 overexpression in mouse cINs are partial, with the induced phenotypes not fully recapitulating those observed in human cells, indicating that additional molecular mechanisms may contribute to species-specific regulation of cIN migration. In this regard, we measured higher expression levels of the tubulin tyrosine ligases *TTLL1* and *TTLL7* by RT-qPCR, which catalyze tubulin glutamylation, in human MGE extracts compared with mouse counterparts (Fig. S2F-S2G). Notably, TTLL1 and TTLL7 generate distinct patterns of tubulin polyglutamylation that can differentially modulate the activity of specific MT-interacting proteins (*49*). Therefore, differences in TTLL1 and TTLL7 expression may also contribute to the regulation of MT-DCX interactions, thereby reducing DCX affinity for MTs in human cINs as compared to mouse ones. Such an effect could act in concert with the higher CCP1 expression in human cINs further lowering overall MT glutamylation levels and thereby further contributing to species-specific differences in cIN migratory dynamics.

When focusing on branching dynamics, the enhanced homotypic repulsion observed between human cINs may not only reflect the increased formation of dynamic branches along their leading processes but also evolutionary changes in the molecular mechanisms underlying cIN-to-cIN repulsion. To address this, we performed CellChat analysis for mouse and human cINs, with scRNAseq datasets to identify candidate cell-cell interactions that could account for the homotypic repulsion observed between human cINs (Fig. S7D-S7F). We hypothesized that species-specific differences in these interactions might either weaken attractive signaling or enhance repulsive cues. Among the cell-cell interactions shared between mouse and human cINs, we identified a potential homotypic NCAM1-NCAM1 interaction, which appeared to be weaker in human cINs (Fig. S7E). This observation is further supported by proteomic comparison of mouse MGEs (E12. and E13.5) and human forebrain assembloids (hFAs; 90 days in couture) protein extracts, which revealed differences in NCAM1 abundance (data not shown). Strikingly, CellChat analysis also identified SLIT1/2-ROBO1-mediated repulsive signaling as a potential human-specific interaction between migrating cINs (Fig. S7F). In mice, although Slit proteins secreted by the ventromedial forebrain germinal regions promote cIN migration away from these regions (*50*), directional guidance toward the cortical wall relies on Slit1/2-independent repulsion from subcortical regions and attraction to cortical regions (*51*). Together, these findings raise the possibility that SLIT1/2-ROBO1 signaling has acquired an evolutionary role in regulating cIN-cIN interactions in the human cortex, potentially contributing a more dispersed pattern of tangential migration within the cortical wall.

An intriguing aspect of the migration of mouse cINs in organized cortical streams is that it has traditionally been attributed largely to the spatial distribution of extrinsic cues, including amongst others, chemokine gradients such as Cxcl12 and long-range attractive signals mediated by Nrg1 (*52, 53*). The distribution of this chemokine was comparable in the developing cortex of both mouse and human. Instead, our findings point to a substantial cell-intrinsic component underlying the organization of cortical streams and suggest that evolutionary changes in the intrinsic migratory properties of cINs may contribute to shaping the spatial distribution of cINs invading the cortical wall. Our work thus reveals how cell-intrinsic evolutionary divergence in migration behavior can be superimposed on a deeply conserved cellular program, providing a molecular framework for understanding how cIN migration has been adapted across mammalian evolution.

Why has such an evolutionary adaptation in cIN migration emerged? We and others have shown that cell migration in the forebrain not only positions neural cells at their final destinations but also provides a dynamic platform for crosstalk with neighboring cells through direct interactions and/or the release of signaling cues (*20, 29, 54*). Our data suggests that the complexity of cIN migration and the organization of cortical streams correlate with the degree of cortical folding across mammalian species. In the lissencephalic mouse cortex, migrating cINs form sharp, densely packed streams, whereas in the developing human cortex, which begins to fold at around GW17 and subsequently develops higher-order folds (including third order ones), cINs exhibit a more dispersed distribution and less sharply defined cortical streams. The ferret occupies an intermediate position in both cortical folding and in the spatial organization of cIN migration, displaying an intermediate degree of cortical stream dispersion and folding, with up to second-order folds. These observations raise the intriguing possibility that evolutionary changes in the organization of cIN migration may contribute to cortical folding by modulating the spatial distribution of chemical and potentially physical cues, influencing local tissue interactions and promoting tissue symmetry breaking during cortical development. Another possibility is that increased dispersion of migrating cINs, which exert traction forces on the forebrain parenchyma (*55*), may homogenize the spatial distribution of these forces across the cortical plate. Such a redistribution could contribute to a progressive stiffening of the cortical plate relative to the underlying, softer progenitor-rich region (*56*), thereby establishing mechanical heterogeneities that could bias tissue deformation and the emergence of cortical folds. In this view, changes in cIN migration may contribute to cortical folding through a physical mechanism acting in concert with additional biological features such as tissue expansion, cortical wall thickness, and the three-dimensional geometry of the early forebrain, whereby the spatial organization of cell-generated forces could help imprint future folding patterns, converging with classical models of cortical folding based on differential biomechanical properties and buckling (*57*).

## Methods summary

Human, mouse and ferret forebrain tissues were used to compare cIN migration at corresponding stages of cortical development. Mouse embryos were analyzed at embryonic day 13.5 (E13.5), ferret embryos at E36, and human fetal tissue at gestational weeks 11-13 (GW11-13). Human fetal tissue was obtained following voluntary pregnancy termination at CHR Hospital-Site Laveu (Liège, Belgium), with informed consent and approval from the Hospital-University Ethics Committee of Liège (#2021/216). Gestational age was estimated by ultrasound and confirmed by tibial length. Mouse procedures were approved by the relevant Belgian ethics committee (#2816, #2455, #2537) and conducted in accordance with Belgian and European animal protection guidelines. Pregnant ferrets were obtained from Euroferret (Denmark) and housed at the animal facility of Miguel Hernández University (Spain). Ferret procedures were approved by the CSIC Ethics Committee and the Generalitat Valenciana, in accordance with Spanish and EU regulations, under license number 2024-VSC-PEA-0256.

To visualize and track migrating cINs, brain organotypic slices and MGE explants were labeled with a lentiviral Dlx5/6-eGFP reporter. Organotypic slices were prepared from E13.5 mouse and GW11-13 human brains, sectioned at 250µm thickness, maintained on ice and harvested on a Millicell (Millipore) culture insert placed on glass-bottomed dishes. Dishes were filled with 1:1 neurobasal/DMEM-F12 medium supplemented in B27, N2, and glutamine and non-essential aminoacids. Slices were kept in a humidified incubator saturated with 5% CO2 for a maximum of 24 hours before time-lapse recording on a spinning disc fluorescence microscopy. MGE explants were generated by culturing MGEs from mouse embryos (E13.5), ferret embryos (E36), and human fetal brains (GW11-13) on top of a mouse-derived cortical feeder. Briefly, glass-bottomed dishes were coated with poly-ornithine and laminin. The feeder layer of cortical cells was prepared by mechanical dissociation of E13.5 mouse cortices in supplemented neurobasal medium. Dissociated cells were plated in coated glass-bottomed dishes and left to adhere for 2 hours at 37°C in a humidified incubator saturated with 5% CO2. The next day, explants were plated above a cortical feeder layer. cINs migrating from MGE explants were imaged by inverted confocal or spinning-disk microscopy for at least 16 hours, beginning 48 hours after plating for mouse and ferret explants and at least 72 hours after plating for human explants. Time-lapse recordings of both explants and organotypic slices were used to quantify migration speed, nucleokinesis, leading-process dynamics, growth-cone splitting, branching complexity, and the frequency and outcome of homotypic contacts between cINs.

Human pluripotent stem cell (hPSC) lines were used to generate human ventral forebrain organoids (hvFOs) and human forebrain assembloids (hFAs). The panel comprised three patient-derived lines carrying loss-of-function mutations in *AGTPBP1*, which encodes CCP1 (CCP1^Mut#1-#3^) and four controls, including two independent hiPSC lines (CCP1^WT#2-#3^), one hESC line (H9; CCP1^WT#1^), and an isogenic control of CCP1^Mut#2^ (CCP1^Iso#2^). Most experiments were performed using CCP1^WT#2^, CCP1^Mut#2^, and CCP1^Iso#2^, whereas the remaining lines were used to assess phenotype reproducibility. The regional identity of hvFOs was validated by RT-qPCR. To characterize their cellular composition and developmental trajectories, scRNAseq was performed on dissociated hvFOs using the Chromium GEM-X Single Cell 3′ v4 platform. Libraries were processed using Cell Ranger multi (v9.0.1) against the GRCh38 reference genome (release 112), including intronic reads in quantification. Downstream analyses were performed in Seurat (v5) and included quality control, normalization, dimensionality reduction, data integration, clustering, and cell-type annotation. Cells were filtered based on detected features, unique molecular identifiers (UMIs), and mitochondrial and ribosomal transcript content. The resulting datasets contained the major cell populations expected in hvFOs, including progenitors, dividing progenitors, oligodendrocyte precursor cells (OPCs), maturing interneurons, cortical-like interneurons (clINs), hindbrain-like interneurons (hilINs), and hypothalamic-like interneurons (hylINs) and a pool of mature clINs. Broadly comparable developmental trajectories were observed across the analyzed lines, providing a framework for assessing CCP1-associated phenotypes while accounting for genetic background and cellular composition. To investigate the effects of CCP1 loss of function on cIN migration, time-lapse imaging was performed on organotypic slices of hFAs and explants of hvFOs cultured on cortical feeder-cell layers as described above. For 90-day-old hFA imaging, they were embedded in low-melting-point agarose and sectioned coronally at 250 µm using a vibratome. Slices were maintained on membrane culture inserts and imaged by confocal microscopy at 15 minutes intervals for at least 16 hours. For hvFO explant imaging, organoids were dissected, labeled with lentiviral reporters, and cultured on cortical feeder-cell layers before imaging. Migratory behavior was quantified using the same general analytical framework applied to cross-species MGE explants, including migration dynamics, nucleokinesis, leading-process extension and branching, growth-cone splitting, and homotypic interactions between cINs.

Immunohistochemical analyses were performed on mouse (E13.5), ferret (E36), and human (GW10-13) forebrains to analyze CCP1 expression, and to compare cIN distribution across the developing cortical wall and examine the spatial expression of guidance cues. Brains were fixed overnight in 4% paraformaldehyde at 4°C, cryoprotected overnight in 30% sucrose, embedded in OCT, and cryosectioned at 20 or 65 µm. Sections were permeabilized and blocked in PBS containing 0.3% Triton X-100 and 10% donkey serum for 1 hour at room temperature, incubated with primary antibodies overnight at 4°C, and subsequently labeled with Alexa Fluor-conjugated secondary antibodies and DAPI. Images were acquired using a Zeiss LSM980 confocal microscope or a slide scanner with 20×, 40×, or 63× objectives. LHX6-positive cINs were quantified across bins spanning the cortical wall to compare their relative distribution across cortical depth between species. Immunolabeling for guidance cues was used to assess their spatial expression patterns in the developing cortex. To assess the effects of increased CCP1 activity on cIN distribution in vivo, Cre-inducible constructs encoding wild-type CCP1 (CCP1^wt^) or catalytically inactive CCP1 (CCP1^dead^) were transduced by IUE into the MGE of E13.5 Dlx5/6:Cre-GFP mouse embryos. Brains were collected at E17.5 and sectioned at 100 µm using a vibratome. Sections were processed for immunohistochemistry and imaged to quantify the distribution of transduced cINs across the cortical wall. The distribution of labeled cells was analyzed using a bin-based approach and compared between CCP1^wt^ and CCP1^dead^ expressing groups to determine whether increased CCP1 activity altered the pattern of cortical invasion.

A two-dimensional numerical model of cIN migration was developed to investigate how branching dynamics and homotypic repulsion influence population-level cortical invasion (DOI: 10.5281/zenodo.22869350). Individual cells were represented as dynamically branching structures composed of nodes connected by harmonic oscillators, initially representing the trailing process, soma, and leading process. Branches could be added or removed probabilistically, while cells moved through the simulated environment by stochastic node displacements. The model included a cortical geometry extending from the pallium to the cortical plate, with boundaries constraining cell movement, and a predefined guidance-cue field that modulated migration probability. Cortical geometry and the guidance-cue field were kept constant across simulated conditions. Homotypic repulsion was modeled as a force between leading nodes and neighboring branches that decayed with distance according to an inverse-cube relationship. The proportion of cells assigned repulsive interactions was varied to represent mouse (2%), human (100%), and mouse CCP1 gain-of-function (50%) conditions. Simulations were used to assess how differences in branching and repulsive interactions affected the organization of migratory streams and the distribution of cINs across the cortical wall.

## Acknowledgements

The authors thank the following technological GIGA platforms: In vitro imaging (Alexandre Hego and Gaëtan Lefevre, Raafat Stephan, and Sandra Ormenese), Viral Vector (Emmanuel Di Valentin), Bioformatics (Arnaud Lavergne), Genomics (Wouter Coppieters and Latifa Karim), and Proteomics for their technical support (Gabriel Mazzucchelli and Christopher Kune). Some schemes were designed with BioRender (https://BioRender.com/ipj6w6t). We thank Gabriele Lignani for sharing the pLv-Dlx5/6-eGFP construct, Ira Espuny-Camacho for hiPS culture training, Olivier Peulen and Raphaël Pfeiffer for guidance with metabolic analyses. We thank Véronique Henriot for cloning the shRNA constructs that target CCP1. Note on AI tools: ChatGPT (GPT-6) was used for grammar correction, while Claude (Opus 5.5 and Sonnet 5.5) was used to review the scripts.

## Author contribution

L.N. and A.Bonafina conceptualized the study and experiments. A.Bonafina performed most experiments and analyzed data. A.Boutsen designed and performed experiments involving human induced pluripotent stem cells (hiPSCs), performed mitochondrial time-lapse imaging and analysis, and conducted bioinformatic analyses. H.Y.C. designed, performed, and analyzed the histological and in utero electroporation experiments. M.J.-T. designed and performed biochemical experiments and analyzed the results. B.C. maintained hiPSCs in culture and performed and analyzed biochemical experiments. E.E. contributed to the preparation, handling, and labeling of human brain slices. A.C. and A.Bonafina prepared samples and performed time-lapse imaging of ferret cINs. A.C. and V.B. provided ferret samples as well as feedback and guidance on the ferret cIN experiments. L.N., R.C., S.P.H., and A.F. contributed to the preparation of the ethics application for human tissue collection and to the collection and handling of human tissue samples. H.B. and R.I. analyzed mitochondrial experiments and designed and validated some of the vectors used in this study.

N.R. performed RT-qPCR analyses of human and mouse MGE samples and proximity ligation assay (PLA) analyses. S.T. performed fluorescence-activated cell sorting for proteomic analysis. C.K. and G.M. performed and processed the proteomic experiments.

I.G.L. and P.V. contributed to experiments assessing mitochondrial activity, with P.V. providing guidance. I.G.L. performed and analyzed oxygraphy measurements in organoids.

R.L.B. and S.C. contributed to the conceptualization and planning of the assembloid protocol.

C.J. provided constructs for CCP1 overexpression and expression of the catalytically inactive CCP1 mutant and provided insights into CCP1 biology. B.L. designed, performed, and analyzed biochemical and molecular biology experiments. J.-Y.R. conceptualized and designed the computational model of surrogate cells invading the cortex. A.Bonafina and L.N. wrote the manuscript, with input from all authors.

## Data and materials availability

All annotated data are available at the ProteomeXchange Consortium via the PRIDE (accession number) other data are in the main paper or the supplement.

## List of Supplementary Materials (to be updated)

### Materials and Methods

Table S1 – S9

Fig S1 – S8

Movie S1-S8

---

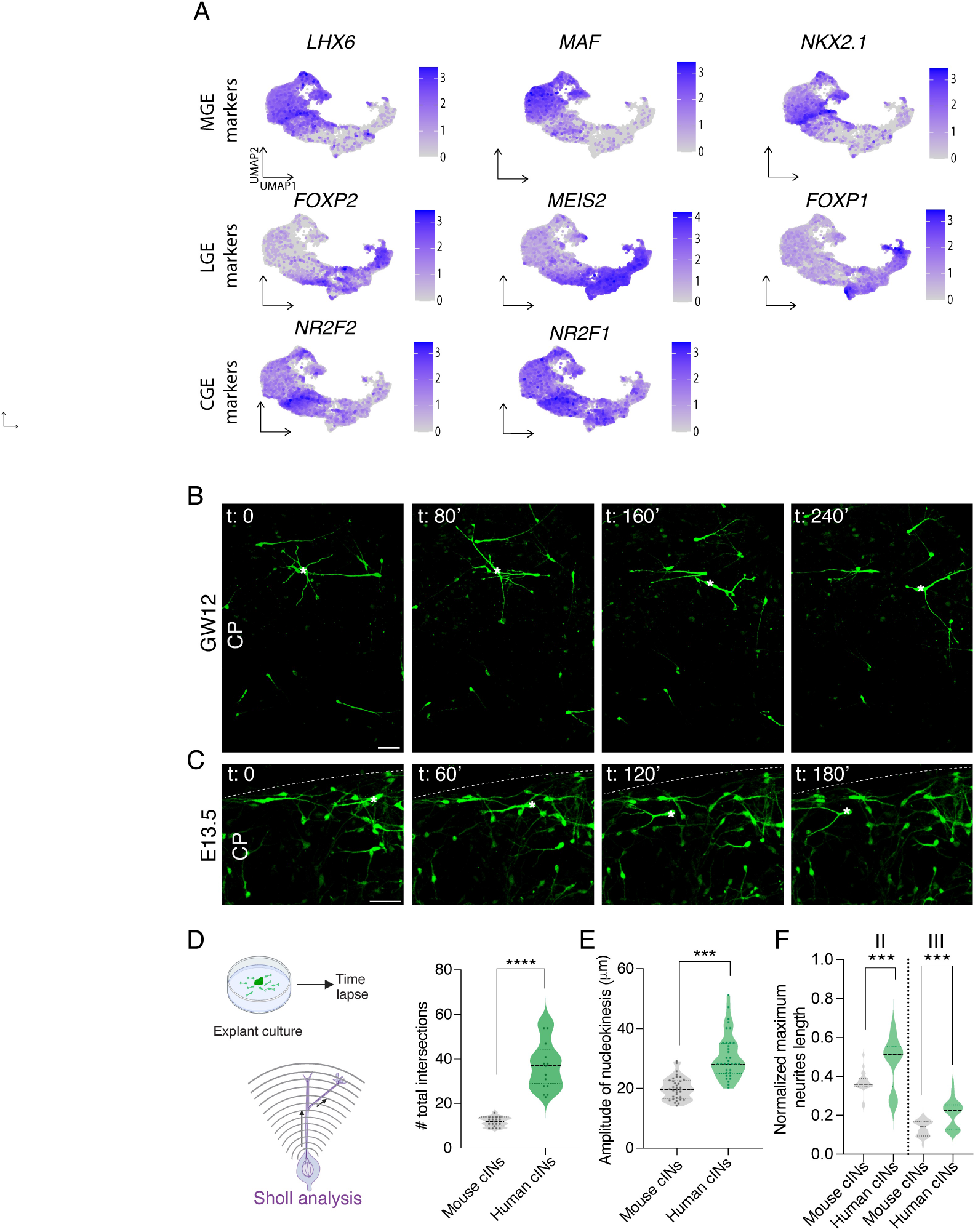

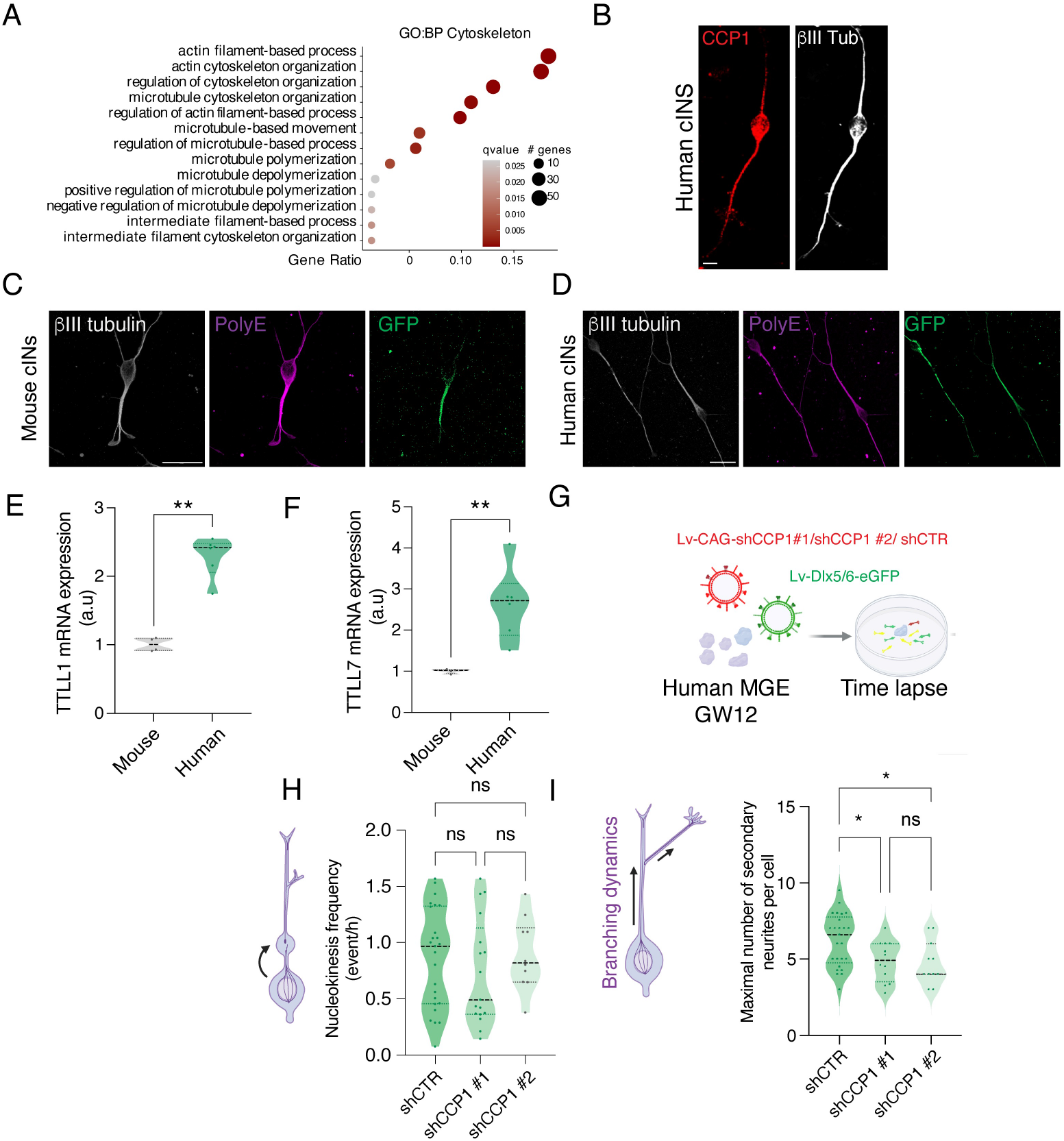

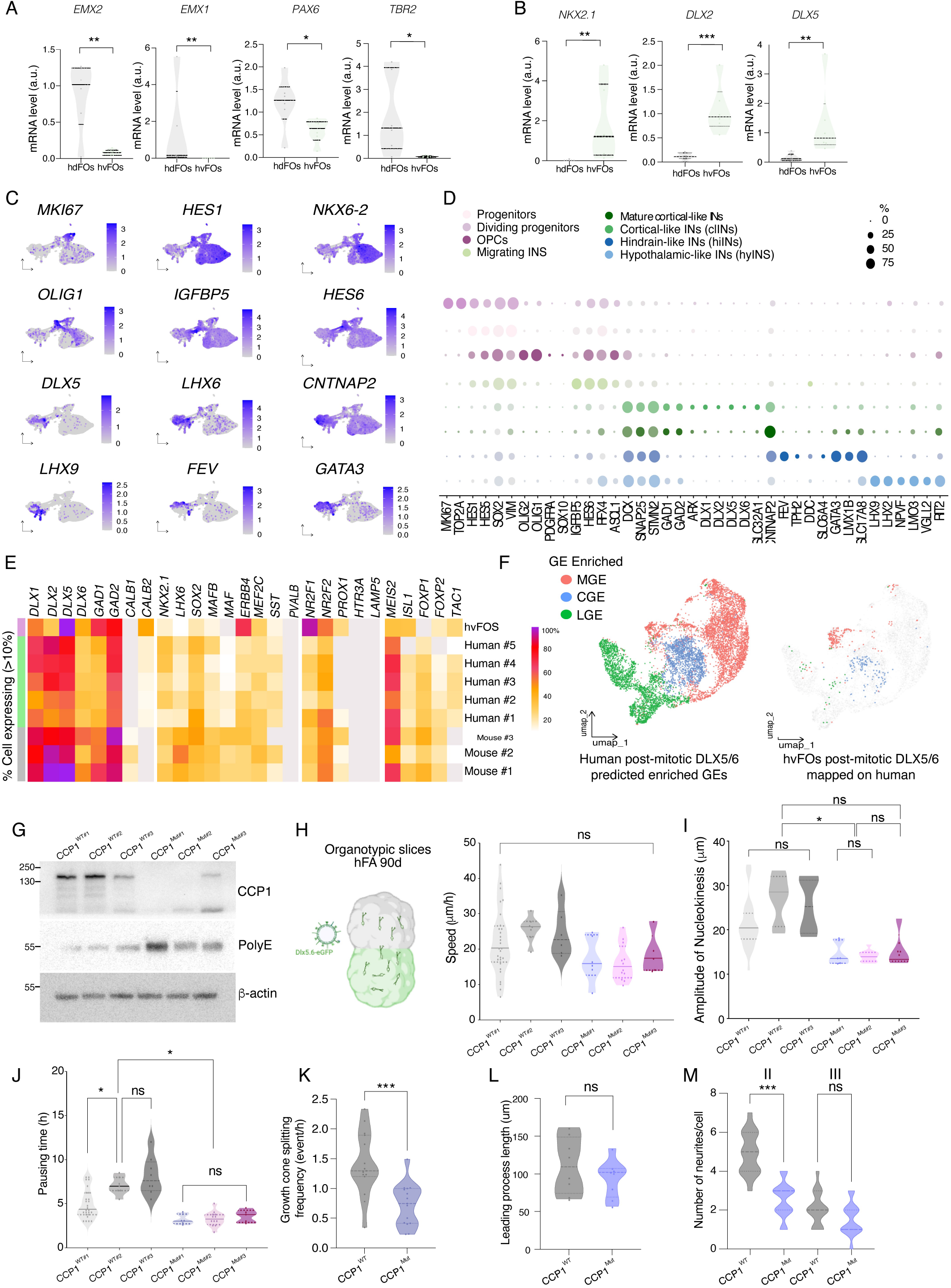

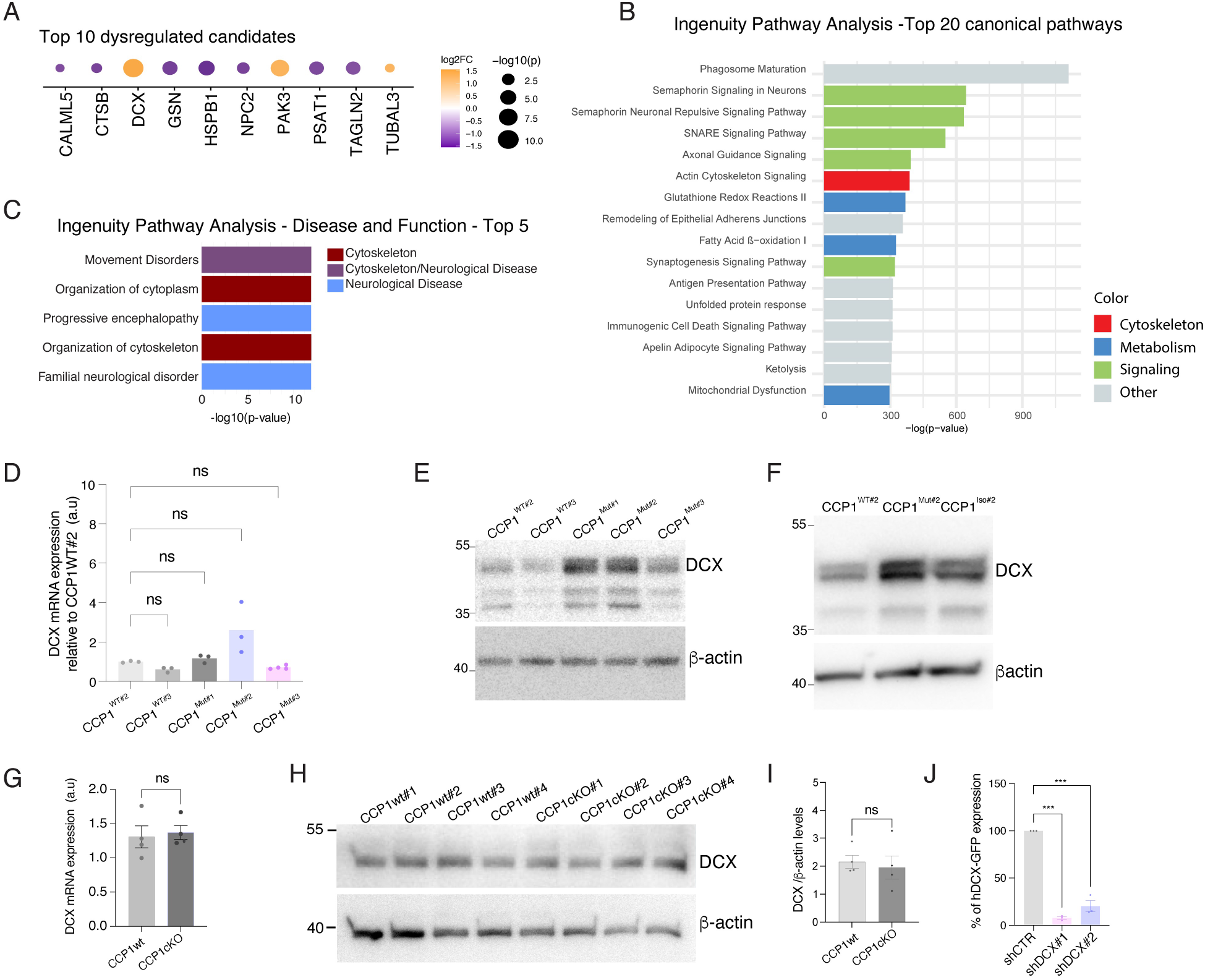

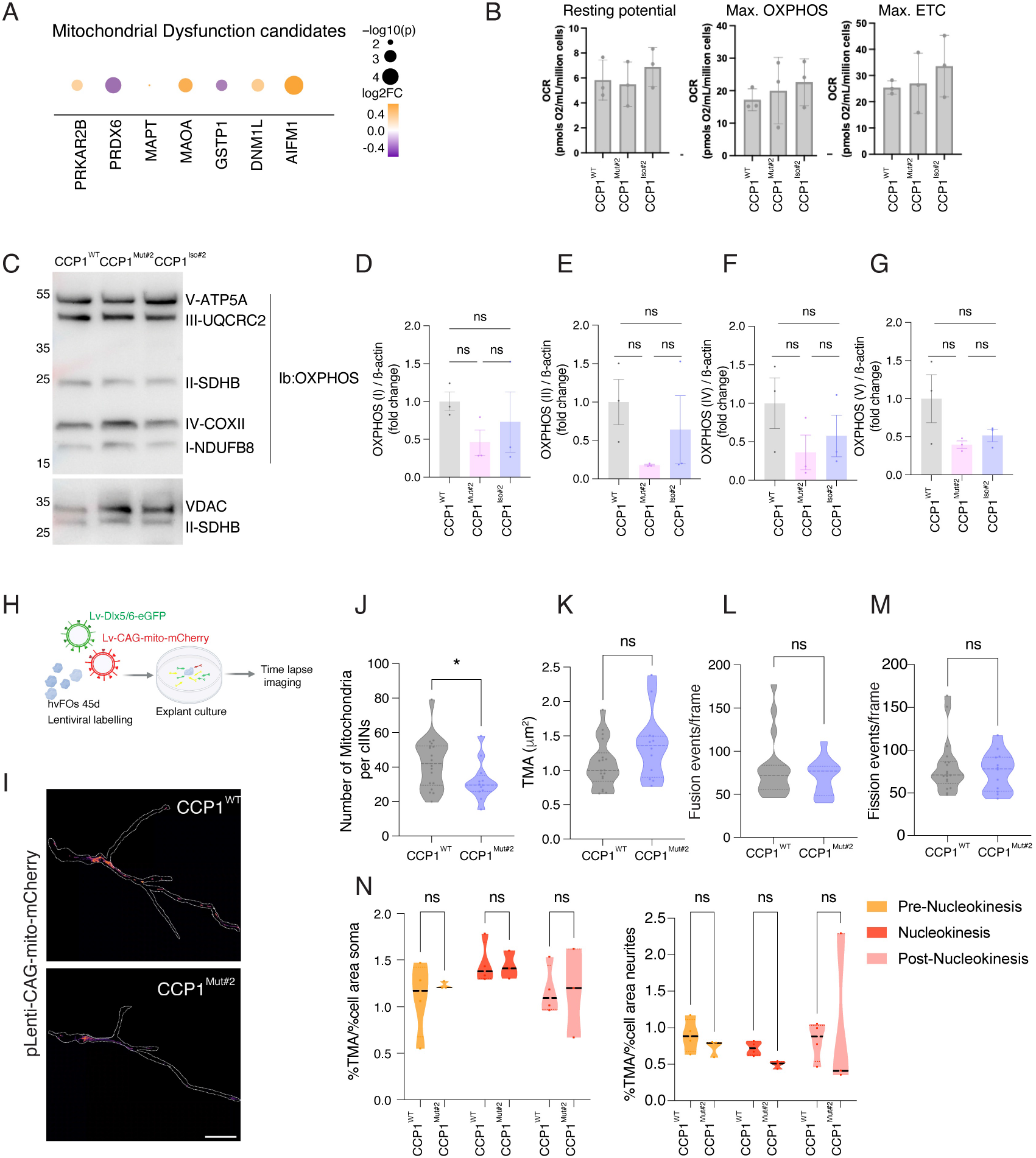

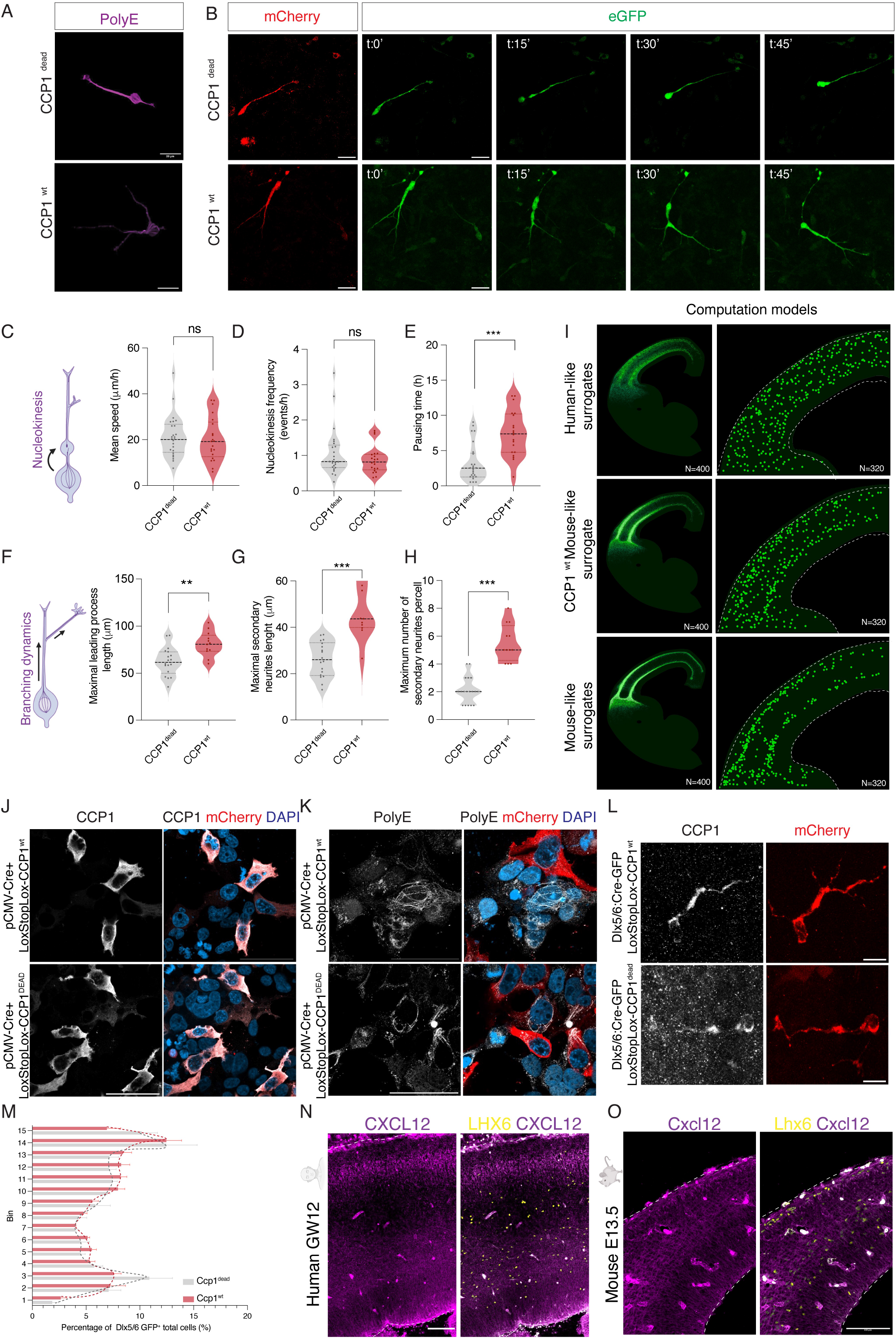

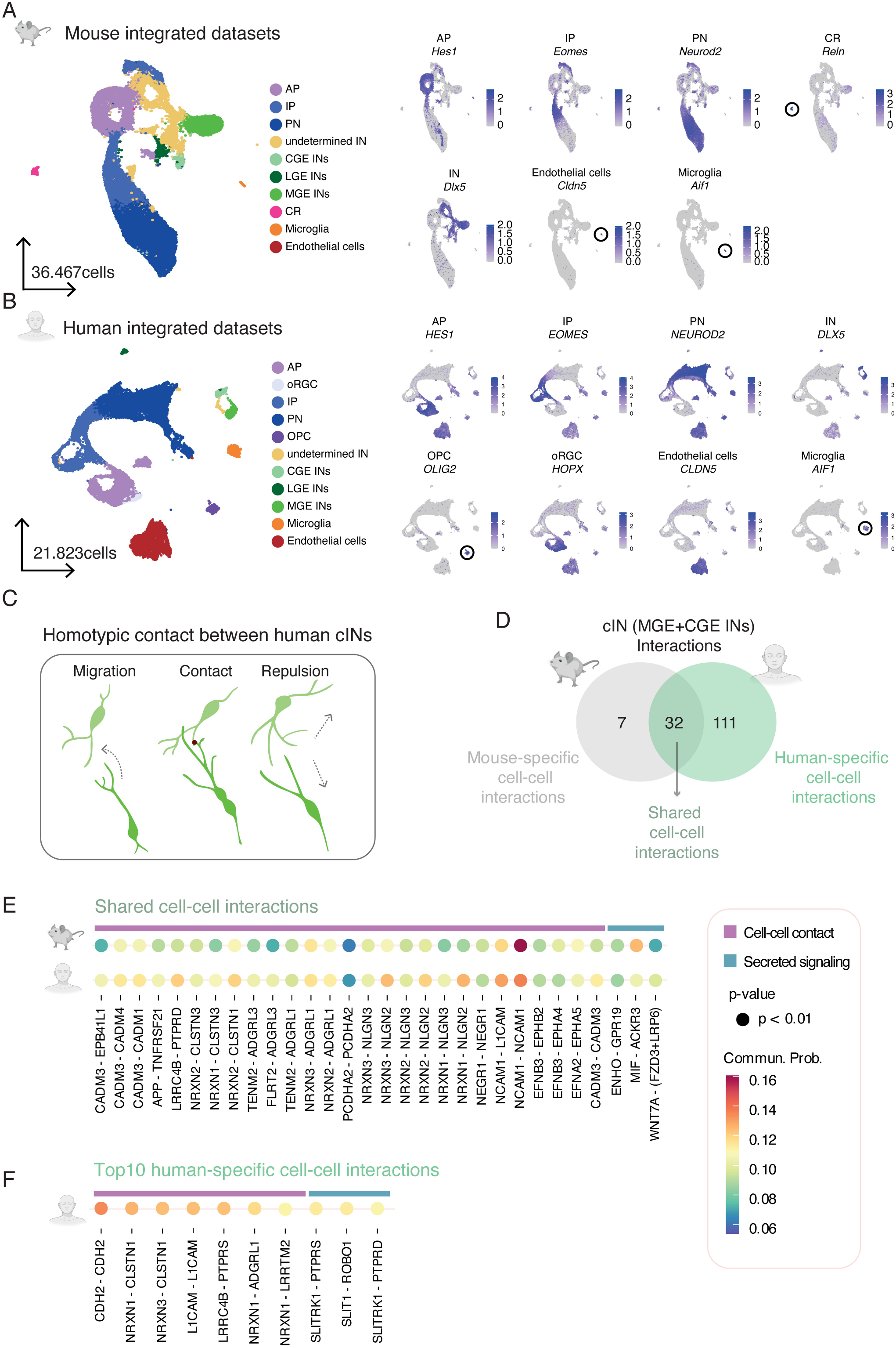

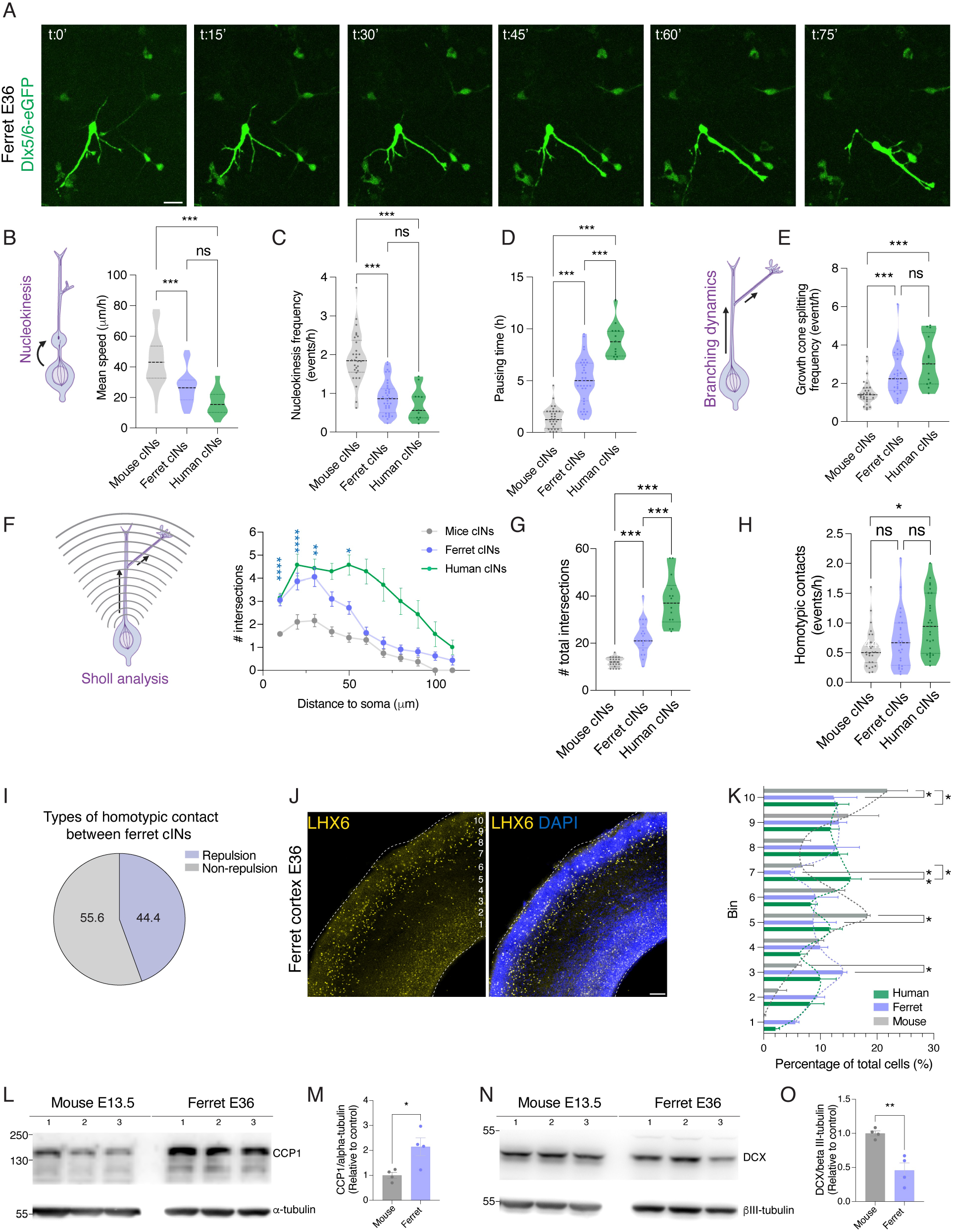

