## Supplemental information for "Evolution Shapes Cortical Interneuron Migration"

**Material & Methods**

KEY RESOURCES TABLE

| **REAGENT or RESOURCE** | **SOURCE** | **IDENTIFIER** |
| --- | --- | --- |
| ***Antibodies*** | | |
| Polyclonal rabbit anti-CCP1 | Proteintech | 14067-1-AP, RRID: AB_2225559 |
| Monoclonal mouse anti-BIII Tubulin (Tuj1 clone) | BioTechne | MAB1195, RRIB: AB_357520 |
| Monoclonal rabbit anti-βIII-tubulin | Cell Signaling Technology | 5568: RRID: AB_10694505 |
| Monoclonal mouse anti-LHX6 | Santa-Cruz | Sc-271433, RRID: AB_10649856 |
| Polyclonal rabbit anti-PolyE | Adipogen | AG-25B-0030, RRID: AB_2490540 |
| Monoclonal rabbit anti Alpha-tubulin (HRP conjugate) | Cell Signaling | 9099, RRID: AB_10695471 |
| Polyclonal goat anti-GFP | Abcam | Ab6673, RRID: AB_305643 |
| Monoclonal rat anti-GFP | Nacalai Tesque | 04404-26, RRID: AB_10013361 |
| Monoclonal mouse anti-DCX | Santa-Cruz | Sc-271390, RRID: AB_10610966 |
| Monoclonal mouse anti beta-Actin (HRP conjugate) | Cell Signaling | 12262, RRID: AB_2566811 |
| Monoclonal mouse anti-OXPHOS | Abcam | Ab110411, RRID: AB_2756818 |
| Polyclonal rabbit anti-VDAC | Abcam | Ab15895, RRID: AB_2214787 |
| Polyclonal goat anti-mouse igG (Fc specific) | Sigma | M4280-1mL, RRID: AB_260521 |
| Polyclonal goat anti-mCherry | Biorbyt | orb11618, RRID: AB_2687829 |
| Monoclonal mouse anti-CXCL12 | R&D | MAB310, RRID: AB_2276927 |
| ***Chemicals, Peptides, and Recombinant Proteins*** | | |
| Neurobasal Medium | Gibco, Thermo Fisher Scientific | 21103049 |
| B-27 Supplement | Gibco, Thermo Fisher Scientific | 17504044 |
| N-2 Supplement | Gibco, Thermo Fisher Scientific | 17502048 |
| Penicillin/Streptomycin | Gibco, Thermo Fisher Scientific | 15140122 |
| DMEM/F-12, GlutaMAX™ supplement | Gibco, Thermo Fisher Scientific | 31331028 |
| KnockOut™ Serum Replacement | Gibco, Thermo Fisher Scientific | 10828028 |
| GlutaMAX™ Supplement | Gibco, Thermo Fisher Scientific | 35050-038 |
| MEM Non-Essential Amino Acids Solution | Gibco™ | 11140035 |
| Beta mercapto ethanol | Sigma-Aldrich | M7154 |
| Heparin | Sigma-Aldrich | H3149 |
| LDN193189 | Miltenyi Biotech | 130-103-925 |
| B27 -VitA | Gibco, Thermo Fisher Scientific | 12587010 |
| Insulin | Sigma-Aldrich | I9278 |
| SAG | Sigma-Aldrich | SML314 |
| IWP2 | Sigma-Aldrich | 10536 |
| SB431542 | Sigma-Aldrich | S4317 |
| XAV939 | StemCell | 72-672 |
| Poly-D-lysine hydrobromide | Sigma | P0899-100MG |
| Laminin | Sigma-Aldrich | L2020 |
| Recombinant Mouse N-Cadherin Fc  Chimera Protein, CF | R&D Systems | 6626-NC-050 |
| Fibronectin | Sigma | F4759-1MG |
| Basic Fibroblast Growth Factor (bFGF) | Gibco, Thermo Fisher Scientific | 100-18B |
| ROCK inhibitor (Y-27632) | BioConnect | S1049 |
| Accutase | Merck-Sigma | SCR005 |
| Dulbecco’s Phosphate Buffered Saline (DPBS) | Gibco, Thermo Fisher Scientific | 14040133 |
| Antigen Retrieval Solution | Agilent-Dako | S1699 |
| BSA | Roth | 8076.3 |
| Donkey Serum | Jackson ImmunoResearch | 017-000-121 |
| Triton X-100 | Sigma-Aldrich | T8787 |
| Tween-20 | Sigma-Aldrich | P9416 |
| DAPI | Sigma-Aldrich | D9542 |
| Mounting medium | Agilent-Dako | S3023 |
| Protease inhibitor | Sigma-Aldrich | P8340 |
| Phosphatase inhibitor | Sigma-Aldrich | 4906845001 |
| PageRuler Plus Prestained Protein Ladder | Gibco, Thermo Fisher Scientific | 26619 |
| BSA Fraction V | Gibco, Thermo Fisher Scientific | 15260-037 |
| FluorSave Reagent | Merck-Sigma | 345789 |
| Neg-50 Frozen Section Media | Epredia | 6506 |
| Sucrose | VWR Chemicals | 27478.365 |
| ***Critical Commercial Assays*** | | |
| Quick-RNA Miniprep | Zymo Research | R1054 |
| Pierce ECL Western Blotting Substrate | Gibco, Thermo Fisher Scientific | 32106 |
| Papain Dissociation System | Worthington | LK003160 |
| RevertAid H Minus First Strand cDNA Synthesis Kit | Thermo Fisher Scientific | K1632 |
| TotalSeq™-B Hashtag Antibodies 8, 9, 10 | BioLegend | B0258, B0259, B0260 |
| Chromium GEM-X Single Cell 3' v4 kit | 10x Genomics | PN-1000690 |
| Duolink® In Situ PLA® Probe Anti-Mouse MINUS | Sigma-Aldrich (Duolink) | #DUO92004 |
| Duolink® In Situ PLA® Probe Anti-Rabbit PLUS | Sigma-Aldrich (Duolink) | #DUO92002 |
| Duolink® In Situ Detection Reagents Red | Sigma-Aldrich (Duolink) | #DUO92008 |
| Duolink® In Situ Wash Buffers, Fluorescence | Sigma-Aldrich (Duolink) | #DUO82049 |
| Pierce BCA Protein, Assay Kit | Gibco, Thermo Fisher Scientific | 23225 |
| GoTaq Master Mix | Promega | A6002 |
| ***Deposited data*** |  |  |
| Human GW11, GW12, subpallium scRNA-seq | (*1*) | GEO: GSE165388 |
| Human GW12_01, GW12_02, GW13, Ganglionic eminences scRNA-seq | (*2*) | GEO: GSE135827 |
| Mouse E13.5, CGE LHX6neg/MGE LHX6pos scRNA-seq | (*3*) | GEO: GSE104156 |
| Mouse E13, LGE/MGE/CGE scRNA-seq | (*4*) | GEO: GSE188528 |
| Mouse E14.5, LGE/MGE/CGE scRNA-seq | (*5*) | GEO: GSE190593 |
| hvFOs scRNA-seq | This paper | GEO: to be updated when accepted |
| Human GW13, Telencephalon #1, #2 scRNA-seq | (*6*) | GEO: GSE310125 |
| Human GW13, Cortex scRNA-seq | (*7*) | EGA EGAS00001004107 |
| Mouse E14 Cortex + GE Emx1Cre N=1, N=2, N=3 scRNA-seq | (*8*) | GEO: GSE152281 |
| Mouse E14.5 somatosensory cortex rep1, rep2 scRNA-seq | (*9*) | GEO: GSE155677 |
| Mouse E14.5 cortex WT1, WT2, WT3, WT4, WT5, WT6 scRNA-seq | (*10*) | GEO: GSE123335 |
| Mouse E14.5 Neocortex rep1, rep2 scRNA-seq | (*11*) | GEO: GSE161690 |
| Mouse E13.5 Neocortex scRNA-seq | (*12*) | GEO: GSE107122 |
| Mouse E13.5, E14.5 somatosensory cortex scRNA-seq | (*13*) | GEO: [GSE153162](https://www.ncbi.nlm.nih.gov/gds/?term=GSE153162%5bAccession%5d) |
| ***Experimental Models:*** ***Cell lines*** |  |  |
| hESC H9 WA09 | WiCell | CVCL_9773 |
| hiPSC GM23338*D | Coriell Institute | CVCL_F182 |
| hiPSC GM23340*D | Coriell Institute | CVCL_F184 |
| Patient hiPSC lines (CCP1 LOF, n=3) | Derived in-house; fibroblasts given by Carsten Janke (Curie Institute) | No public identifier. |
| HEK293T cells | ATCC (CRL-1573) | RRID: CVCL_0063 |
| ***Experimental Models:*** ***Organisms/Strains*** |  |  |
| Mouse Dlx5,6:Cre-GFP *RjOrl*:Swiss (outbred) | The Jackson laboratory | 023724 |
| Mouse CCP1fl/fl | (*14*) |  |
| Ferret (Mustela putorius furo) | Provided by Victor Borrell Laboratory |  |
| Human fetal brain tissue (GW10-13) | St Rosalie Hospital, Liège | Ethics approval n° 2021/216 |
| ***Oligonucleotides*** | | |
| Primers for qPCR, see Table S3 | This paper | N/A |
| ***Recombinant DNA*** | | |
| pLvEF1α-dTomato-H1-shRNA ***(***see table S4 for shRNA sequences) | Biosettia | SORT-B11 |
| pLv-CAG-mCherry-mCCP1(WT) | Provided by Carsten Janke | N/A |
| pLv-CAG-mCherry-mCCP1 (dead) | Provided by Carsten Janke | N/A |
| pLv-CAG-Lox-Stop-Lox-mCherry-mCCP1(WT) | Provided by Carsten Janke + Addgene | Addgene 51269 |
| pLv-CAG-Lox-Stop-Lox-mCherry-mCCP1(Dead) | Provided by Carsten Janke + Addgene | Addgene 51269 |
| pLv-CAG-Mito-mCherry | Provided by Pierre Vanderhaeghen | N/A |
| pLv-Dlx5/6-eGFP | Provided by Gabriele Lignani | N/A |
| pCMV-EGFP-hDCX | Addgene | Addgene 32852 |
| pLv-Dlx5/6-mCherry-mCCP1 | Provided by Gabriele Lignani + Carsten Janke | N/A |
| pLenti.PGK.LifeAct-Ruby.W | Vector Builder | Addgene: 51009 |
| ***Software and Algorithms*** |  |  |
| Fiji - Image J | (*15*) | https://imagej.nih.gov/ij/ |
| mTrackJ - Image J | (*16*) | https://imagescience.org/meijering/software/mtrackj/ |
| GraphPad Prism 9 | GraphPad Software | https://www.graphpad.com |
| R 4.5.0 & 4.5.2 | R version | R Core Team (2025). _R: A Language and Environment for Statistical Computing_. R Foundation for Statistical Computing, Vienna, Austria.  <<https://www.R-project.org/>>. |
| QSeurat 5.0.0 & 5.4.0 | Seurat version | (*17*) |
| Python v3.13 | Python version | [https://www.python.org/](https://www.python.org/?utm_source=chatgpt.com) |
| PyCharm 2025.1.1.1 | PyCharm Software | [https://www.jetbrains.com/pycharm/](https://www.jetbrains.com/pycharm/?utm_source=chatgpt.com) |
| QuPath-0.7.0 |  | https://qupath.github.io/ |
| CellChat v2.2.9001 | CellChat R Package | (*18*) |
| ***Other*** |  |  |
| Vibratome Leica VT1000S/VT1200 | Leica Microsystems | VT1000S/VT1200 |
| MatTek glass-bottom 35mm dishes | MatTek Corporation | P35G-0-20-C |
| CELLview^TM^ 4 compartments, glass-bottom 35x10mm dishes | Greiner Bio-One | 627871 |
| FluoroDishes glass-bottom 35mm/23mm | WPI | FD35-100 |
| FemtoJet microinjector | Eppendorf | 5252000013 |
| Tweezer Platinum electrodes | Nepagene | CUY650P3 |
| Electroporator | BTX | ECM 830 |
| Nikon A1R confocal microscope | Nikon | N/A |
| Zeiss LSM 880 confocal microscope | Zeiss | N/A |
| Zeiss LSM 980 confocal microscope | Zeiss | N/A |
| LightCycler 480 Instrument |  |  |
| Amersham ImageQuant 800 imaging system | Cytiva | N/A |
| Epredia CryoStar NX70 cryostat | Epredia | N/A |
| SuperFrost Plus adhesive slides Adhesion Microscope Slides | Epredia | J1800AMNZ |
| EMS-Core Sampling Tool, 0.35mm | Electron Microscopy Sciences | 69039-03 |
| µ-Slide 8 Well | Ibidi | 80826 |
| Nikon Spinning Disc microscope | Nikon | N/A |
| Lipofectamine 2000 | Thermo Fisher Scientific | 11668-019 |
| Ml7 hydrochloride | Tocris | 110448-33-4/4310 |
| Axioscan 7 Slide Scanner | Zeiss | N/A |

CONTACT FOR REAGENT AND RESOURCE SHARING

EXPERIMENTAL MODEL AND SUBJECT DETAILS

***Mouse genetics.*** Male and female CCP1fl/fl (*14*) and Dlx5,6:Cre-GFP (*19*) mice were used following Belgian and European animal protection guidelines 86/609/EEC on animal protection. All procedures were ethics-approved (#2816, #2455, #2537) and supervised by authorized staff. Animals were bred in-house and maintained at 19-22 °C, 40-50% humidity, on a reversed 12:12 light/dark cycle (lights on at 23:00), with ad libitum food and water. Wild-type pregnant SWISS females were purchased from Janvier Labs (France) at specified plug dates and kept in the Animal Facility until the specified date of use.

***Ferrets.*** Pregnant pigmented ferrets (Mustela putorius furo) were obtained from Euroferret (Denmark) and kept on a 16h:18h light:dark cycle at the Animal Facility of the Universidad Miguel Hernández (SEA). Ferret procedures were approved by the CSIC Ethics Committee and the Generalitat Valenciana, in accordance with Spanish and EU regulations, under license number 2024-VSC-PEA-0256.

***Human fetuses*.** Human fetal brain tissue was obtained from voluntary pregnancy terminations at CHR Hospital-Site Laveu (Liège, Belgium) with signed informed consent, following GDPR, the Declaration of Helsinki, and Good Clinical Practice. Ethical approval was granted by the Hospital-University Ethical committee of Liège to collect human fetal brain tissue (n° 2021/216). Gestational age (GW10-13) was estimated by ultrasound and confirmed by tibial length.

***hESCs/hiPSCs.*** Human pluripotent stem cell lines (hPSCs) used in this study were validated using standardized methods and were tested for and maintained mycoplasma free. Multiple control hPSC cell lines were utilized: H9 hESC (WiCell, WA09, metadata: female, no disease associated), and hiPSCs GM23338*D/ GM 23340*D (Coriell Institute, male, no disease associated). A total of three patient cell lines were obtained from the Carsten Janke (Curie Institute, Orsay) and derived in the host laboratory from fibroblasts collected from three subjects (two males and one female), carrying loss of function mutation for *AGTPBP* gene (coding for CCP1) (See **Table S1**). The patient line mutant #2 was also CRISPR genome edited (Synthego) to generate its isogenic control. Ethical approval was granted by the Hospital-University Ethical committee of Liège to collect patient fibroblasts and derive hiPSCs (n° 2017/167).

METHODS DETAILS

***Culture of hESCs/hiPSCs***

The hESCs and hiPSCs were cultured using standard procedures for feeder-free cultures and maintained in mTeSR plus medium (StemCell Technologies) on geltrex (Thermo Fisher) coated plates at 37°C, 5% CO2, and ambient oxygen level. Passaging was done using EDTA (Sigma). Cultures were routinely tested for Mycoplasma contamination and maintained Mycoplasma-free. A total of four control cell lines (hESC H9, CCP1^WT#1^; hiPSCs GM23338D, CCP1^WT^ ; GM23340D, CCP1^WT#3^ ; isogenic control of CCP1^Mut#2^ , CCP1^Iso#2^) and three hiPSC lines derived from fibroblasts collected from three patients with loss of function mutation for *AGTPBP1* gene leading to a loss or major reduction of CCP1 expression were used for the experiments (**Table S1**).

***Culture and generation of human forebrain organoids and forebrain assembloids***

Culture protocols to generate human dorsal and ventral forebrain organoids (hdFOs and hvFOs, respectively) were adapted from published work (*20, 21*). Briefly, hPSCs (hESCs or hiPSCs) (**Table S1**) were cultured until reaching 70–80% confluence. Cells were then dissociated into single-cell suspensions using accutase (StemCell Technologies), then 9,000 cells were seed per well in ultra-low-attachment 96-well plate (Nunclon Sphera). On day 0, cells were cultured in hPSC medium supplemented with FGF-2 (4 ng/ml) and Y-27632 (20 μM) (**Table S5**). On day 1, the medium was supplemented with the SMAD inhibitors LDN-193189 (100 nM; Sigma-Aldrich) and SB-431542 (10 µM; Tocris), as well as XAV939 (2 mM, StemCell,). The medium was replaced every other day until day 10. On day 3, FGF-2 was omitted and Y-27632 was reduced to 10 μM. On day 10, the medium was replaced by a neural induction medium (NIM; **Table S5**), to promote forebrain organoid differentiation. From day 10 to day 20, hdFOs and hvFOs were maintained in NIM, with medium changes every other day. NIM consisted of DMEM/F12 GlutaMAX (Thermo Fisher Scientific), 1% (v/v) N2 supplement (Thermo Fisher Scientific), 2% (v/v) B27 supplement without vitamin A (Thermo Fisher Scientific), and 0.1% (v/v) heparin (Sigma-Aldrich). hdFOs were cultured in NIM without additional morphogens, whereas hvFOs were cultured in NIM supplemented with Smoothened Agonist (SAG; 100 nM) and IWP2 (2.5 mM). On day 21, hvFOs and hdFOs were transferred from 96-well plate to 6-cm² dishes containing neuronal differentiation medium (NDM-A ‘young’; **Table S5**), with a maximum of 24 organoids per dish to limit spontaneous fusion. NDM-A ‘young’ consisted of a 1:1 mixture of DMEM/F12 and Neurobasal (Thermo Fisher Scientific), supplemented with 0.5% (v/v) NEAA, 0.5% (v/v) N2, 0.5% (v/v) GlutaMAX, 1% (v/v) Pen/Strep, 2% (v/v) B27 without vitamin A, and 250 µM insulin (Sigma-Aldrich) and 50 µM β-mercaptoethanol. Medium was changed every other day. Human forebrain assembloids (hFAs) were generated by culturing together 30 days old hdFOs and hvFOs of similar size into the same well of a round-bottom 96-well plate to promote fusion for one day. Once fused, resulting hFAs were transferred to 24-well plates containing neuronal differentiation medium (NDM-A ‘old’, **Table S5**) for growth and continued maintenance, with medium refreshed three times per week.

***Viral labeling of hvFOs prior fusion***

30-day-old human ventral forbrain oganoids (hvFOs) were transferred to an ultra-low-attachment 96-well U-bottom plate (Corning) containing 200 µL of NDMA-old medium and incubated overnight with DLX5/6-eGFP lentivirus (Lv-Dlx5/6-eGFP) at 37 °C and 5% CO₂. The following day, the medium was replaced with 200 µL of fresh NDMA-old medium. After one week, Dlx5/6-eGFP expression was detectablet, and the labeled hvFOs were subsequently fused with human dorsal forebrain organoids (hdFOs) to generate human forebrain assembloids (hFAs).

***Human brain tissue collection and preparation***

Human fetal brain tissues from gestational week (GW) 10-13 were collected immediately post-surgery and transported to the lab within 1 hour in preoxygenated artificial cerebrospinal fluid maintained on ice (aCSF: 85mM NaCl, 2.5mM KCl, 7mM MgCl2, 0.5mM CaCl2, 1mM NaH2PO4, 26mM NaHCO3, 10mM D-glucose, 50mM sucrose). Forebrain sections were embedded in 4% low-melting agarose diluted in HBSS and later processed for organotypic slices in chilled aCSF. MGE were dissected in cold oxygenated aCSF, and fully processed for explants preparation, or snap frozen in liquid nitrogen, and stored at -80°C until analysis.

***Mouse and human organotypic forebrain slice culture, lentiviral labelling, and time-lapse recording***

Organotypic slice cultures were performed following a protocol previously established in the laboratory (*22*). Briefly, heads from E13.5 mouse embryos were collected in ice-cold HBSS containing Ca²⁺ and Mg²⁺ (Biowest). Brains from human GW11-13 fetuses were collected in cold aCSF. Mouse and human brain were next embedded in 4% low-melting-point agarose diluted in HBSS. Brains were further sectioned into 250-µm-thick coronal slices using a Leica VT1000S/VT2000S vibratome. Slices were mounted on 0.4-µm Millicell (Millipore) culture inserts in FluoroDishes (WPI) immediately transferred to NDM-A ‘old’medium (**Table S5**).

To label and track cINs, slices were transduced overnight with lentiviral vectors Lv-Dlx5/6-eGFP or Lv-Dlx5/6-tdTomato on a shaker in a humidified incubator. The following day, slices were placed on 0.4-µm Millicell (Millipore) culture inserts in MatTek dishes (CELLView) containing NDM-A ‘old’ medium. Slices were maintained at 37°C and 5% CO₂ under ambient oxygen conditions for 48 hours before time-lapse recordings. For the analysis of cIN migration, recordings were performed for 16-24 hours using a Zeiss LSM 980 confocal microscope. Images were acquired every 20 minutes, with an average of 20-30 z-stacks per acquisition.

***Organotypic slice culture from human forebrain assembloids and time-lapse recording***

To assess cortical like-interneuron (clINs) migration in human forebrain assembloids (hFAs), 90-day-old hFAs were used. To label and track clINs, ventral forebrain organoids (hvFOs) were transduced at day 30 with the lentiviral vector Lv-Dlx5/6-eGFP, one week prior to fusion with dorsal forebrain organoids (hdFOs), as described above. At day 90, hFAs were embedded in 4% low-melting-point agarose diluted in HBSS containing Ca²⁺ and Mg²⁺ and sectioned into 250-µm-thick coronal slices using a Leica VT2000S vibratome. Slices were immediately transferred to NDM-A “old” medium (**Table S5**) and placed on 0.4-µm Millicell culture inserts in MatTek dishes containing NDM-A “old” medium. Time-lapse recordings were initiated within 4-12 hours after slicing and performed for at least 16 hours using a Zeiss LSM 980 confocal microscope. Slices were maintained at 37°C and 5% CO₂ under ambient oxygen conditions for 48 hours before time-lapse recordings. Images were acquired every 15 minutes, with an average of 20-30 z-stacks per acquisition, until the entire clIN signal was captured.

***Medial ganglionic eminence explant culture and time-lapse recording***

CELLview™ compartmented glass-bottom dishes were coated with poly-ornithine (0.1 mg/mL) for 45 min at 37°C, followed by incubation with laminin (5 µg/mL; Sigma-Aldrich) for 2 h at 37°C. A feeder layer of cortical cells was prepared as previously described (*22*). Cells were plated onto the coated glass-bottom dishes and allowed to adhere for 2 h at 37°C in a humidified incubator with 5% CO₂. Neurobasal complete medium, consisting of Neurobasal medium supplemented with 1% GlutaMAX, 1% N2 supplement, 2% B27 supplement, and 1% penicillin/streptomycin, was then gently added to the cells. The dishes were maintained in the incubator for 24 hours. The following day, MGE explants dissected from mouse (E13.5), ferret (E34-E36), or human (GW11-13) forebrains were dissected in HBSS supplemented with Ca²⁺ and Mg²⁺ (mouse and ferret) or aCSF (human). Explants were cut into pieces of approximately 100-300 µm^2^ and collected using a P10 pipette. The explants were centrifuged at low speed, and the medium was replaced with NDMA ‘old’ medium. Lentivirus-based labeling (Dlx5/6-eGFP, CAG-mito-Cherry, PGK-LifeAct-mRFPruby lentivirus, among others) was performed at this stage using a multiplicity of infection (MOI) of 15-25. The lentiviral preparation was added to the explants, which were incubated for at least 4 h to allow transduction. The explants were washed three times by low-speed centrifugation to remove residual lentiviral particles and then plated onto the cortical feeder layer.

Time-lapse imaging was performed 48 hours after plating for mouse and ferret explants and at least 72 hours after plating for human explants, using a Zeiss LSM 980 confocal microscope or Nikon Spinning DISC (GIGA imaging platform, ULiège; UMH Alicante). All recordings were performed for a minimum of 16 hours. Images were acquired every 15 min, with an average of 10 z-steps per time point. For all experiments, the dishes were maintained at 37°C in a humidified chamber with 5% CO₂.

Explants transduced with Lv-PGK-LifeAct-mRFPruby were imaged using Airyscan mode every 20 s. For MLCK inhibition, ML7 hydrochloride was added to the culture medium at a final concentration of 10 mM, 5 h before the start of the time-lapse recording, as previously described (*22*).

For hvFO explants, hvFOs aged 35-60 days were dissected into explants, typically 4-5 hvFOs per condition. Explants were then labelled with lentivirus as described above and subsequently placed onto the cortical feeder layer. Time-lapse imaging was performed 72 hours later. For mitochondrial labelling, explants were labelled using Lv-CAG-mito-mCherry lentivirus, and then clINs were imaged every 25 seconds, for 10 minutes.

***Immunohistochemistry***

*Forebrain cryosections.* Embryonic mouse (E13.5), human (GW11-13), and ferret (E34-E36) brains were dissected, fixed overnight in 4% paraformaldehyde (PFA) at 4°C, and cryoprotected overnight in 30% sucrose in PBS at 4°C. Brains were embedded in Neg-50 Frozen Section Media (Neg-50) and frozen on dry ice. Coronal sections were cut using a CryoStar NX70 cryostat (Epredia) at thicknesses of 12, 20, and 50 μm for mouse, human, and ferret samples, respectively. Sections were mounted onto SuperFrost Plus adhesive slides and stored at -80°C.Mouse and human sections were washed three times and blocked for 1 hour at room temperature (RT) in PBS containing 10% normal donkey serum (NDS), 0.2% Tween-20, and 0.1% Triton X-100. Sections were incubated overnight at 4°C with primary antibodies in blocking buffer with PBS, washed three times, and incubated with Alexa Fluor–conjugated secondary antibodies (1:500) for 2 hours at RT. Three additional washes followed; all washes used PBS containing 0.05% Tween-20. Ferret sections were washed twice in PBS and three times in PBS containing 0.25% Triton X-100, then blocked for 2 hours at RT in PBS containing 10% NDS, 0.25% Triton X-100, and 0.2% gelatin. Primary antibodies were applied in blocking buffer overnight at 4°C, followed by 1 hour at RT. Sections were washed four times, incubated with secondary antibodies in blocking buffer for at least 3 hours at RT, and washed four additional times in PBS containing 0.25% Triton X-100.

Primary antibodies were used at the following concentrations: rabbit anti-RFP 1:1000, rat anti-GFP, 1:1000, mouse anti-Lhx6 1:200, goat anti-GFP 1:1000, mouse anti-Cxcl12 1:500, rabbit anti-CCP1 1:500.

*Electroporated mouse brain sections.* Electroporated mouse brains were dissected, fixed overnight in 4% PFA at 4°C, embedded in 4% low-melting-point agarose in PBS, and sectioned coronally at 100 μm using a vibrating microtome (Leica VT1000S). Sections were stored in PBS containing 0.05% sodium azide at 4°C. Before staining, sections were permeabilized for 30 min in PBS containing 0.2% Triton X-100 and blocked in the same solution supplemented with 10% NDS and 5% bovine serum albumin (BSA). Primary antibodies were applied for 2–3 days at 4°C in PBS containing 0.2% Triton X-100, 5% NGS, and 5% BSA. Primary antibodies were used at the following concentrations: goat anti-mCherry 1:500, rat anti-GFP, 1:1000, mouse anti-Lhx6 1:200, rabbit anti-CCP1 1:500. After five washes, sections were incubated with secondary antibodies for 2 hours at RT in PBS containing 0.2% Triton X-100, 2.5% NGS, and 2.5% BSA. Sections were subsequently washed three times. All washes used PBS containing 0.05% Tween-20.

*Human forebrain assembloids (hFAs).* hFAs were fixed in 4% PFA for 45 min at RT and cryoprotected in 30% sucrose in PBS at 4°C. Three to five hFAs per condition were embedded in Neg-50, frozen in chilled isopentane, and stored at −80°C. Sections (14 μm) were mounted onto SuperFrost Plus adhesive slides, air-dried overnight at RT, and stored at -20°C. Before staining, sections were washed twice in PBS, subjected to antigen retrieval in Antigen Retrieval Solution at 90°C for 15 min, and washed three times in PBS. Sections were permeabilized and blocked for 1 hour at RT in PBS containing 1% BSA, 5% NDS, and 0.5% Triton X-100. Primary antibodies were applied overnight at 4°C in PBS containing 3% BSA, 5% NDS, and 0.3% Triton X-100. After four washes in PBS containing 0.1% Tween-20, sections were incubated with secondary antibodies in the same antibody diluent for 2 hours at RT.

For all sample types, nuclei were counterstained with DAPI, and sections were mounted using an antifade fluorescence mounting medium. Images were acquired using a Zeiss LSM 980 confocal microscope with 20×, 40×, or 60× objectives or a Axioscan 7 slide scanner with a 20× objective.

*Medial ganglionic eminence explant cultures*. Cells were fixed in 4% PFA for 15 min and permeabilized and blocked in dPBS containing 0.3% Triton X-100 and 10% donkey serum for 30 min at RT. Cells were then incubated overnight at 4°C with primary antibodies diluted in PBS containing 0.3% Triton X-100 and 1% donkey serum.

Primary antibodies were used at the following concentrations: rabbit anti-CCP1 1:500, rabbit anti-polyE 1:500, mouse anti-TUJ1 1:1000, rabbit anti-BetaIII tubulin 1:1000, mouse anti-DCX 1:500, goat anti-mCherry 1:500, rat anti-GFP, 1:1000, goat anti-GFP 1:1000. The following day, cells were incubated with secondary antibodies coupled to Alexa fluorophores (1:500) and DAPI (20µg/mL) for 2 hours at room temperature. Images were acquired using a Zeiss LSM 980 confocal microscope with a 20X, 40X or 63X objective. For the analysis of DCX and microtubules, MGE explants were dissociated and cINs were washed once with 1× PHEM buffer at RT and then incubated for 2 min with PHEM-P (1× PHEM, 0.5% Triton X-100, and 10% glycerol). CINs were subsequently fixed in 4% PFA diluted in PHEM, then washed with dPBS. Immunolabeling was performed as described above. Primary antibodies were used at specific concentration cited above. Images were acquired using a Zeiss LSM 980 confocal microscope equipped with Airyscan super-resolution imaging

*Immunofluorescence analysis of CCP1 expression and tubulin polyglutamylation in HEK293 cells*

HEK293 cells were co-transfected using Lipofectamine with pCMV-Cre and either loxStoplox-CCP1^WT-^Cherry or loxStoplox-CCP1^Dead-^Cherry constructs (**Table S4**). Cells were fixed in 4% paraformaldehyde (PFA) for 15 min and processed for immunofluorescence staining using primary antibodies against CCP1 (rabbit, 1/1000) or polyglutamylation (PolyE, rabbit 1/500), plus antibody anti-BIII Tubulin (Tuj1 clone, 1/1000) as indicated. Nuclei were counterstained with DAPI. Images were acquired using a Zeiss LSM 980 confocal microscope equipped with Airyscan super-resolution imaging.

***Western Blotting***

*Human ventral forebrain organoids.* Between 8 and 12 hvFOs were pooled per condition and lysed in 100-200 μl of RIPA lysis buffer (50 mM Tris-HCl, 150 mM NaCl, 1 mM EDTA, 1% Triton X-100, 1% Sodium Deoxycholate, 0.1% SDS) containing phosphatase and protease inhibitors. On ice, samples were mechanically dissociated using a pestle and incubated for 30 min. Lysates were centrifuged at 10,000 × g for 10 min at 4°C, and protein concentration in the supernatant was determined by BCA assay according to the manufacturer’s instructions. Samples were stored at -20°C until use. 20 µg of total protein per sample was diluted in 2X sample buffer, denatured at 95°C for 5 min (or at 37°C for 30 min for mitochondrial proteins), and resolved by SDS–PAGE on 10% polyacrylamide gels. Proteins were transferred to methanol-activated PVDF membranes at 300 mA for 2 hours. Membranes were blocked in TBS-T (TBS containing 0.1% Tween-20) with 5% BSA for 1 hour at 4°C and incubated overnight at 4°C with primary antibodies using the following dilutions: rabbit anti-CCP1 1 :1000, mouse antti-DCX 1 :1000, mouse anti TUJ1 1 :1000, rabbit anti-PolyE 1 :1000, rabbit anti BIII-tubulin 1 :1000, Monoclonal rabbit anti-Alpha tubulin 1 :5000, mouse anti- Beta actin 1 :2000, Polyclonal rabbit anti-VDAC 1 :1000, Monoclonal mouse anti-OXPHOS 1 :1000. Membranes were washed three times for 10 min with TBS-T and incubated with the appropriate secondary antibodies (1:5000), followed by three washes of 10 min. Protein bands were detected using enhanced chemiluminescence (ECL). Membranes were detected on an Amersham ImageQuant 800 imaging system (Cytiva).

*Medial ganglionic eminence extracts from mouse, human and ferret forebrains.* Proteins from E13.5 mouse, GW11-12 human, or E36 ferret medial ganglionic eminences were extracted using Tris-HCl buffer (10mM Tris-HCl, pH 7.5, 150mM NaCl, 0.5mM EDTA, 0.5% NP-40) supplemented with protease, phosphatase and deacetylase inhibitors. Proteins were separated in 7% to 15% polyacrylamide gels. For western blotting, 20 µg of total protein per sample was diluted in 2X sample buffer, denatured at 95°C for 5 min, and resolved by SDS–PAGE on 10% polyacrylamide gels. Proteins were transferred to methanol-activated PVDF membranes at 300 mA for 2 hours. Membranes were blocked in TBS-T, 5% BSA for 1 hour at 4°C. Next, membranes were incubated with primary antibodies overnight in rotation at 4°C. The membranes were washed three times with Tris-buffered saline with 0.1% Tween (TBS-T) for 10 min each and incubated with secondary antibodies for 1 hour at RT. After three washes in TBS-T, we treated the membrane with the ECL Western Blotting Detection solution (Millipore, Billerica, MA, USA) for protein detection. Bands were quantified using ImageJ software. Primary antibodies were used at specific concentration specified above. Membranes were detected on an Amersham ImageQuant 800 imaging system (Cytiva).

***Proximity ligation assay (PLA)***

E13.5 mouse MGE explants, or explants from 60-day-old hvFOs, were transduced with a lentiviral vector expressing Lv-Dlx5/6-eGFP. Explants were cultured in four-compartment dishes coated with N-cadherin and fixed in 4% paraformaldehyde (PFA) for 10 min, followed by two washes in phosphate-buffered saline (PBS). Fixed explants were maintained in PBS until processing. Samples were processed using the Duolink® PLA fluorescence detection system to assess the proximity between DCX and βIII-tubulin, according to the manufacturer’s instructions, with minor modifications. Primary antibodies against doublecortin (DCX, Santa Cruz Biotechnology), βIII-tubulin (Cell Signaling Technology), and GFP (Nacalai Tesque) were diluted in Duolink® Antibody Diluent. Primary antibodies were incubated for 1 hour at RT followed by ON incubation at 4°C. After the amplification step, GFP and βIII-tubulin were detected by incubating samples for 2 hours at room temperature with Alexa Fluor 488–conjugated donkey anti-rat and Alexa Fluor 647–conjugated donkey anti-rabbit secondary antibodies, respectively (1:1000), diluted in Duolink® Antibody Diluent. The Alexa Fluor 488 signal was used to identify Dlx5/6-GFP-positive cells, whereas the Alexa Fluor 647 signal was used to define the βIII-tubulin-positive region of interest (ROI) for subsequent PLA quantification. Samples were then washed three times for 15 min in PBS. Nuclei were counterstained with DAPI (1:5000 in PBS) for 15 min at room temperature, followed by a final 5-min wash in PBS. Samples were maintained in PBS and protected from light until imaging. Images were acquired using an LSM 880 confocal laser-scanning microscope (Zeiss) using a 40× objective. 1 µm Z-stack images were acquired, and the z-sections included in the analysis were selected prior to image processing.

***RNA extractions and RT-qPCR***

For each biological replicate, 5-12 hvFOs were pooled. Total RNA was extracted using the Quick-RNA Miniprep Kit according to the manufacturer’s instructions. RNA concentration and purity were assessed using a NanoDrop spectrophotometer, and RNA samples were stored at

-80°C until further use. For reverse transcription, 500 ng of purified RNA was used to synthesize cDNA with the RevertAid H Minus First Strand cDNA Synthesis Kit and random hexamer primers, following the manufacturer’s instructions. The resulting cDNA was diluted to 3.125 ng/µL and amplified using GoTaq qPCR Master Mix. Each sample–primer pair was analyzed in technical triplicates. (**Table S3**).

***Single cell RNA sequencing analysis of human ventral forebrain organoids***

Eight to twelve 71 to 84-day-old hvFOs from two to three independent differentiations of hvFOs were processed. hvFOs were cut into four pieces and washed once with HBSS to remove necrotic tissue and dead cells. hvFOs were then incubated for 20 min at 37°C in 500 µL of dissociation solution containing 18 U/mL papain and 200 U/mL DNase I. Enzymatic dissociation was stopped by adding an equal volume of ovomucoid inhibitor, followed by mechanical dissociation using pipetting 25 times with an uncut P1000 tip. The resulting cell suspension was passed through a 40 µm cell strainer and incubated on ice for 5 min to allow dead cells to aggregate. Cells were then pelleted by centrifugation for 5 min at 4°C and resuspended in dPBS containing 0.04% BSA.

***Microtubules isolation and co-sedimentation assay***

The protocol developed by Minotti et al. (1991) (*23*) was adapted to isolate soluble and polymerized tubulin fractions from 60 days old hvFOs . A volume of 200 µL of lysis buffer (20 mM Tris-HCl, pH 6.8; 0.14 M NaCl; 0.5% Nonidet P-40; 1 mM MgCl₂; 2 mM EGTA; Roche protease inhibitor cocktail; 4 µg/mL Taxol) was added at room temperature to 2–3 hvFOs. Samples were homogenized twice for 5 s using a motorized pestle mixer. The homogenate was centrifuged for 5 min at 300 × g at 20°C. An aliquot (50 µL) of the resulting supernatant (S1) was carefully layered onto 100 µL of cushion buffer (80 mM PIPES, pH 7.0; 1 mM MgCl₂; 1 mM EGTA; 60% glycerol; 4 µg/mL Taxol) and centrifuged for 40 min at 100,000 × g at 20°C using a TLA55 fixed-angle rotor (TLA-100 ultracentrifuge, Beckman Coulter). The resulting supernatant (S3), corresponding to the soluble tubulin fraction, was frozen on dry ice, while the pellet (P3), corresponding to the polymerized tubulin fraction, was resuspended in 50 µL of lysis buffer and frozen on dry ice.

***Validation of DCX-targeting shRNAs by flow cytometry***

HEK293 cells were co-transfected with pCMV-EGFP-hDCX and pLv-EF1a-dTomato-H1-shRNA plasmid encoding either a control shRNA (shCTR) or an shRNA targeting hDCX (shDCX#1 or shDCX#2) using Fugene HD (Roche). EGFP fluorescence was analyzed 24 h later by flow cytometry using a BD FACSAria III flow cytometer and FACSDiva v8.0 software. EGFP fluorescence intensity was used to assess the effects of DCX-targeting shRNAs on EGFP-hDCX reporter expression. The shRNA sequences and identifiers are listed in **Table S4**.

***Sample preparation for Proteomics***

Ninety-day-old human forebrain assembloids (hFAs) were dissociated in a dissociation solution containing 18 U/mL papain and 200 U/mL DNase I and subsequently filtered through a 40µm filter. Dlx5/6-eGFP⁺ cells were FACS-sorted using MA900 (SONY). Sorting was performed in Semi-purity mode with 100-nozzle size. Breifley, 5,000 cells per sample for the CCP1^Mut#1^ and CCP1^Mut#2^ conditions and 7,000 cells per sample for the CCP1^WT#2^ control condition were recovered after FACS (n = 5 biological replicates per condition). Cells were collected in Total Recovery vials (Waters Corp., Milford, USA) and stored at -80°C until processing. To minimize sample losses, the whole preparation was performed in the collection vial (one-pot workflow), each reagent being added without pipetting up and down and mixed by a short centrifugation (1 min, 12,000 x g), which also allowed the recovery of condensation after each incubation. Samples were thawed at room temperature and heated at 90°C for 10 minutes without agitation. Nucleic acids were then digested with 1 unit of Benzonase (Merck, Germany, Cat. N: 70664-3) in 50 mM Tris-HCl pH 7.4 for 15 minutes at room temperature, and the samples were evaporated to dryness in a speed vacuum. Proteins were resolubilized and reduced in 6 µL of 5 mM dithiothreitol (DTT) in 50 mM Tris-HCl pH 7.4 for 30 minutes at 56°C, then alkylated by addition of 2 µL of 40 mM iodoacetamide (IAA, 10 mM final concentration) for 30 minutes at room temperature in the dark. The samples were digested in solution with 600 ng of trypsin (Promega Germany, 2 µL at 300 ng/µL in 50 mM Tris-HCl pH 7.4) overnight at 37°C. The reaction was stopped by addition of 1 µL of 5.5% trifluoroacetic acid (TFA) to reach a 0.5% final concentration. Each sample was spiked with 1 µL of a commercial mixture of protein digest standards originated from non-human biological material, the MassPREP™ Digestion Standard Mixture (Waters, Corp., Milford, USA), in 0.5% TFA, corresponding to 10 fmol of standard per injection to an instrument quality control. Sample injection order was randomized and 9 µL (≈75% of the digest) were injected per sample. Samples were injected onto an ACQUITY UPLC M-Class system (Waters, Corp., Milford, USA) coupled online with a timsTOF Pro 2 mass spectrometer (Bruker Daltonics, Bremen, Germany) through a CaptiveSpray nanoelectrospray ion source in positive ion mode. The chromatographic separation was performed in trap-and-elute mode with a 10 µL loop, using solvent A (0.1% formic acid in water) and solvent B (0.1% formic acid in acetonitrile). Peptides were first loaded for 2 min at 20 µL/min with 98% A / 2% B on an ACQUITY UPLC M-Class Symmetry C18 trap column (100 Å, 5 µm, 180 µm x 20 mm; Cat. No. 186007496; Waters, Corp., Milford, USA), then separated on an ACQUITY UPLC M-Class HSS T3 C18 analytical column (100 Å, 1.8 µm, 75 µm x 250 mm; Cat. No. 186007474; Waters, Corp., Milford, USA) maintained at 40°C and equipped with a CaptiveSpray emitter tip. Elution was performed at a flow rate of 1 µL/min using the following linear gradient: 0-1 min, 2% B; 1.5 min, 7% B; 20.5 min, 27% B; 22.5 min, 40% B; 23.5-25.5 min, 90% B; 26-30 min, 2% B (column re-equilibration). The total run time of each analysis was 30 min. Ion mobility and m/z values were calibrated using the Agilent ESI-L Tuning Mix (Agilent Technologies, Santa Clara, USA). The mass spectrometer was operated in data-dependent acquisition with parallel accumulation–serial fragmentation (DDA-PASEF). Ion mobility separation was performed from 0.75 to 1.25 V·s/cm² (1/K0) with a ramp time of 150 ms and an accumulation time of 150 ms (duty cycle ≈ 100%). Each acquisition cycle consisted of one full TIMS-MS scan followed by 6 PASEF MS/MS ramps, for a total cycle time of 1.1 s. Only precursors with a charge state between 2 and 5 and a minimal intensity of 1,000 cps were selected for MS/MS analysis, with a target intensity of 15,000. Singly charged ions were excluded from MS/MS selection using a filter based on their m/z and ion mobility values. The parameters for MS spectrum acquisition are: mass range from 100 to 1700 m/z. The isolation window was set to +/- 1 Da with a dynamic exlusion of 0.4 min. The collision energy was linearly ramped as a function of ion mobility from 20 eV at 1/K0 = 0.6 V·s/cm² to 59 eV at 1/K0 = 1.6 V·s/cm².

***In utero electroporation in mouse***

To label cINs, MGE–directed in utero electroporation was performed as previously described (*24, 25*), with some modifications. Timed-pregnant mice carrying Dlx5,6: Cre-GFP embryos at E13.5 were anesthetized with isoflurane, and their uterine horns were exposed through an abdominal incision. A solution containing either pCMV-LoxSTOPlox-CCP1^wt^ or pCMV-LoxSTOPlox-CCP1^dead^ plasmid DNA (2 μg/μl) in HBSS was injected into a lateral ventricle of each targeted embryo using a glass micropipette. Forceps-type electrodes (CUY650P3; NEPAGENE) were positioned across the embryonic head to target the MGE. Five 60 V pulses, each lasting 50 ms, were delivered at 950 ms intervals using an electroporator (BTX). Following electroporation, the uterine horns were returned to the abdominal cavity, and the abdominal muscle and skin were sutured closed. Embryonic brains were collected at E17.5, and only GFP-positive embryos were included in subsequent analyses.

***Oxygraphy***

Per replicate, 30 to 35 hvFOs, aged 85, 90 or 93-day-old, were dissociated using the Papain Dissociation System (Worthington Biochemical Corporation) in accordance with the manufacturer’s instructions. The final cell pellet was washed, resuspended in MiR05 buffer, and adjusted to a concentration of 500,000-2,000,000 cells/mL. One hundred microlitres of this cell suspension were injected per chamber of the oxygraphy O2K, with subsequent injections of the following compounds in a specific order until reaching the final concentration, depending on the purpose. For measuring Resting, maximum OXPHOS (phosphorylating) and ETC (non-phosphorylating) capacity: (1) F-catalase (Merck, Cat#C9322), 556 U/ml; (2) digitonin (Merck, Cat#300410), 10 μg/ml; (3) pyruvate (Merch P2256), 5 mM; (4) malate (Merck, Cat#02288), 0.5 mM; (5) ADP (Calbiochem, Cat#117105), 1 mM; (6) glutamate (Merck, Cat#G1251), 10 mM; (7) succinate (Merck, Cat#S2378), 10 mM; (8) CCCP (Merck, Cat#C2759), add in 1 μM increments through repeated injections until maximum respiration is reached; (9) rotenone (Merck, Cat#R8875), 75 nM; (10) glycerophosphate (Merck, Cat#G7886), 10 mM; (11) antimycin A (Merck, Cat#A8674), 250 nM; (12) sodium ascorbate (Merck, Cat#A7631), 2 mM; (13) N,N,N′,N′-Tetramethyl-p-phenylenediamine dihydrochloride (TMPD) (Merck, Cat#T3134) 0.5 mM; (14) cytochrome C (Merck, Cat#C7752), 10 μM; (15) sodium azide (Merck, Cat#S2002), 100 mM.

QUANTIFICATION, STATISTICAL AND BIOINFORMATICAL ANALYSES

***Time lapse recording analyses***

Average speed, amplitude, and frequency of nucleokinesis, as well as total nuclear pause time, were analyzed as previously described using the TrackJ plugin in ImageJ (*22, 26, 27*). The StackReg plugin in ImageJ was used to correct for minor drift during imaging when necessary (*28*).

Nuclear translocations (nucleokinesis) were defined as displacements ≥5 µm in mouse and ferret cINs and ≥10 µm in human cINs/clINs. These thresholds were established by manually measuring soma displacement during visually identified nucleokinetic events, characterized by somatic swelling followed by translocation. The resulting thresholds were subsequently applied to the ImageJ tracking analysis to identify nucleokinetic events. Nucleokinesis amplitude was defined as the mean significant nuclear displacement per cell (µm). Normalized nucleokinesis amplitude was calculated by normalizing the amplitude for each cell to the average soma length of the corresponding neuron. Nucleokinesis frequency was calculated as the number of significant nuclear displacements divided by the total recording time. Mean speed was calculated as total nuclear displacement divided by the total tracking time. Only cells undergoing at least two completed nucleokinesis events were included in the analysis. Growth cone frequency was assessed manually by counting the number of bifurcations of the leading process over time. To quantify the maximum number of neurites and the maximum length of the leading process and neurites, cINs and clINs were tracked using Imaris, and the frame corresponding to maximum neuronal extension was identified. At this time point, the length of the leading process, number of neurites, and neurite length were quantified. Sholl analysis was also performed at this time point, using a 20µm inner soma circle and extending the analysis to the end of the neuron. The area of actomyosin contraction was visualized with Fire look-up table and quantified using ImageJ software as previously described (*14*). For quantification of homotopic contacts between cINs, multiple z-stacks were acquired throughout the time-lapse recordings to enable three-dimensional analysis. The z-stacks were analyzed using the 3D tools in Imaris to assess direct cell-cell contacts. The number of contacts occurring during the recording period was quantified and expressed as contact frequency over time. The nature of each contact, classified as mutually repulsive or non-repulsive, was assessed according to the criteria described previously (*29*).

***Mitochondrial network and fusion-fission event analyses***

Time-lapse recordings were visually inspected to identify Dlx5/6-eGFP- and CAG-mito-Cherry-positive cINs. Cells were manually outlined based on eGFP expression, and mitochondrial dynamics were analyzed using Nellie (*30*). Movies were acquired every 30 s for 3-4 h. Nellie was used to quantify mitochondrial objects and average mitochondrial size, as well as the frequency of fusion and fission events over time.

***Mitochondrial distribution during nucleokinesis***

To quantify mitochondrial distribution during nucleokinesis, 16 to 20h time-lapse recordings were visually inspected to identify Dlx5/6-GFP-positive clINs undergoing nucleokinesis. For each event, representative frames corresponding to three stages were selected: pre-nucleokinesis (soma swelling), mid-nucleokinesis (somal translocation), and post-nucleokinesis (neurite extension and branching). For each stage, the total cell area and the area of each cellular compartment (anterior soma, posterior soma, and neurites) were measured in Fiji by manually defining regions of interest (ROIs). Total mitochondrial area and the mitochondrial area within each cellular compartment were quantified using the same ROIs.

The proportion of total cell area occupied by each compartment and the proportion of total mitochondrial area (TMA) within each compartment were calculated by dividing the respective compartment area by the total cell area or total mitochondrial area. To normalize differences in the relative size of each cellular compartment, the percentage of total mitochondrial area within each compartment was divided by the corresponding percentage of total cell area, yielding a normalized measure of mitochondrial enrichment.

***Histological analyses***

Fluorescence intensity was quantified using ImageJ (*15*). βIII-tubulin-positive regions were manually selected as regions of interest (ROIs), and DCX and βIII-tubulin fluorescence intensities were measured within the same ROIs. DCX fluorescence intensity was normalized to βIII-tubulin fluorescence intensity and expressed as a DCX/βIII-tubulin ratio in arbitrary units (a.u.).

LHX6-positive cells were quantified using QuPath software across the cortical region, from the pallium-sub pallium boundary to the cortical area. The cortical wall was divided into 10 equally sized bins along its radial axis, and the number of LHX6-positive cells was quantified in each bin. Cell distribution was expressed as the percentage of the total number of LHX6-positive cells within each bin. The same binning strategy and cortical region were used across species to allow direct comparison of LHX6-positive cell distribution.

***PLA analyses***

Image processing and quantitative analysis were performed in Fiji/ImageJ using a custom macro. In brief, for each image, selected z-sections were converted into sum-intensity projections. The DCX/βIII tubulin PLA channel was thresholded using the Otsu method, and the Alexa Fluor 647 channel was binarized using the Default automatic thresholding method. Binary masks generated from the PLA and Alexa Fluor 647 channels were then combined using a pixel-wise logical and operation to identify PLA-positive puncta overlapping the tubulin-positive region. ROIs corresponding to Dlx5/6-GFP-positive cells were manually delineated and applied to the corresponding Alexa Fluor 647 binary mask. DCX/βIII tubulin PLA signal was quantified as the total area occupied by PLA-positive puncta overlapping the tubulin-positive region, normalized to the total tubulin-positive area.

***RT-qPCR analyses***

Amplification data were acquired using the LightCycler 480 Instrument (Roche) and were analyzed by the comparative Ct method using the Ct values and the average value of PCR efficiencies obtained from LinRegPCR software. To allow direct comparison between species, TTLL1 and TTLL7 were amplified in mouse and human samples using the same primer pairs, designed in regions conserved between the two species (Supplementary **Table S3**). PCR efficiency was calculated by LinRegPCR from the log-linear phase of each individual reaction, and the mean efficiency was determined separately for each primer pair in each species. Gene expression was normalized to Tyrosine 3-Monooxygenase/Tryptophan 5-Monooxygenase Activation Protein Zeta (YWHAZ in human, Ywhaz in mouse) and/or hypoxanthine guanine phosphoribosyl transferase (HPRT1 in human, Hprt in mouse).

***Western blot analyses***

Membranes were detected on an Amersham ImageQuant 800 imaging system (Cytiva). The density of the bands revealed was quantified using the gel quantification plugin on ImageJ software. Absolute values were normalized on loading control (α-tubulin, βIII-tubulin) bands of the same samples run in the same gel. Results are expressed as percentage of control change.

***Proteome analyses***

Proteomic data analysis. For label-free quantification (LFQ), raw files (.d) were analyzed with FragPipe ver. 19.0 using MSFragger ver. 3.6 (*31*) Philosopher ver. 4.6.0 (*32*) and IonQuant ver. 1.8.9 (*33, 34*) MS/MS spectra were searched against the Uniprot reviewed Homo sapiens database (20,394 protein sequences), supplemented with common contaminants, the sequences of the MassPREP™ digestion standards and reversed decoy sequences. Strict trypsin specificity was used with a maximum of two missed cleavages and a peptide length of 7 to 50 amino acids (peptide mass 500 to 5,000 Da). Carbamidomethylation of the cysteines was set as fixed modification, and oxidation of methionines and acetylation of protein N-termini were set as variable modifications. The initial precursor and fragment mass tolerances were set at 20 ppm, with automatic mass calibration and parameter optimization, and isotope errors of 0, 1 and 2 were allowed. Peptide spectrum matches (PSMs) were rescored with MSBooster (*35*) using deep learning-predicted retention times and fragment spectra, and Percolator (*36*) and proteins were inferred with ProteinProphet (*37*), Peptide spectrum match (PSM), peptide and protein false discovery rates (FDR) were all set at 0.01 using a sequential filtering approach based on reversed decoy sequences. Quantification was performed with IonQuant using the MaxLFQ algorithm (47) with normalization and match-between-runs (MBR) enabled (m/z tolerance 10 ppm, retention time tolerance 0.4 min, ion mobility tolerance 0.05 1/K0, MBR retention time tolerance 1 min, MBR ion FDR 0.01). Proteomic data detected in at least 3 of the 5 samples per group were treated using Perseus ver. 2.0.3.0. Differentially expressed proteins were detected based on Welch t-test.

Significantly dysregulated proteins were first submitted to STRING for protein-protein interaction network analysis, using a high-confidence interaction score threshold (combined score > 0.7). Enrichment for cytoskeleton organization (*p* value: 4.39e-06 & FDR : 8.2e-04) and metabolic processes (*p* value: 4.83e-05 & FDR : 5.2e-03) GO/pathway terms were assessed within STRING's Analysis tab, ranked by FDR. They were then submitted to Ingenuity Pathway Analysis (IPA, Qiagen) for canonical pathway enrichment, using Fisher's exact test as implemented in IPA. Pathways were considered significant at *p < 0.05. Dysregulated proteins were cross-referenced against previously reported direct CCP1 substrates from the literature (*38, 39*), identifying proteins present in both the dysregulated protein list and existing direct-target datasets. Additionally, Given CCP1's substrate specificity for glutamate (E) and aspartate (D) residues, protein sequences were retrieved manually from UniProt and visually inspected for C-terminal E/D repeat stretches.

***Public Single-cell RNA-sequencing data processing and quality control***

*Human subpallium and GE datasets* - Human subpallium and GE scRNAseq datasets were retrieved from two publicly available datasets, comprising five samples spanning GW11-12-13 (**Table S6**) [GSE165388 (*2*), GSE135827 (*1*)]. Human pallium scRNA-seq data were retrieved from two publicly available datasets, comprising three samples (**Table S6**) [GSE310125](https://www.ncbi.nlm.nih.gov/gds/?term=GSE310125%5bAccession%5d) (*7*), EGAS00001004107 (*6*). Pallium_Human #1, Ensembl gene identifiers were converted to HGNC gene symbols by querying Ensembl Biomart, retaining only successfully mapped, non-duplicated symbols.

*Mouse CGE, LGE and MGE datasets* - Mouse CGE, LGE and MGE scRNA-seq datasets were retrieved from two publicly available datasets, comprising eight samples spanning embryonic days E13-13.5-14.5 (**Table S6**) [GSE104156 (*3*), GSE188528 (*4*), GSE190593 (*5*)]. Mouse pallium scRNA-seq data were retrieved from six publicly available datasets, comprising 16 samples (**Table S6**) [GSE152281 (*8*), GSE155677 (*9*), GSE123335 (*10*), GSE161690 (*11*), GSE107122 (*12*), GSE153162 (*13*)].

*Gene identifier harmonization* - Mouse gene symbols follow a sentence-case convention (e.g. Lhx6) incompatible with the human upper-case convention (e.g. LHX6). Therefore, mouse gene symbols were converted to upper case for convenience during preprocessing (e.g. Lhx6 to LHX6). However, cross-species integration was performed using a table of one-to-one orthologs built from Ensembl (see below).

*Per-sample quality control* – For each sample, Seurat objects were created considering genes expressed in a minimum of three cells and cells with a minimum of 200 features. Adaptive per-sample quality thresholds were applied using MAD-based bounds (median ± 3 × MAD) with hard thresholds. Applied threshold are detailed in **Table S6**. GSE135827 (Subpallium_Human#3, #4 and #5) did not undergo mitochondrial filtering, as mitochondrial genes were absent from this dataset as retrieved from GEO. Doublets were identified per sample using scDblFinder (*40*) and removed prior to normalization.

*Removal of erythroid contamination* - GSE165388 (Subpallium_Human#1 and #2) samples contained erythroid contamination. To identify erythroid cells, counts were temporarily log-normalized to perform module scoring on a 19-gene erythroid signature (*HBA1, HBA2, HBB, HBG1, HBG2, HBM, HBQ1, ALAS2, GYPC, GYPA, GYPB, AHSP, HEMGN, BLVRB, BPGM, SLC25A37, SLC25A39, SNCA, FAM210B*) (*41*) this normalization was not carried forward into subsequent processing steps. Cells with module scores exceeding median + 3×MAD were removed. This step was not applied to the other human subpallial samples, which showed negligible erythroid contamination.

*Individual sample preprocessing* - Variance stabilization and normalization were performed using SCTransform v2. Mitochondrial content was regressed out for all samples except GSE135827 (Subpallium_Human#3, #4 and #5). PCA was run using SCT variable features, retaining a maximum of 50 components, consistent with common practice in scRNA-seq workflows. The number of PCs retained was selected using a quantitative cumulative-variance criterion (*42*). A conservative threshold of ≥90% cumulative variance was applied, above the cited 70% baseline, to minimize the risk of discarding biologically relevant signal and to standardize dimensionality selection across the multiple datasets and species in this study, with a minimum of 30 PCs. Shared nearest-neighbor (SNN) graphs were constructed and clusters identified at a resolution of 0.5 using the Louvain algorithm (https://doi.org/10.1088/1742-5468/2008/10/P10008). UMAP embeddings were computed using the selected PC dimensions.

***Subpallium analyses***

*Removal of non-lineage contaminating populations* – Within subpallial samples, samples were screened for non-GE contaminating populations: endothelial cells (*CLDN5, PECAM1, FLT1*), pericytes (*RGS5, ACTA2*), microglia (*CX3CR1, P2RY12, TMEM119, AIF1, TYROBP*), meningeal fibroblasts (*COL1A1, DCN, LUM*), cortical glutamatergic neurons (*NEUROD6, TBR1, SATB2, FEZF2*), choroid plexus epithelium (*TTR, CLDN11, AQP1*), thalamic neurons (*TCF7L2, LHX9, ZIC1, GBX2, ZIC4*), and mature astrocytes (*GFAP, AQP4, S100B*)(*41, 43, 44*). Contaminating clusters were removed from the analysis.

*Cross-GE reclassification in mouse samples* – Within mouse subpallial sample, each sample was further investigated for cross-GE contamination using identity markers for MGE (*NKX2-1, LHX6, MAF, MAFB*) (*45, 46*), LGE (*FOXP1, FOXP2, MEIS2*) (*47*), and CGE (*NR2F2, NR2F1*) (*48*). This step was specific to mouse samples, which were surgically dissected by GE compartments prior to sequencing, allowing cross-compartment marker expression to be interpreted as contamination ; human subpallial samples were dissected as whole subpallium without compartment-level separation, so no equivalent baseline for identifying cross-GE contamination was available. Given known transcriptional gradients between GE compartments, only clusters with unambiguous marker enrichment were reclassified, to avoid over-correction and introducing biases.

*Human-mouse orthologous mapping -* To enable cross-species transcriptomic comparison, a table of human-mouse orthologs was retrieved from Ensembl BioMart (release 115 ; human GRCh38.p14, mouse GRCm39) by querying human Ensembl gene identifiers against mouse ortholog attributes (gene stable ID, gene name, homology type) (*49*). Orthology relationships were classified as one-to-one, combined many-to-one/one-to-many, or many-to-many. Each sample's genes were classified according to this table (**Table S7**). Genes belonging to many-to-many families, representing 0.34-1.12% of genes per sample, were excluded due to ambiguous family membership. For orthology groups containing multiple paralogs, the paralog with the highest mean expression across cells was retained as the representative gene, selected independently within each sample. Representative symbols were assigned as the human gene symbol for one-to-one relationships, or a composite label for selected paralogs; full paralog family membership was retained for reference. Prior to integration, only orthologous genes shared across all 13 samples were retained (9,602 one-to-one ; 203 combined many-to-one/one-to-many).

*Cross-species integration (Harmony)* - Samples were merged into a single object, SCTransform v2 was applied. As described above, PCA was run on 50 components using shared variable features and the number of PCs retained for downstream analyses was selected using the ≥90% cumulative-variance criterion (minimum of 30 PCs). Batch correction was performed using Harmony (*50*), with cell sample identity and species as grouping variables, using regularization parameters (theta) of 1 and 2, respectively. The higher theta assigned to species reflects the greater expected transcriptional distance between human and mouse relative to inter-sample variation within species.

*Cross-species integration quality* - A local inverse Simpson's index (LISI) score was computed on a representative subsample of 5,000 cells to assess the extent of cross-species mixing in the integrated embedding (*50*). Values close to 1 indicate poor mixing (cells cluster by species), while values close to 2 (the maximum for two species) indicate well-mixed populations. The species LISI score reached 1.6 out of a maximum of 2 (*50*). Conservation of biological structure following integration was assessed through two independent, non-circular approaches rather than embedding-derived metrics such as LISI or silhouette score on GE identity. Standard cluster-separation metrics were not used, as GE-compartment identity was itself derived from clustering the same Harmony-corrected embedding, making any such metric circular. Instead, conservation was assessed via expression of markers not used in the original GE-identity classification (such as *SST, SIX3, NPY*, and *TAC1*), and via concordance with dissection-derived GE identity in mouse samples, an independent ground-truth label obtained prior to any computational processing (see below).

*GE subregion assignment* - Following integration, SNN graphs were constructed, clusters identified, and UMAP embeddings computed on the Harmony-corrected PCs. GE subregion identity was assessed using module scores of the MGE, CGE, and LGE gene signatures defined above, computed on the full integrated object. Cells were clustered at resolution 1, and each cluster was assigned a dominant GE identity (MGE-, LGE-, or CGE-enriched, or Progenitor) based on module score distributions and canonical marker expression. Concordance between computationally predicted and surgical dissection-derived GE identity was assessed in mouse datasets, for which dissection labels were available (**Table S8**), showing 92% concordance for MGE, 96% for LGE, and 44% for CGE. The comparatively lower CGE concordance reflects genuine biological ambiguity: CGE and LGE share overlapping transcriptional identity and specification factors.

*Post-mitotic DLX5/6 cells isolation* - To focus on DLX5/6 expressing migrating INs the integrated object was subset to G1-phase cells expressing *DLX5* or *DLX6* above a normalized expression threshold of 1. As the dataset spans the full continuum of GE development, *DLX5/6* expression increases gradually along the transition from progenitor to postmitotic INs identity rather than switching on discretely. A threshold above baseline detection was therefore applied deliberately, rather than any-detectable-expression (>0), to exclude transitional cells with ambiguous, low-level *DLX5/6* signal and retain only cells with clear, robust expression consistent with committed migrating INs identity.

*DEG investigation in post mitotic DLX5/6 MGE cINs* - MGE-enriched cells were subset from the postmitotic integrated object, retaining only cells derived from MGE-enriched samples. Raw counts were aggregated per sample to generate eight pseudobulk samples; five human samples, each corresponding to one individual subpallial or GE sample, and three mouse samples, each corresponding to one mouse dataset with pooled GE subregions pooled per dataset. Lowly expressed genes were filtered using edgeR's filterByExpr framework (*51*), retaining 9,178 genes. PCA on variance-stabilized counts (vst) (*52*) confirmed species as the dominant source of variance, with dataset of origin as a secondary axis. PCA on variance-stabilized counts (vst) (*52*) showed species as the main source of variation between samples, followed by dataset of origin. To ensure that differences observed between human and mouse were not confounded by technical differences between the individual datasets used, this variation was corrected for using RUVg, a method designed to estimate and remove such "batch effects" (*53*). For this correction to be effective, it requires a set of reference genes expected to vary between datasets but not between species, yielding a ‘W1’ factor. Genes with low overall variability and weak association with species identity were therefore selected as negative controls, ensuring the correction was designed to target dataset-level technical variation rather than genuine biological differences between species.

Differential expression was tested using edgeR's quasi-likelihood framework (model: ~ W1 + species; TMM normalization), with genes considered differentially expressed at FDR < 0.05 and |log2FC| > 0.25, 2,968 DEGs (1,422 up in human, 1,546 up in mouse). To assess robustness given the imbalance between human (n=5) and mouse (n=3) samples, the analysis was repeated on a 3-vs-3 subset (the three GE human samples vs. three GE mouse samples), yielding 2,676 DEGs, of which 2,291 were shared with the full analysis (Jaccard similarity = 0.68), indicating the core transcriptional differences were not primarily driven by the sample-size imbalance or the subpallium-dissected human samples.

The DEG list was submitted to Ingenuity Pathway Analysis (IPA, Qiagen). Canonical pathway enrichment was assessed by Fisher's exact test; the 'Nervous System Development and Function' category was examined from Disease and Function results, with activation z-scores (>0.5 activated, <−0.5 inhibited). Terms related to nervous system morphogenesis, neurite branching, neuritogenesis, and motor function were extracted, and their molecule union defined the migration/morphology gene set.

Over-representation analysis (ORA, clusterProfiler enrichGO) (*53*) was performed on this gene set against gene ontology term “biological process” GO:BP (org.Hs.eg.db), using the full DEG list as background universe; redundant terms were simplified (semantic similarity cutoff 0.7). Significant terms (BH-adjusted p < 0.05) were annotated into pathway families including cytoskeleton. Genes contributing to significant microtubule-related Gene Ontology (GO): Biological process (BP) terms were extracted from the cytoskeleton ORA output.

***Pallium analyses***

*Cell-type annotation in pallial samples* -Within pallial sample, AP (*PAX6, SOX2, HES1, VIM, NES*), IP (*EOMES, TBR2, NEUROG2, INSM1*), oRGC (*HOPX1, HEY1*), PN (*NEUROD2, NEUROD6, TUBB3*), Cajal-Retzius cells (*RELN, CALB2, LHX5)*, IN undetermined (*DLX1, DLX2, DLX5, DLX6, ASCL1*) (*54*):, MGE- (*LHX6, NKX2-1, SST, MAF*), CGE- (*NR2F1, NR2F2, SP8, VIP*), and LGE-derived migrating INs (*FOXP1, FOXP2, MEIS2*) (*55*), OPCs (*OLIG1, OLIG2, PDGFRA, SOX10*) (*56, 57*), microglia (*CX3CR1, P2RY12, TMEM119, AIF1, TYROBP*) (*58*), and other (e.g., blood (*CLDN5, PECAM1, FLT1*), pericytes (*RGS5, ACTA2, PDGFRB*) (*59, 60*). Annotation was performed on individual samples and integrated samples. Annotating separately prior to integration serves as a control for the integration step: if integration fails, the independently identified populations from each sample will not overlap in the integrated embedding

*Pallium, within species integration and processing -* For each species, pallium samples were merged into a single Seurat object following individual per-sample SCTransform v2 normalization. Integration features were selected across the constituent datasets using 3,000 features and assigned as variable features for the merged object. PCA was run on 50 components using these shared variable features. Batch correction was performed using Harmony, with sample identity (orig.ident) as the grouping variable and a regularization parameter (theta) of 1. For each species, UMAP embeddings were computed. SNN graphs were constructed, and clusters were identified at a resolution of 0.5 using the Louvain algorithm.

*Cell type annotation* - Cell populations were identified in each integrated object using canonical marker genes used for the no integrated samples described here above. For each population, module scores were computed using AddModuleScore (nbin = 15) on the corresponding marker set, and cluster identity was assigned based on module score enrichment together with expression of individual marker genes. Clusters displaying mixed or ambiguous marker expression were subset and independently reclustered to resolve population identity at higher resolution. Contaminating and lower-quality clusters were removed from downstream analyses.

*CellChat analysis* **-** Cell-cell communication was inferred independently within the human and mouse cortical datasets using CellChat. For each species, MGE- and CGE-derived INs were pooled into a single cIN population group, with pericyte/meningeal/endothelial populations excluded prior to CellChat object creation. A CellChat object was created from the RNA assay for each species using the species-matched CellChatDB ligand-receptor database (CellChatDB.mouse or CellChatDB.human), restricted to the default signaling subset (subsetDB). Standard CellChat preprocessing was applied, including identification of overexpressed genes and overexpressed ligand-receptor interactions within each cell population. Communication probabilities were computed using a truncated mean approach (trim = 0.1) with population size correction. Interactions supported by fewer than 10 cells were removed. Communication was extracted specifically for the cINs-to-cINs signaling axis by restricting both sources and targets to the pooled cINs population.

*Controls for sequencing depth, transcript complexity and population size* - Because human and mouse datasets differed in sequencing depth, the analysis was repeated after downsampling both species to a comparable median UMI count per cell (SampleUMI, max.umi = 4,800, upsample = FALSE) ; this threshold was set empirically to the median UMI count of the mouse dataset (4,815, rounded to 4,800), which served as the common lower reference. TThe human dataset resampled independently across three random seeds to account for stochasticity in downsampling. Interaction sets were compared pairwise across the three human runs using the Jaccard index, and interactions identified in all three runs were considered a robust "core" set for downstream comparison with mouse.

As a further control, a double-capping procedure was applied on top of UMI capping: for each cell exceeding a target number of detected genes, a random subset of non-zero entries was set to zero directly on the sparse count matrix until the target gene count was reached, without altering per-gene proportions among retained features. CellChat was then re-run on these UMI- and feature-capped matrices using the same processing parameters described above (trim = 0.1, min.cells = 10), including an increased bootstrap number (nboot = 50) for communication probability estimation.

To assess whether the higher number of detected interactions in human cINs reflected differences in group size rather than species-specific signaling, the mouse cINs population was additionally subsampled to match the human cINs population size (n = 920 cells), with all other populations retained at full size, and CellChat was re-run on this size-matched object.

Across all control analyses, the elevated number of significant ligand-receptor interactions detected in human cINs relative to mouse was maintained: downsampling to matched UMI depth yielded high concordance across the three human seeds (Jaccard similarity 0.94–0.98) without narrowing the species gap; double capping for both UMI and transcript complexity did not eliminate the discrepancy; and matching cIN population size in the mouse dataset to the smaller human cIN population further decreased, rather than increased, the number of mouse interactions detected, arguing against a sampling-size artifact as an explanation for the cross-species difference.

**Human ventral forebrain organoid (hvFO) single cell RNA sequencing**

Samples were multiplexed using TotalSeq™-B Hashtag antibodies 8, 9, and 10 (BioLegend, B0258, B0259, B0260) and pooled prior to library preparation. Libraries were generated using the Chromium GEM-X Single Cell 3' v4 platform (10x Genomics). Raw sequencing data were processed using Cell Ranger multi (v9.0.1) with a combined gene expression and antibody capture (hashtag) workflow, against the human reference genome (Ensembl, GRCh38, release 112), with intronic reads included. This yielded 15,414 (Pool 1, 83.1% recovery), 15,441 (Pool 2, 91.3% recovery), and 12,903 (Pool 3, 87.0% recovery) cells. Demultiplexed filtered feature-barcode matrices were converted into Seurat objects, retaining cells with a minimum of 250 detected features and genes detected in a minimum of 3 cells. Mitochondrial and ribosomal content were computed per cell. Quality control thresholds for hvFO samples were determined by iterative inspection of per-sample QC metric distributions. Thresholds were applied uniformly to remove low-quality cells and likely doublets: fewer than 5,000 features, fewer than 13,000 unique molecular identifiers (UMIs), less than 10% mitochondrial content, and less than 30% ribosomal content. Retrospective comparison against a standard median ± 3×MAD criterion, applied to the QC-filtered dataset, showed that only 0-2.6% of retained cells would have been additionally excluded under this stricter statistical threshold across all four metrics, indicating the empirical thresholds were reasonably conservative. Prior to integration, correct ventral patterning was confirmed by examining canonical dorsal markers (*EOMES, PAX6, TBR1, EMX1*) (*61*) and ventral markers (*NKX6-2, NKX2-1, DLX1, GAD1*) (*45, 62*) across samples. The second biological replicate of CCP1#2 was identified as aberrantly dorsally patterned and excluded from all downstream analyses. Per-sample normalization was performed using SCTransform (*63*), with mitochondrial content regressed out. Cell cycle phase was assigned using CellCycleScoring with S-phase and G2M gene lists from Tirosh et al. (*64*). PCA was run on 50 principal components using SCT variable features. UMAP embedding without integration confirmed strong sample- and cell-line-level batch effects requiring correction. Canonical Correlation Analysis (CCA), implemented via IntegrateLayers with SCT normalization, was selected for the final analysis based of cluster mixing across cell lines in UMAP space. Integration quality was assessed using the local inverse Simpson's index (LISI) computed on the CCA-integrated embedding. Overall median LISI was 1.89 (of a maximum of 3) grouped by cell line. LISI varied modestly across subclusters (cell-line LISI range: 1.11–2.30), with progenitor populations showing the highest mixing and GABAergic hypothalamic-like INs showing comparatively lower mixing, consistent with this being a smaller and more line-variable population. SNN graphs were constructed on the integrated embedding using 30 dimensions, and clusters were identified at resolution 0.6 using the Louvain algorithm [DOI: 10.1088/1742-5468/2008/10/P10008]. UMAP embeddings were computed on the CCA-corrected dimensional reduction. Cell types were identified by iterative clustering and marker-based annotation on the integrated dataset. Clusters were first identified at resolution 0.6 using the Louvain algorithm on 30 dimensions of the integrated (CCA) reduction, and UMAP embeddings were computed on the same dimensions. Major populations were identified: progenitors (*HES5, SOX2, HES1, VIM*) (*65*), dividing progenitors (*MKI67, TOP2A, CCNB2, PCNA*) (*64*), oligodendrocyte precursor cells (OPCs; *OLIG1, OLIG2, SOX10, PDGFRA*) (*66*), and interneurons (*GAD1, SNAP25*) (*62*). Clusters were assigned to one of five broad identities (Progenitors, Dividing Progenitors, OPCs, Interneurons). Each population was independently subset and re-processed to resolve finer substructure. Interneurons were re-clustered and five interneuron subtypes were identified: migrating (*IGFBP5, TNC, ZFP36, FOXA2, RFX4*) (*67, 68*), cortical-like (*GAD1, GAD2, SLC32A1,* *ARX, ASCL1*) (*2*), mature cortical like (*GABRA2, GABRB2, GABRG2, NEFL, NEFM*) (*2*), hindbrain-like (*TPH2, DDC, SLC6A4, FEV, GATA3*) (*69*), and hypothalamic-like (*LHX2, LHX9, NPVF, LMO3, VGLL2*) (*70*). Organoid-intrinsic cellular stress, arising from hypoxia, anaerobic glycolysis, and ER stress associated with limited oxygen diffusion in 3D culture and considered an in vitro artifact absent from fetal tissue, was identified using a manual implementation of the Gruffi approach (*71*), a semi-supervised method for detecting and removing stressed cells in organoid single-cell data based on glycolysis and ER-stress gene module scoring. Module scores were computed on normalized RNA data for three gene sets: glycolysis (*LDHA, ENO1, ENO2, GAPDH, PKM, PGK1, PGAM1, TPI1, ALDOA, HK2, PFKL, PFKM, GPI, LDHB*) and ER stress (*HSPA5, DDIT3, ATF4, XBP1, HERPUD1, DNAJB9, DDIT4, P4HB, ATF6, EIF2AK3, ERN1, TRIB3*) (*71*). Glycolysis and ER stress score distributions were inspected per cluster to define classification thresholds (glycolysis score > 0.65; ER stress score > 0.3), following the Gruffi framework's requirement that both signatures be elevated to distinguish genuine hypoxia-driven stress from other high-glycolysis states such as proliferating progenitors. Cells exceeding both thresholds were classified as stressed. The cluster dominated by stressed cells, together with cells classified as stressed outside of this cluster, were excluded from downstream analyses.

*Label transfer onto human GE atlas* ***-*** To ensure the cortical identity of the cortical-like interneurons (clINs), cells expressing *DLX5* or *DLX6* from the clIN cluster, with a normalized expression threshold of 1, were isolated. This query population was mapped onto the human subset of the cross-species integrated GE atlas (postmitotic DLX5/6+ population, MGE/CGE/LGE identity, as described above), using Seurat's anchor-based label transfer workflow. The reference was re-embedded with PCA and UMA, retaining the UMAP model to enable projection of query cells into reference UMAP space. Transfer anchors were identified using SCT-normalized data and the reference PCA reduction, without recomputing Pearson residuals for the query. GE subregion identity (MGE-, CGE-, or LGE-enriched) was transferred from the reference atlas via MapQuery, and query cells were projected into the reference UMAP embedding (*72, 73*). All query cells were retained regardless of prediction confidence. The proportion of query cells assigned to each predicted GE identity was computed.

*Velocyto and pseudotime computation of hvFOs* - To enable RNA velocity analysis, the Seurat object was exported for downstream processing in Python. Cell barcodes, UMAP coordinates, and full metadata were extracted and saved as a CSV file. The count matrix was exported in Matrix Market format, and PCA cell embeddings and gene names were saved as separate CSV files.

Spliced and unspliced transcript counts were quantified from Cell Ranger BAM output using velocyto (v0.17.17) (*74*), run per sample the reference annotation human transcriptome (release 112-7.1.0), generating one loom file per sample. The proportion of spliced, unspliced, and ambiguous reads was assessed per sample. Across samples, an average of 67.4% ± 1.2% of reads were spliced, 24.0% ± 1.1% unspliced, and 8.6% ± 0.2% ambiguous (mean ± SEM, n=8 samples). Cell barcodes in each loom file were reformatted to match the exported Seurat metadata and made unique across samples prior to concatenation. The exported Seurat object (count matrix, PCA embedding, UMAP coordinates, and metadata) was loaded into AnnData using Scanpy (*75*) and merged with the concatenated spliced/unspliced loom data using scVelo's merge utility (*76*).

The merged object was filtered and normalized, and first- and second-order moments were computed using default parameters. RNA velocity was estimated using the stochastic model. Velocity was visualized on the existing UMAP embedding, both as a grid and as a streamline projection, colored by cell subcluster identity. To estimate a velocity-based pseudotemporal ordering, full transcriptional dynamics were recovered using the dynamical model. Latent time is derived from this model's likelihood-based estimation of gene-specific transcription, splicing, and degradation rates and is not available under the stochastic model, so the dynamical model was required specifically for this step. Latent time was then computed, referred to as velocity pseudotime. Distributions of velocity pseudotime across subclusters were compared per cell line/condition. For condition-specific analyses, cells were subset by sample identity into experimental groups, and velocity was recomputed independently within the pooled control group, following the same filtering, normalization, and moment-estimation steps described above.

***Computational model of the cerebral cortex invasion by interneurons***

To obtain further insight into the influence of the various experimental observations, a 2D numerical model was designed to describe the migration of interneurons in both mouse and human brain slices (DOI: 10.5281/zenodo.22869350). The surrogates were designed as a series of nodes linked by harmonic oscillators. The basic geometry consists of three nodes, corresponding to the trailing process, the soma, and the leading process, respectively, each of which being attributed a mass (m1, m2, m3). During the evolution of the system, additional nodes can be added, either to the leading process or as level 2 and level 3 side branches. The intermediate nodes were assigned a ‘light’ mass (m3/10), whereas the terminal node of each branch was assigned a mass m3. Branches were created or suppressed with given probability at each time step. To maintain the geometry, successive nodes are also linked by three-body (angular) harmonic forces. For the sake of simplicity, all oscillators were assigned the same equilibrium length and harmonic force constants.

*Dynamics.* The temporal evolution of the surrogates was computed using molecular dynamics, with a time step chosen sufficiently small to allow the oscillations of the oscillators to be damped by friction and the whole system of coupled oscillators to evolve as an object with variable length and geometry. Surrogate movement was obtained by allowing trailing-process nodes and the head nodes of all branches, including the leading process, to jump to neighboring positions. These jumps were performed with a defined probability, sufficiently small to ensure numerical stability. The extent of each jump was sampled from a uniform distribution between 0 and 0.4 times the oscillator equilibrium length. Jumps occurred within an angle of 110 degrees from the orientation of the relevant oscillator. In addition, those surrogates that have only an extended leading process and no branches are given a probability to perform nucleokinesis, during which the soma and trailing process move halfway towards the leading process. This nucleokinesis is followed by a defined duration pause in the surrogate movement.

Due to the small time-step, rigid oscillator force constants, and friction, the surrogates diffuse in a regular manner. The parameters of the different models, human (H42), mouse (M44) and mouse-with gain of-function (M-GOF4) (**Table S2**). They differ in the frequency and pause duration for nucleokinesis and in the probabilities for creating/deleting level2/3 branches.

*Affinity.* The last aspect of the model was the brain geometry. A 2D template extending from the pallium to the end of the cortical plate was used to define the confining geometry of the system. Confining forces with a *r^-3^* decay were used to ensure smooth but efficient repulsive interactions with the limits of the system. The interaction with the biochemical cues known to drive migration (*54*) was modeled by incorporating Monte Carlo-type sampling into the jump probabilities. Specifically, the probability of a jump was modulated by the local value of an affinity field, which was predefined and kept identical for all simulations. The effective ‘temperature’ of these mixed molecular dynamics/Monte Carlo simulation was kept sufficiently high so that the fraction of jumps rejected due to a lower value of the affinity field remained below 1%. As an example, a very large temperature would lead to the effective simulation of a gas of surrogates that fills the entire 2D brain, whereas a 0K temperature setting forces all surrogates to move only according to the gradient of the local affinity field, therefore strongly confining the geometry, with some surrogates stuck eventually stuck in local minimal affinity regions. Throughout this simulation, the temperature is set to 0.05.

*Interaction between surrogates.* With such settings, defined an ensemble of surrogates behaves as a gas of independent, though of complex, evolving geometry, particles diffusing within a confining geometry. As shown in this paper (See Fig. 6) repulsion between branches was found to differ substantially between mouse and human. Therefore, the model was extended to include repulsive interactions between surrogates. The repulsive interaction has a *r^-3^* dependency, as contact forces are impractical for numerical reasons. This interaction occurs between any branch leading node of a surrogate and any node of any branch of a neighboring surrogate. This strong repulsion is set as active or inactive for a variable proportion of neurons (2% of repulsive interactions for Mouse model, 100% for the human model and 50% for Mouse-Gain-of-Function model).

***Quantification and statistical analyses.***

Quantification of measurements were performed using GraphPad Prism 9. Statistical differences between two experimental groups were evaluated using parametric or non-parametric t-tests, after verifying the normality of distribution for each group. Multiple comparisons were performed using one way-ANOVA or two-way ANOVA, plus indicated post-hoc test. Results are expressed as mean ±SEM. The experimental n, statistical test used and statistical significance for each experiment are indicated in the figure legends as well as in (**Table S9**).

**Supplemental tables**

**Table S1** List of hESC/hiPSCs used in this study

| **Cell line** | **Age at sampling** | **Sex** | **Genotype** | **Ethnicity** | **Source & identifier if available** |
| --- | --- | --- | --- | --- | --- |
| CCP1^WT#1^: hESC H9 WA09 | Blastocyst stage | Female | Wild type | N/A | RRID: CVCL_9773 |
| CCP1^WT#2^: hiPSC GM23338*D | 55-year-old | Male | Wild type | Caucasian | RRID: CVCL_F182 |
| CCP1^WT#3^: hiPSC GM23340*A | 44-year-old | Male | Wild type | Caucasian | RRID: CVCL_F184 |
| CCP1Mut^#1^: hiPSC  Family A | 15-months-old | Female | p.[M740fs]; [Y872*] | Not inbred, caucasian | Derived in-house; fibroblasts from Carsten Janke laboratory (Curie Institute) (*77*). |
| CCP1Mut^#2^: hiPSC  Family B | 15-months-old | Female | p.[Q816*]; [Q816*] | Inbred, persian |  |
| CCP1Mut^#3^: hiPSC  Family D | 12-year-old | Male | p.[T811M]; [H950L] | Not inbred, caucasian |  |
| CCP1^Iso#2 :^ isogenic  hiPSC | 5m | Male | Correction of p.Q856* |  | hiPSC corrected by the Synthego company |

Table S2 Computational model paramaters.

|  | Mouse (exp) | Human (exp, relative) | Mouse(sim) | M-GOF (sim, relative) | Human (sim, relative) |
| --- | --- | --- | --- | --- | --- |
| Amplitude NK (um) | 19.99 | 86% | 12.57 | 100% | 250% |
| Pause time NK (/15h) | 2.71 | 330% | 0.15 | 2400% | 1620% |
| Mean speed (um/h) | 74.5 | 24.8% | 74.5* | 63% | 42% |
| Frequency NK (n/hr) | 3.18 | 20.8% | 3.11 | 40% | 9% |
| Neuron size (um) | 87.2 | 181% | 44 | 100% | 250% |
| Growth cone splitting (n/hr) | 0.83 | 198% | 2.32 | 473% | 549% |
| Nr secondary br. | 2.91 | 236% | 1.48 | 187% | 215% |
| Nr tertiary br. | 3.2 | 4.48** | 0 | 0 | 5.81** |

* fixed values (defines time unit). The distance unit is fixed by the size of the cortical plate.

** absolute value

Table S3 List of qRT-PCR primers used in this study

| Name | Forward | Reverse |
| --- | --- | --- |
| CCP1 | GACACTCTTGCTGCATTGCT | GAATGAGCATGTTTCTATGCCG |
| DLX2 | ACGCTCCCTATGGAACCAGTT | TCCGAATTTCAGGCTCAAGGT |
| DLX5 | TTCCAAGCTCCGTTCCAGAC | GAATCGGTAGCTGAAGACTCG |
| EMX1 | CAGCTGGCCGGCAGTCT | TTGTACTTTGTCCTCCGGTTCTG |
| EMX2 | GCTTCTAAGGCTGGAACACG | CCAGCTTCTGCCTTTTGAAC |
| HPRT | ATTTCAAATCCAACAAGGTCTGGC | GCTTGCTGGTGAAAAGGACC |
| NKX2.1 | CGACTCGCTTCTCAGTGTCTGA | CCTCCATGCCCACTTTCTTG |
| PAX6 | GTGTCTACCAACCAATTCCACAAC | CCCAACATGGAGCCAGATG |
| TBR2 | CACCGCCACCAAACTGAGAT | CGAACACATTGTAGTGGGCAG |
| TTLL1 | GAAGAGCCCGTCCAGCAC | CCGGGACCACTTTTTGATCT |
| TTLL7 | ACGGAAGCAGAAAACTTTTATAGT | TCTTATCAAAGAAATCCCATGACCC |
| YWHAZ | GTAGGAGCCCGTAGGTCATC | TTCTCAGCACCTTCCGTCTT |

**Table S4** Plasmids generated in-house: sequences and reference

| Name | Reference | Sequence |
| --- | --- | --- |
| shCTR | RHS4346 | ATCTCGCTTGGGCGAGAGTAAG |
| shDCX#1 | TRCN0000422409 | GTTCCTCAGACAACTTCTTTA |
| shDCX#2 | TRCN0000418308 | GGTAACTTGTCTCCATGATTT |
| shCCP1#1 | TRCN0000275476 | AGCAATGCCAGAGTCTAATTA |
| shCCP1#2 | (*78*) | GCATAGAAACATGCTCATTCG |

Table S5 hvFOs, hdFOs and hFA media composition

| **Product** | **hPSC** | **NIM** | **NDMA Young** | **NDMA Old** |
| --- | --- | --- | --- | --- |
| DMEM/F-12, GlutaMAX™ supplement | 80% | 97% | 48% | 48% |
| KnockOut™ Serum Replacement | 20% | / | / | / |
| NeurobasalTM Medium | / | / | 48% | 48% |
| GlutaMAX™ Supplement | 1% | 1% | 0.5% | 0.5% |
| MEM Non-Essential Amino Acids Solution | 1% | 1% | 1% | 1% |
| Penicillin-Streptomycin | 1% | 1% | 1% | 1% |
| Beta mercapto ethanol | 1 mM | / | 0.5 mM | / |
| Heparin | / | 1 ‰ | / | / |
| N2 | / | / | 0.5% | 0.5% |
| B27 -VitA | / | / | 1% | 1% |
| Insulin | / | / | 2.5 µg/mL | / |

**Table S6** scRNAseq sample thresholds

| **Sample Name** | **Features** | | **Counts** | | **Mitochondria percentage** | | **Number of cells after quality control and contamination removal** |
| --- | --- | --- | --- | --- | --- | --- | --- |
|  | Low | High | Low | High | Low | High |  |
| **Human samples** | | | | | | | |
| Subpallium_Human#1: GSE165388 - subpallium - GW11 | 200 | 5,000 | 500 | 15,000 | 0 | 9.6 | 9,258 |
| Subpallium_Human#2: GSE165388 - subpallium - GW12 | 200 | 4,000 | 500 | 12,407 | 0 | 10 | 7,014 |
| Subpallium_Human#3: GSE135827 - GE - GW12 #1 | 200 | 3,000 | 500 | 7,709 | / | / | 3,927 |
| Subpallium_Human#4: GSE135827 - GE - GW12 #2 | 200 | 3,000 | 500 | 10,000 | / | / | 4,917 |
| Subpallium_Human#5: GSE135827 - GE - GW13 | 200 | 2,413 | 500 | 6,299 | / | / | 4,571 |
| Pallium_Human #1: EGAS00001004107 – GW13 – Cortex | 2,111 | 7,849 | 559 | 30,000 | 0 | 6.8 | 6,642 |
| Pallium_Human #2: GSM9289189 – GW13 – Telencephalon 1 | 0 | 8,000 | 0 | 28,000 | 0 | 1.5 | 8,012 |
| Pallium_Human #3: GSM9289190 – GW13 – Telencephalon 2 | 0 | 8,000 | 0 | 28,000 | 0 | 1.7 | 3,968 |
| **Mouse samples** | | | | | | | |
| Subpallium_Mouse#1 : GSE188528 - CGE - E13 | 1,484 | 3,272 | 2,470 | 8,812 | 0 | 4.6 | 3,438 |
| Subpallium_Mouse#2 : GSE188528 - LGE - E13 | 1,461 | 3,500 | 1,918 | 10,000 | 0 | 4.3 | 3,118 |
| Subpallium_Mouse#3 : GSE188528 - MGE - E13 | 1,608 | 3,414 | 2,886 | 9,708 | 0 | 4.5 | 3,966 |
| Subpallium_Mouse#4 : GSE104156 - CGE - E13.5 | 666 | 3,734 | 500 | 7965 | 0 | 2 | 5,584 |
| Subpallium_Mouse#5 : GSE104156 - MGE - E13.5 | 639 | 3,477 | 500 | 7,268 | 0 | 1.9 | 5,953 |
| Subpallium_Mouse#6 : GSE190593 - CGE - E14.5 | 1,021 | 4,086 | 500 | 11,663 | 0 | 7.4 | 4,696 |
| Subpallium_Mouse#7 : GSE190593 - LGE - E14.5 | 1,000 | 4,093 | 500 | 11,703 | 0 | 7.2 | 5,371 |
| Subpallium_Mouse#8 : GSE190593 - MGE - E14.5 | 1,000 | 4,157 | 500 | 12,124 | 0 | 7 | 5,316 |
| Pallium_Mouse #1 : GSM4610591 – E14 – Cortex + GE | 1,000 | 5,000 | 200 | 20,000 | 0 | 4.3 | 2,037 |
| Pallium_Mouse #2 : GSM4610592 – E14 – Cortex + GE | 1,000 | 5,000 | 200 | 20,000 | 0 | 6 | 2,742 |
| Pallium_Mouse #3 : GSM4610593 – E14 – Cortex + GE | 1,000 | 5,000 | 200 | 20,000 | 0 | 6 | 3,581 |
| Pallium_Mouse #4 : GSM4710393 – E14.5 – Somatosensory cortex | 2,000 | 7,500 | 100 | 40,000 | 0 | 8 | 2,785 |
| Pallium_Mouse #5 : GSM4710394 – E14.5 – Somatosensory cortex | 2,000 | 6,882 | 100 | 33,179 | 0 | 8 | 3,943 |
| Pallium_Mouse #6 : GSM3500949 – E14.5 – Neocortex | 0 | 4,500 | 0 | 10,000 | 0 | 3.1 | 1,222 |
| Pallium_Mouse #7 : GSM3500950 – E14.5 – Neocortex | 0 | 4,500 | 0 | 10,000 | 0 | 3.6 | 1,639 |
| Pallium_Mouse #8 : GSM3500951 – E14.5 – Neocortex | 0 | 4,218 | 0 | 8,504 | 0 | 3 | 2,002 |
| Pallium_Mouse #9 : GSM3500952 – E14.5 – Neocortex | 0 | 3,975 | 0 | 7,714 | 0 | 3 | 2,451 |
| Pallium_Mouse #10 : GSM3500947 – E14.5 – Neocortex | 0 | 4,379 | 0 | 8,801 | 0 | 2.7 | 1,142 |
| Pallium_Mouse #11 : GSM3500948 – E14.5 – Neocortex | 0 | 4,343 | 0 | 8,733 | 0 | 2.9 | 672 |
| Pallium_Mouse #12 : GSM4912739 – E14.5 – Neocortex | 0 | 2,802 | 0 | 4,851 | 0 | 4.6 | 1,678 |
| Pallium_Mouse #13 : GSM4912743 – E13.5 – Neocortex | 0 | 3,000 | 0 | 5,000 | 0 | 5 | 1,420 |
| Pallium_Mouse #14 : GSM2861511 – E14.5 – Neocortex | 0 | 3,000 | 0 | 5,500 | 0 | 6 | 1,406 |
| Pallium_Mouse #15 : GSM4635074 – E13.5 – Somatosensory cortex | 380 | 2,500 | 0 | 6,000 | 0 | 6.5 | 5,135 |
| Pallium_Mouse #16 : GSM4635075 – E14.5 – Somatosensory cortex | 575 | 4,000 | 0 | 12,000 | 0 | 5 | 2,610 |

**Table S7** Orthology classification

| Name | One to One | Many to One / One to Many | Many to Many |
| --- | --- | --- | --- |
| GSE165388 – subpallium – GW11 | 13,757 (96.34%) | 365 (2.56%) | 158 (1.12%) |
| GSE165388 – subpallium – GW12 | 13,160 (96.61%) | 327 (2.40%) | 135 (1.00%) |
| GSE135827 – GE – GW12 #1 | 11,734 (96.80%) | 285 (2.35%) | 103 (0.86%) |
| GSE135827 – GE – GW12 #2 | 12,305 (96.59%) | 314 (2.46%) | 120 (0.95%) |
| GSE135827 – GE – GW13 | 11,125 (96.80%) | 269 (2.34%) | 99 (0.87%) |
| GSE188528 – CGE – E13 | 11,043 (97.43%) | 253 (2.23%) | 40 (0.35%) |
| GSE188528 – LGE – E13 | 11,303 (97.43%) | 257 (2.22%) | 41 (0.35%) |
| GSE188528 – MGE – E13 | 11,233 (97.41%) | 255 (2.21%) | 43 (0.37%) |
| GSE104156 – CGE – E13.5 | 11,866 (97.49%) | 263 (2.16%) | 43 (0.35%) |
| GSE104156 – MGE – E13.5 | 11,412 (97.50%) | 253 (2.16%) | 40 (0.34%) |
| GSE190593 – CGE – E14.5 | 11,508 (97.36%) | 262 (2.22%) | 50 (0.42%) |
| GSE190593 – LGE – E14.5 | 11,552 (97.35%) | 268 (2.26%) | 47 (0.40%) |
| GSE190593 – MGE – E14.5 | 11,627 (97.37%) | 264 (2.21%) | 51 (0.43%) |

**Table S8** Correspondence between the surgical and transcriptional identities of mouse GE cells.

|  | Surgical CGE cells | Surgical LGE cells | Surgical MGE cells |
| --- | --- | --- | --- |
| Transcriptomically enriched CGE cells | 43.54% | 1.36% | 5.73% |
| Transcriptomically enriched LGE cells | 49.13% | 96.42% | 2.19% |
| Transcriptomically enriched MGE cells | 7.32% | 2.21% | 92.09% |

Table S9 Statistical table referring to all experiments analyzed in this work.

| **Fig. name** | **Additional information** | | **Statistical test and p-value** | **Value ± SEM** | **Replicates** |
| --- | --- | --- | --- | --- | --- |
| Fig. 1G |  | | Unpaired t test  **** p<0.0001 | Mouse is 30.86± 3.11 µm/h  Human is 10.54 ± 1.991 µm/h | 18 cINs from 5 E13.5 mouse embryos  15 cINs from 5 GW11-13 human fetuses |
| Fig. 1H |  | | Unpaired t test  **** p<0.0001 | Mouse is 1.399 ± 0.1783event/h  Human is 0.311 ± 0.049 event/h | 18 cINs from 5 E13.5 mouse embryos  15 cINs from 5 GW11-13 human fetuses |
| Fig. 1I |  | | Unpaired t test  ** (p=0.0024) | Mouse is 10.44 ± 1.626 h  Human is 22.5 ± 3.492 h | 18 cINs from 5 E13.5 mouse embryos  15 cINs from 5 GW11-13 human fetuses |
| Fig. 1J |  | | Unpaired t test  Ns (p=0.1545) | Mouse is 0.2525± 0.01072  Human is 0.2197 ± 0.02317 | 17 cINs from 55 E13.5 mouse embryos  9 cINs from 5 GW11-13 human fetuses |
| Fig. 1M |  | | Unpaired t test  **** p<0.0001 | Mouse is 74.54 ± 3.058 µm/h  Human is 18.49 ± 1.665 µm/h | 35 cINs from 4-5 E13.5 mouse embryos  34 cINs from 5 GW11-12 human fetuses |
| Fig. 1N |  | | Unpaired t test  **** p<0.0001 | Mouse is 3.18 ± 0.3185 event/h  Human is 0.662 ± 0.08173 event/h | 35 cINs from 4-5 E13.5 mouse embryos  34 cINs from 5 GW11-12 human fetuses |
| Fig. 1O |  | | Unpaired t test  Ns, (p=0.1821) | Mouse is 0.2251 ± 0.004  Human is 0.2162 ± 0.006 | 32 cINs from 4-5 E13.5 mouse embryos  24 cINs from 5 GW11-12 human fetusesX cINs from X embryos |
| Fig. 1P |  | | Unpaired t test  **** p<0.0001 | Mouse is 2.714 ± 0.1706 h  Human is 8.945 ± 0.3214 h | 35 cINs from 4-5 E13.5 mouse embryos  34 cINs from 5 GW11-12 human fetuses |
| Fig.  1Q |  | | Ordinary two-way ANOVA followed by Tukey’s multiple comparisons test.  Two-way ANOVA:  Interaction:  ***p<0.001  10 : **, p= 0.054  20: *** p = 0.0003  30: *** p = 0.0004  40: ****, p<0.0001  50: ****, p<0.0001  60: **, p =0.0012  70: ***, p< 0.012  80: ***, p = 0.0005  90: ** p=0.0014  100: * p = 0.019  110: *: ns, p =0.0382 | 10 : Mouse is 1.579±0.0118; human is 2.712±0.244  20: mouse is 2.102±0.323, human is 4.5±0.359  30: mouse is 2.158±0.289, human is 4.429 ±0.359  40: Mouse is 1.684±0.217; human is 4.429±0.343  50: Mouse is 1.421±0.207; human is 4.571±0.359  60: Mouse is 1.316±0.203; human is 3.929±0.486  70 : Mouse is 0.737±0.185; human is 4.071±0.569  80: Mouse is 0.526±0.160; human is 3±0.469  90: Mouse is 0.368±0.191; human is 2.429±0.488  100: Mouse is 0.020±0.011; human is 2.022±0.144  110: Mouse is 0±0.0; human is 1.429±0.402 | 21cINs from 4-5 E13.5 mouse embryos  21 cINs from 5 GW11-12 human fetuses |
| Fig. 1R |  | | Unpaired t test  ** p=0.0019 | Mouse is 0.8367± 0.1064 event/h  Human is 1.745± 0.2221 event/h | 26 cINs from 4-5 E13.5 mouse embryos  37 cINs from 5 GW11-12 human fetuses |
| Fig. 1S | Secondary neurite  Tertiary neurite | | One-way ANOVA  ***p<0.0001, plus Tukey’s multiple comparisons test.  Human cINS vs Mouse cINS  **** p<0.0001  Mouse cINs vs  Human cINS  **** p<0.0001 | Mouse is 3.05 ± 0.38  Human is 6.5 ± 0.29  Mouse is 1.5 ± 0.2157  Human is 4.9 ± 0.34 | 34 cINs from 4-5 E13.5 mouse embryos  22 cINs from 5 GW11-12 human fetuses  34 cINs from 4-5 E13.5 mouse embryos  22 cINs from 5 GW11-12 human fetuses |
| Fig. 1T |  | | Unpaired t test  ** p=0.0019 | Mouse is 0.7188 ± 0.019  Human is 0.6495 ± 0.02 | 21 cINs from 4-5 E13.5 mouse embryos  37 cINs from 5 GW11-12 human fetuses |
| Fig. 2F |  | | Mann Whitney test **p=0.0095 | Mouse is 1 ± 0.0109 a.u.  Human is 1.619 ± 0.2677 a.u. | 4 E13.5 mouse embryos  6 GW11-12 human fetuses |
| Fig. 2H |  | | Unpaired t test  * p=0.0270 | Mouse is 1± 0.2683  Human is 2.461 ± 0.6 | 9 E13.5 mouse embryos  6 GW11-12 human fetuses |
| Fig. 2I |  | | Unpaired t test  * p=0.0456 | Mouse is 1 ± 0.2370  Human is 0.3553 ± 0.08154 | 12 E13.5 mouse embryos  8 GW11-12 human fetuses |
| Fig. 2M |  | | One-way ANOVA  Ns; p=0.0769 | shCTR is 18.58 ± 1.366 µm/h  shCCP1 #1 is 15.96 ± 1.328 µm/h  shCCP1 #2 is 22.81 ± 2.043 µm/h | 40 cINs from 3-4 fetuses GW11-13 embryos for shCTR  shCCP1 #1  20 cINs from 3-4 fetuses GW11-13 embryos for shCCP1#1  10 cINs from 3 fetuses GW11-13 embryos for  shCCP1 #2 |
| Fig. 2N |  | | One-way ANOVA  ***p=0.0001, plus Tukey’s multiple comparisons test  shCTR vs shCCP1 #1: ***  (p=0.0002)  shCTR vs shCCP1 #2: ****  (p=0.0277)  shCCP1 #1 vs shCCP1#2: ns  (p=0.5786) | shCTR is 25.08 ± 1.082 µm  shCCP1 #1 is 16.78 ± 0.9459 µm  shCCP1 #2 is 19.20 ± 2.867 µm | 27 cINs from 3-4 fetuses GW11-13 embryos for shCTR  shCCP1 #1  16 cINs from 3-4 fetuses GW11-13 embryos for shCCP1#1  10 cINs from 3 fetuses GW11-13 embryos for  shCCP1 #2 |
| Fig. 2O |  | | One-way ANOVA  ***p=0.0001, plus Tukey’s multiple comparisons test  shCTR vs shCCP1 #1: *  (p=0.0402)  shCTR vs shCCP1 #2: ****  (p<0.0001)  shCCP1 #1 vs shCCP1#2: **  (p=0.0021) | shCTR is 9.915± 0.6353h  shCCP1 #1 is 7.507 ± 0.4088 h  shCCP1 #2 is 3.1± 0.8682 h | 30 cINs from 3-4 fetuses GW11-13 embryos for shCTR  shCCP1 #1  14 cINs from 3-4 fetuses GW11-13 embryos for shCCP1#1  10 cINs from 3 fetuses GW11-13 embryos for  shCCP1 #2 |
| Fig. 2P |  | | One-way ANOVA  ***p=0.0005, plus Tukey’s multiple comparisons test  shCTR vs shCCP1 #1: **  (p=0.0011)  shCTR vs shCCP1 #2: *  (p=0.0128)  shCCP1 #1 vs shCCP1#2: ns  (0.9108) | shCTR is 1.967 ± 0.2376event/h  shCCP1 #1 is 0.8381± 0.041event/h  shCCP1 #2 is 0.9884 ± 0.1296 event/h | 30 cINs from 3-4 fetuses GW11-13 embryos for shCTR  shCCP1 #1  14 cINs from 3-4 fetuses GW11-13 embryos for shCCP1#1  10 cINs from 3 fetuses GW11-13 embryos for  shCCP1 #2 |
| Fig. 3C |  | | One-way ANOVA  **p=0.0016, plus Tukey’s multiple comparisons test.  CCP1^WT#2^ vs  CCP1^MUT#2^: **  (p=0.0013)  CCP1^WT#2^ vs  CCP1 ^Iso#2^: ns  (p=0.0696)  CCP1 ^Iso#2^ vs  CCP1^MUT#2^: *  (0.0183) | CCP1^WT#2^ is 1± 0.05  CCP1^MUT#2^ is 0.2393± 0.0386  CCP1^Iso#2^ is 0.6829 ± 0.123 | N=3 differentiations, 8-12 organoids pooled |
| Fig. 3J |  | | Unpaired t test,  Ns (p=0.1473) | hFA clINs is 22.06 ± 1.762 µm/h  Human cINs is 18.49 ± 1.665 µm/h | 34 cINs from 3 GW12 fetuses  28 clINs from 10-12 assembloids (hFAs) of 4-5differentiation |
| Fig. 3K |  | | Unpaired t test,  Ns (p=0.2122) | hFA clINs is 8.244 ± 0.4730 h  Human cINs is 8.945 ± 0.3214 h | 32 cINs from 3 GW12 fetuses  26 clINs from 10-12 assembloids (hFAs) of 4-5differentiation |
| Fig. 3L |  | | Unpaired t test,  Ns (p=0.9907) | hvFOs clINs is 1.751± 0.1542 events/h  Human cINs is 1.748± 0.2064 events/h | 40 cINs from 3 GW11-13 fetuses MGE explants  25 clINs from 4-5 hvFOs explants of 3 differentiations |
| Fig. 3N |  | | One-way ANOVA  Ns, p=0.4001 | CCP1^WT#2^ is 24.65 ± 2.352 µm/h  CCP1^MUT#2^ is 19.35 ± 2.159 µm/h  CCP1^Iso#2^ is 23.77 ± 3.332 µm/h | 27 clINs from 4-5 hvFOs CCP1^WT#2^; 19 clINs from CCP1^MUT#2^,  22 clINs from 4-5 hvFOs CCP1 ^Iso#2^ 4-5 hvFOs explants of 3 differentiations |
| Fig. 3O |  | | One-way ANOVA  Ns, p=0.7252 | CCP1^WT#2^ is 0.8419 ± 0.089 events/h  CCP1^MUT#2^ is 0.6853 ± 0.159 events/h  CCP1^Iso#2^ is 0.7301 ± 0.1058 events/h | 25 clINs from 4-5 hvFOs CCP1^WT#2^; 18 clINs from CCP1^MUT#2^,  16 clINs from 4-5 hvFOs CCP1 ^Iso#2^ 4-5 hvFOs explants of 3 differentiations |
| Fig. 3P |  | | One-way ANOVA  ***p<0.0001, plus Tukey’s multiple comparisons test.  CCP1^WT#2^ vs  CCP1^MUT#2^: ****  (p<0.001)  CCP1^WT#2^ vs  CCP1 ^Iso#2^: ns  (p=0.2206)  CCP1 ^Iso#2^ vs  CCP1^MUT#2^: **  (p=0.0052) | CCP1^WT#2^ is 27.46 ± 1.4 µm  CCP1^MUT#2^ is 16 ± 1.343µm  CCP1^Iso#2^ is 23.52 ± 1.616 µm | 24 clINs from 4-5 hvFOs CCP1^WT#2^; 19 clINs from CCP1^MUT#2^,  21 clINs from 4-5 hvFOs CCP1 ^Iso#2^ 4-5 hvFOs explants of 3 differentiations |
| Fig. 3Q |  | | One-way ANOVA  ***p<0.0001, plus Tukey’s multiple comparisons test.  CCP1^WT#2^ vs  CCP1^MUT#2^: ****  (p<0.0001)  CCP1^WT#2^ vs  CCP1 ^Iso#2^: ns  (p=0.172)  CCP1 ^Iso#2^ vs  CCP1^MUT#2^: *  (p=0.023) | CCP1^WT#2^ is 8.324 ± 0.4774 h  CCP1^MUT#2^ is 4.798 ± 0.3563 h  CCP1^Iso#2^ is 6.913 ± 0.565 h | 27 clINs from 4-5 hvFOs CCP1^WT#2^; 19 clINs from CCP1^MUT#2^,  22 clINs from 4-5 hvFOs CCP1 ^Iso#2^ 4-5 hvFOs explants of 3 differentiations |
| Fig. 3R |  | | One-way ANOVA  ***p<0.0001, plus Tukey’s multiple comparisons test.  CCP1^WT#2^ vs  CCP1^MUT#2^: ****  (p<0.0001)  CCP1^WT#2^ vs  CCP1 ^Iso#2^: ns  (p=0.9047)  CCP1 ^Iso#2^ vs  CCP1^MUT#2^: ****  (p<0.0001) | CCP1^WT#2^ is 1.751 ± 0.1542 events/h  CCP1^MUT#2^ is 0.6512 ± 0.09611 events/h  CCP1^Iso#2^ is 1.841 ± 0.1819 events/h | 25 clINs from 4-5 hvFOs CCP1^WT#2^; 20 clINs from CCP1^MUT#2^,  18 clINs from 4-5 hvFOs CCP1 ^Iso#2^ 4-5 hvFOs explants of 3 differentiations |
| Fig. 3S |  | | One-way ANOVA  **p=0.0011, plus Tukey’s multiple comparisons test.  CCP1^WT#2^ vs  CCP1^MUT#2^: **  (p=0.0012)  CCP1^WT^ vs  CCP1 ^Iso#2^: ns  (p=0.7678)  CCP1 ^Iso#2^ vs  CCP1^MUT#2^: *  (p=0.0113) | CCP1^WT#2^ is 105.2± 5.3 µm  CCP1^MUT#2^ is 80.26 ± 4.3 µm  CCP1^Iso#2^ is 100.8 ± 3.8 µm | 22 clINs from 4-5 hvFOs CCP1^WT#2^; 17 clINs from CCP1^MUT#2^,  19 clINs from 4-5 hvFOs CCP1 ^Iso#2^ 4-5 hvFOs explants of 3 differentiations |
| Fig. 3T |  | | One-way ANOVA  *p=0.0161, plus Tukey’s multiple comparisons test.  CCP1^WT#2^ vs  CCP1^MUT#2^: *  (p=0.0205)  CCP1^WT#2^ vs  CCP1 ^Iso#2^: ns  (p=0.904)  CCP1 ^Iso#2^ vs  CCP1^MUT#2^: *  (p=0.048) | CCP1^WT#2^ is 54.12 ± 6.3 µm  CCP1^MUT#2^ is 28.9 ± 5.5 µm  CCP1^Iso#2^ is 50.57 ± 6.318µm | 7 clINs from 4-5 hvFOs CCP1^WT#2^; 7 clINs from CCP1^MUT#2^,  7 clINs from 4-5 hvFOs CCP1 ^Iso#2^ 4-5 hvFOs explants of 3 differentiations |
| Fig. 3U |  | | One-way ANOVA  ***p<0.0001, plus Tukey’s multiple comparisons test.  CCP1^WT#2^ vs  CCP1^MUT#2^: **  (p=0.0052)  CCP1^WT#2^ vs  CCP1 ^Iso#2^: ns  (p=0.1822)  CCP1 ^Iso#2^ vs  CCP1^MUT#2^: ****  (p<0.0001) | CCP1^WT#2^ is 3.833 ± 0.25  CCP1^MUT#2^ is 2.611 ± 0.2  CCP1^Iso#2^ is 4.48 ± 0.3133 | 24 clINs from 4-5 hvFOs CCP1^WT#2^; 19 clINs from CCP1^MUT#2^,  21 clINs from 4-5 hvFOs CCP1 ^Iso#2^ 4-5 hvFOs explants of 3 differentiations |
| Fig. 3V |  | | Unpaired t test  CCP1^WT#2^ vs  CCP1^MUT#2^: **  (p=0.00) | CCP1^WT#2^ is 10.34 ± 3.7  CCP1^MUT#2^ is 22.25± 1.94 | 7 clINs from 4-5 hvFOs explants CCP1^WT#2^; 11 clINs from CCP1^MUT#2^,  clINs from 4-5 hvFOs explants of 3 differentiations. |
| Fig. 3W |  | | One-way ANOVA  Ns, p=0.8560 | DMSO CCP1^WT#2^ is 24.86 ± 7 μm/h  DMSO CCP1^MUT#2^ is 18.88 ± 4.4 μm/h  ML7 CCP1^WT#2^ is 20.30 ± 3.5 μm/h  ML7 CCP1^MUT#2^ is 21.56± 3.034 μm/h | 7 clINs from 4-5 hvFOs explants CCP1^WT#2^ DMSO or Ml7  12 clINs from 4-5 hvFOs explants CCP1^Mut#2^ DMSO  15 clINs from 4-5 hvFOs explants CCP1^Mut#2^ Ml7 |
| Fig. 3X |  | | One-way ANOVA  **p-0.0073, plus Tukey’s multiple comparisons test.  DMSO CCP1^WT#2^ vs  DMSO CCP1^MUT#2:^ **  (p=0.0076)  ML7 CCP1^WT#2^ vs  DMSO CCP1^MUT#2:^ *  (p=0.0434)  ML7 CCP1^MUT#2^ vs  DMSO CCP1^MUT#2:^ *  (p=0.0135) | DMSO CCP1^WT#2^ is 9.375 ± 0.9953 h  DMSO CCP1^MUT#2^ is 3.94 ± 0.7115 h  ML7 CCP1^WT#2^ is 7.708± 0.8024 h  ML7 CCP1^MUT#2^ is 8.2 ± 0.7751h | 7 clINs from 4-5 hvFOs explants CCP1^WT#2^ DMSO or Ml7  12 clINs from 4-5 hvFOs explants CCP1^Mut#2^ DMSO  15 clINs from 4-5 hvFOs explants CCP1^Mut#2^ Ml7 |
| Fig. 3Y |  | | One-way ANOVA  ***p=0.001, plus Tukey’s multiple comparisons test.  DMSO CCP1^WT#2^ vs  DMSO CCP1^MUT#2:^ **  (p=0.0084)  DMSO CCP1^WT#2^ vs  ML7 CCP1^MUT#2:^ *  (p=0.0243)  ML7 CCP1^WT#2^ vs  DMSO CCP1^MUT#2:^ *  (p=0.0107)  ML7 CCP1^WT#2^ vs  ML7 CCP1^MUT#2:^ *  (p=0.0312) | DMSO CCP1^WT#2^ is 1.865 ± 0.33 events/h  DMSO CCP1^MUT#2^ is 0.644 ± 0.1482 events/h  ML7 CCP1^WT#2^ is 1.753 ± 0.24 events/h  ML7 CCP1^MUT#2^ is 0.9113± 0.1558 events/h | 10 clINs from 4-5 hvFOs explants CCP1^WT#2^ DMSO or Ml7  15 clINs from 4-5 hvFOs explants CCP1^Mut#2^ DMSO or  Ml7 |
| Fig. 4D |  | | Unpaired t test  CCP1^WT#2^ vs  CCP1^MUT#2:^ *p<0.05  (p=0.0354) | CCP1^WT#2^ is 0.48 ± 0.1164  CCP1^MUT#2^ is 1.806 ± 0.4084 | 7-8 hvFOs of 3 independents  differentiations |
| Fig. 4G |  | | One-way ANOVA  ***p<0.0001, plus Tukey’s multiple comparisons test.  CCP1^WT#2^ShCTR vs  CCP1^MUT#2^ShCTR^:^ *  (p=0.0119)  CCP1 ^MUT#2^Sh#1 vs  CCP1^MUT#2^ ShCTR^:^ ****  p<0.0001  CCP1 ^MUT#2^Sh#2 vs  CCP1^MUT#2^Sh#CTR^:^ **** p<0.0001 | CCP1^WT#2^ShCTR is 1.873± 0.1898 (event/h)  CCP1^MUT#2^ShCTR is 0.7015± 0.047 (event/h)  CCP1^MUT#2^Sh#1 is 2.768 ± 0.3245(event/h)  CCP1^MUT#2^Sh#2 is 2.572 ± 0.3142 (event/h) | 21clINs from CCP1^WT#2^ShCTR  19 clINs from CCP1^Mut#2^ShCTR  22 clINs from CCP1^Mut#2^Sh#1  22 clINs from CCP1^Mut#2^Sh#2  of 4-5 hvFOs of 3 independents  differentiations |
| Fig. 4H |  | | One-way ANOVA  ***p<0.0001, plus Tukey’s multiple comparisons test.  CCP1^WT#2^ShCTR vs  CCP1^MUT#2^ShCTR^:^ **  (p=0.0011)  CCP1 ^MUT#2^Sh#1 vs  CCP1^MUT#2^ ShCTR^:^ ****  p<0.0001  CCP1 ^MUT#2^Sh#2 vs  CCP1^MUT#2^Sh#CTR^:^ **** p<0.0001 | CCP1^WT^ShCTR is 3.231± 0.3586 vs  CCP1^MUT#2^ShCTR is 1.467 ± 0.1888  CCP1 ^MUT#2^Sh#1 is 4.288 ± 0.3568  CCP1^MUT#2^Sh#2 is 3.964 ± 0.3446 | 13 clINs from CCP1^WT#2^ShCTR  15 clINs from CCP1^Mut#2^ShCTR  13 clINs from CCP1^Mut#2^Sh#1  15 clINs from CCP1^Mut#2^Sh#2  of 4-5 hvFOs of 3 independents  differentiations |
| Fig. 4I |  | | One-way ANOVA  ***p<0.0001, plus Tukey’s multiple comparisons test  CCP1^WT#2^ShCTR vs  CCP1^MUT#2^ShCTR^:^ *** p<0.0001  CCP1^WT#2^ShCTR vs CCP1^MUT#2^Sh#1^:^ ***  *** p<0.0001  CCP1^WT#2^ShCTR vs CCP1^MUT#2^Sh#2^:^ ***  One way ANOVA p<0.0001 | CCP1^WT^ShCTR is 16.72± 1.093 µm vs  CCP1^MUT#2^ShCTR is 7.411 ± 0.2710 µm  CCP1 ^MUT#2^Sh#1 is 10.29 ± 1.252 µm  CCP1^MUT#2^Sh#2 is 8.705 ± 0.5246 µm | 13 clINs from CCP1^WT#2^ShCTR  15 clINs from CCP1^Mut#2^ShCTR  13 clINs from CCP1^Mut#2^Sh#1  15 clINs from CCP1^Mut#2^Sh#2  of 4-5 hvFOs of 3 independents  differentiations |
| Fig. 4J |  | | One-way ANOVA  ***p<0.0001, plus Tukey’s multiple comparisons test  CCP1^WT#2^ShCTR vs  CCP1^MUT#2^ShCTR^:^ *** p<0.0001  CCP1^WT#2^ShCTR vs CCP1^MUT#2^Sh#1^:^ ** p=0.0049  CCP1^WT#2^ShCTR vs CCP1^MUT#2^Sh#2^:^ ***p<0.0001 | CCP1^WT^ShCTR is 8.772± 0.8591 h vs  CCP1^MUT#2^ShCTR is 4.251± 0.71670 h  CCP1 ^MUT#2^Sh#1 is 4.972 ± 0.6298 h  CCP1^MUT#2^Sh#2 is 2.417 ± 0.2795 h | 13 clINs from CCP1^WT#2^ShCTR  15 clINs from CCP1^Mut#2^ShCTR  13 clINs from CCP1^Mut#2^Sh#1  15 clINs from CCP1^Mut#2^Sh#2  of 4-5 hvFOs of 3 independents  differentiations |
| Fig. 4K |  | | One-way ANOVA  Ns  (p=0.97421) | DMSO + CCP1^WT#2^ShCTR is 21.89 ± 3.044 μm/h  DMSO+ CCP1^MUT#2^ShCTR is 21.63 ± 3.242 μm/h  ML7 + CCP1^WT#2^Sh#1 is 21.48 ± 2.339 μm/h  ML7 + CCP1^MUT#2^Sh#1 is 23.73± 3.954 μm/h | 14 clINs from CCP1^WT#2^ShCTR+DMSO  10 clINs from CCP1^Mut#2^ShCTR +DMSO  19 clINs from CCP1^WT#2^Sh#1 +ML7  6 clINs from CCP1^Mut#2^Sh#1 +ML7  of 4-5 hvFOs of 3 independents  differentiations |
| Fig. 4L |  | | One-way ANOVA  ** p=0.0074, plus Tukey’s multiple comparisons test  DMSO+ CCP1^MUT#2^ShCTR vs  ML7 +CCP1^MUT#2^Sh#1^:^ *  p=0.0149  DMSO+ CCP1^MUT#2^ShCTR vs  ML7 +CCP1^WT#2^Sh#1^:^ **p=0.0085 | DMSO + CCP1^WT#2^ShCTR is 6.233 ± 0.8134 h  DMSO+ CCP1^MUT#2^ShCTR is 3.25 ± 0.6455 h  ML7 + CCP1^WT#2^Sh#1 is 7.342 ± 0.6753 h  ML7 + CCP1^MUT#2^Sh#1 is 8.063± 0.6852 h | 15 clINs from CCP1^WT#2^ShCTR+DMSO  7 clINs from CCP1^Mut#2^ShCTR +DMSO  19 clINs from CCP1^WT#2^Sh#1 +ML7  6 clINs from CCP1^Mut#2^Sh#1 +ML7  of 4-5 hvFOs of 3 independents  differentiations |
| Fig. 4M |  | | One-way ANOVA  ** p=0.0013, plus Tukey’s multiple comparisons test  DMSO CCP1^WT#2^ShCTR vs  DMSO CCP1^MUT#2^ShCTR^:^ * p=0.0348  DMSO CCP1^MUT#2^ShCTR vs  ML7 CCP1^MUT#2^Sh#1^:^ *  p=0.0254  DMSO CCP1^MUT#2^ShCTR vs  ML7 CCP1^WT#2^Sh#1^:^ ***  p=0.0006 | DMSO + CCP1^WT#2^ShCTR is 1.547 ± 0.1746 event/h  DMSO+ CCP1^MUT#2^ShCTR is 0.7244 ± 0.1263 event/h  ML7 + CCP1^WT#2^Sh#1 is 1.925 ± 0.1785 event/ h  ML7 + CCP1^MUT#2^Sh#1 is 1.796± 0.2106 event/ h | 16 clINs from CCP1^WT#2^ShCTR+DMSO  8 clINs from CCP1^Mut#2^ShCTR +DMSO  19 clINs from CCP1^WT#2^Sh#1 +ML7  6 clINs from CCP1^Mut#2^Sh#1 +ML7  of 4-5 hvFOs of 3 independents  differentiations |
| Fig. 5C |  | | One-way ANOVA;  ***p<0,001, plus Tukey’s multiple comparisons test  CCP1^WT#2^ vs  CCP1^Mut#2^: * (p=0.0147)  CCP1 ^Mut#2^ vs  CCP1^Iso#2^: ***(p=0.0007)  CCP1 ^WT#2^ vs  CCP1^Iso#2^: ns (p=0.00776) | CCP1^WT^ is 1± 0  CCP1^MUT#2^ is 1.936 ± 0.2283  CCP1^Iso#2^ is 0.3224 ± 0.1114 | N=3  differentiations, 6-10 organoids per pooled |
| Fig. 5E |  | | Unpaired t test  ** p<0.001  (p=0.005) | Mouse cINs is 4.4 ± 0.6027  Human cINs is 1.991± 0.2477 | 21 cINs from 3 pool of 4 E13.5 embryos  14 cINs from 3 human fetuses GW 11/13 |
| Fig. 5G |  | | Mann Whitney test  *p<0.05  (p=0.0317) | Mouse cINs is 16.55 ± 4.337  Human clINs is 4.706 ± 1.355 | 101 cINs from 5 independent differentiations of hvFOs  145 cINs from 5 pool of 2-3 E13.3 embryos |
| Fig. 6B |  | | Unpaired t test  **** p<0.0001 | Mean of mouse cINs is 0.565 ±0.06262 events/h  Mean of human cINs is 1.198 ± 0.1241 events/h | 27 cINs from 3 pool of E13.5 mouse embryos  27 cINs from 3 GW11-13 human fetuses |
| Fig. 6C  Fig. 6D |  | | Fisher’s exact test  **** p<0.0001 | Mouse cINs contacts:  Repulsive: 18.52%  Non- repulsive:81.48%  Human cINs contacts:  Repulsive:74.07%  Non- repulsive:25.93% | 27 cINs from 3 pool of E13.5 mouse embryos  27 cINs from 3 GW11-13 human fetuses |
| Fig. 6F |  | | Unpaired t test  **p<0.05  (p=0.0048) | CCP1^dead^ is 1.51± 0.1658 events/h  CCP1^wt^ is 2.189± 0.1549 events/h | 23 cINs from 5-6 embryos for CCP1^wt^  27 cINs from 5-6 embryos for CCP1^dead^ |
| Fig. 6G |  | | Unpaired t test  *p<0.05  (p=0.0487) | CCP1^dead^ is 0.352± 0.05334 events/h  CCP1^wt^ is 0.5177± 0.06303 events/h | 18 cINs from 5-6 embryos for CCP1^wt^  20 cINs from 5-6 embryos for CCP1dead |
| Fig. 6H |  | | Fisher’s exact test  Ns, p=0.1440 | CCP1^dead^ contacts:  Repulsive: 15%  Non- repulsive:85%  CCP1^wt^ contacts:  Repulsive:39%  Non- repulsive:61% | 18 cINs from 5-6 embryos for CCP1^wt^  20 cINs from 5-6 embryos for CCP1^dead^ |
| Fig. 6I  Fig.  6J | Left panel  Center panel  Right panel | | Molecular dynamics (start pallium)  Molecular dynamics (start CP basis) | N=400: 70% of surrogates above CP basis  N=320: instantaneous snapshot of positions | INs probability density averaged over 250 configurations (cf) , spacing 5000 time steps (t.s)  cINs surrogates positions after 3000*5000 t.s.  Avg. distance to CP top line (last 450 cf – spacing 5000 t.s.) |
| Fig. 6 M |  | | Ordinary two-way ANOVA followed by Tukey’s multiple comparisons test.  Two-way ANOVA:  Interaction (bin between human and mouse): ***p<0.001  p = 0.0002  bin 1: ns, p = 0.7972  bin 2: ns, p = 0.1585  bin 3: ns, p = 0.3823  bin 4: ns, p = 0.5089  bin5: ns, p = 0.0803  bin6: ns, p = 0.3321  bin 7: *p = 0.0187  bin 8: ns, p = 0.1135  bin 9: ns, p = 0.5522  bin 10: *p = 0.0148 | bin 1: human= 2,14 ± 0,68; mouse=0,17± 0,17  bin2: human= 8,20± 2,46; mouse=2,522 ±1,537  bin 3: human= 9,96 ± 2,86; mouse  =5,887±0,742  bin 4: human = 6,38 ±1,35; mouse=9,783  ± 0,832  bin 5: human=11,64 ±2,25; mouse=18,323  ±0,470  bin 6: human=8,33 ±1,12; mouse=12,694  ±1,622  bin 7: human= 15,24  ±1,98; mouse= 6,756  ± 2,033  bin 8: human= 13,23  ± 1,57; mouse= 7,044  ±1,237  bin 9: human=.11,79  ± 1,55; mouse=14,976  ± 5,326  bin 10: human= 13,09  ± 1,94; mouse=  ± | 4 E13,5 embryos of different Swiss female/  4 GW11/13 fetuses |
| Fig. 6 O |  | | Ordinary two-way ANOVA followed by Tukey’s multiple comparisons test.  Two-way ANOVA:  Interaction (bin between: LoxStoLox-CCP1^wt^ and LoxStoLox-CCP1^dead^): ***p<0.001  bin 1: ns, p > 0.9999  bin 2: ns, p = 0.2825  bin 3: ****, p < 0.0001  bin 4: ns, p > 0.9999  bin 5: ns, p > 0.9999  bin 6: ns, p > 0.9999  bin 7: ns, p > 0.9999  bin 8: ns, p > 0.9999  bin 9: ns, p > 0.9999  bin 10: ns, p = 0.9997  bin 11: ns, p > 0.9999  bin 12: ns, p > 0.9999  bin 13: ns, p = 0.9997  bin 14: ns, p = 0.9998  bin 15: ns, p > 0.9999 | bin 1: CCP1 wt= 3,26  ±1,57; CCP1 dead=0± 0  bin2: CCP1 wt= 31,73 ± 5,99; CCP1 dead =20,37 ±6,68  bin 3: CCP1 wt= 15,69 ± 5,03; CCP1 dead=48,15±9,8  bin 4: CCP1 wt = 12,73 ± 5,42; CCP1 dead=15,74  ± 7,91  bin 5: CCP1wt=3,41 ±1,97; CCP1 dead =0,0 ±0,0  bin 6: CCP1wt=3,50 ±1,55; CCP1 dead =0,0 ±0,0  bin 7: CCP1wt=2,43 ±1,53; CCP1 dead =0,0 ±0,0  bin 8: CCP1wt=3,51 ±1,68; CCP1 dead =0,0 ±0,0  bin 9: CCP1wt=1,96 ±1,97; CCP1 dead =0,0 ±0,0  bin 10: CCP1wt=4,01 ±1,23; CCP1 dead =0,0 ±0,0  bin 11: CCP1wt=1,55 ±1,05; CCP1 dead =0,0 ±0,0  bin12: CCP1wt=1,77 ±0,75; CCP1 dead =0,0 ±0,0  bin13: CCP1wt=4,01 ±1,23; CCP1 dead =0,0 ±0,0  bin14: CCP1wt=5,46 ±3,73; CCP1 dead =9,26 ±4,9  bin15: CCP1wt=5 ±1,82; CCP1 dead =6,48 ±3,34 | 5 E17.5 embryos Dlx5/6:cre-GFP/ LoxStoLox-CCP1^wt^  3 E17.5 embryos Dlx5/6:cre-GFP/ LoxStoLox-CCP1^dead^ |
| Fig. 6P |  | | Mann Whitney test  *p<0.05  (p=0.0357) | CCP1^dead^ is 15.74± 7.911 CCP1^wt^ is 38.12 ± 4.1 | 5 E17.5 embryos Dlx5/6:cre-GFP/ LoxStoLox-CCP1^wt^  3 E17.5 embryos Dlx5/6:cre-GFP/ LoxStoLox-CCP1^dead^ |
| Fig. 6Q |  | | Mann Whitney test  *p<0.05  (p=0.0358) | CCP1^dead^ is 4± 1 CCP1^wt^ is 10.41 ± 1.2 | 5 E17.5 embryos Dlx5/6:cre-GFP/ LoxStoLox-CCP1^wt^  3 E17.5 embryos Dlx5/6:cre-GFP/ LoxStoLox-CCP1^dead^ |
| Fig. S1D |  | | Unpaired t test  **** p<0.0001 | Mouse is 11.89 ± 0.48  Human is 37.71± 2.794 | 18 cINs from 3 E13.5 mouse embryos  15 cINs from 5 GW11-13 human fetuses |
| Fig. S1E |  | | Unpaired t test  **** p<0.0001 | Mouse is 20 ± 0.6329 microns  Human is 30.76± 1.294 microns | 35 cINs from 4-5 E13.5 mouse embryos  34 cINs from 5 GW11-12 human fetuses |
| Fig. S1F | Secondary neurite  Tertiary neurite | | One-way ANOVA  ***p<0.0001, plus Tukey’s multiple comparisons test.  Human cINS vs Mouse cINS  *** p=0.007  Mouse cINs vs  Human cINS  **** p<0.0001 | Mouse is 0.3696 ± 0.011  Human is 0.4794 ± 0.028  Mouse is 0.1339 ± 0.008  Human is 0.2126± 0.01384 | 22 cINs from 4-5 E13.5 mouse embryos  24 cINs from 5 GW11-12 human fetuses  22 cINs from 4-5 E13.5 mouse embryos  24 cINs from 5 GW11-12 human fetuses |
| Fig. S2F |  | | Mann Whitney test **p=0.0095 | Mouse is 1 ± 0.04912 a.u.  Human is 2.292± 0.1215 a.u. | 4 E13.5 mouse embryos  6 GW11-12 human fetuses |
| Fig. S2G |  | | Mann Whitney test **p=0.0095 | Mouse is 1 ± 0.02823 a.u.  Human is 2.642± 0.3594 a.u. | 4 E13.5 mouse embryos  6 GW11-12 human fetuses |
| Fig. S2I |  | | One-way ANOVA  Ns; p=0.5177 | shCTR is 0.8674 ± 0.088 event/h  shCCP1 #1 is 0.7281± 0.1141 event/h  shCCP1 #2 is 0.8924 ± 0.1006 µm/h | 25 cINs from 3-4 fetuses GW11-13 embryos for shCTR  shCCP1 #1  17 cINs from 3-4 fetuses GW11-13 embryos for shCCP1#1  10 cINs from 3 fetuses GW11-13 embryos for  shCCP1 #2 |
| Fig. S2J |  | | Ordinary one-way ANOVA followed by Tukey’s multiple comparisons test.  ShCTR vs  ShCCP1#1^:^ *  (p=0.0449)  ShCTR vs  ShCCP1#2^:^ *  (p=0.0318) | ShCTR is 6.1212 ± 0.3449  ShCCP1#1 is 4.851 ± 0.3821  ShCCP1#2 is 4.726 ± 0.4283 | 25 cINs from 3-4 fetuses GW11-13 embryos for shCTR  shCCP1 #1  17 cINs from 3-4 fetuses GW11-13 embryos for shCCP1#1  10 cINs from 3 fetuses GW11-13 embryos for  shCCP1 #2 |
| Fig. S3A | EMX2  EMX1  PAX6  TBR2 | | Shapiro-Wilk for all Unpaired t-test: ** p=0.0025  Unpaired Mann Whitney: ** p=0.0087  Unpaired t-test: * p=0.0397  Unpaired t-test: * p=0.0286 | hdFOs is 0.8621 ± 0.194  hvFOs is 0.0838 ± 0.016  hdFOs is 0.155 ± 1.053  hvFOs is 0.0005 ± 0.0004  hdFOs is 1.200 ± 0.236  hvFOs is 0.586 ± 0.108  hdFOs is 1.887 ± 0.707  hvFOs is 0.078 ± 0.079 | 8 to 12 hdFOs for each of the 3 differentiations  8 to 12 hvFOs for each of the 3 differentiations  Technical triplicates was additionally performed with its CT averaged |
| Fig. S3B | NKX2.1  DLX2  DLX5 | | Shapiro-Wilk for all Unpaired Mann Whitney: ** p=0.0022  Unpaired T-test: ***, p=0.001  Unpaired Mann Whitney: ** p=0.0022 | hdFOs is 0.003 ± 0.013  hvFOs is 1.212 ± 0.763  hdFOs is 0.120 ± 0.029  hvFOs is 1.085 ± 0.207  hdFOs is 0.123 ± 0.053  hvFOs is 0.817 ± 0.494 | 8 to 12 hdFOs for each of the 3 differentiations  8 to 12 hvFOs for each of the 3 differentiations  Technical triplicates was additionally performed with its CT averaged |
| Fig. S3H |  | | Ordinary one-way ANOVA followed by Tukey’s multiple comparisons test.  Ns, no significative. | CCP1^WT#1^ is 22.06 ± 1.76 µm/h  CCP1^WT#2^ is 25.76 ± 1.38 µm/h  CCP1^WT#3^ is 24.51 ± 2.71 µm/h  CCP1^MUT#1^ is 17.08 ± 2.61 µm/h  CCP1^MUT#2^ is 16.22 ± 1.48 µm/h  CCP1^MUT#3^ is 18.63 ± 2.55 µm/h | 27/28 clINs from 3-4 hvFOs CCP1^WT#1^ from 3 differentiation  7 clINs from 5 hvFOs CCP1^WT#2^ from 3 differentiation  6/7 clINs from 4-7 hvFOs CCP1^WT#3^ from 3 differentiation  6 clINs from 3-4 hvFOs CCP1^MUT#1^ from 3 differentiation  11/12 clINs from 4-5 hvFOs CCP1^MUT#2^ from 3 differentiation  5 clINs from 3 hvFOs CCP1^MUT#3^ from 3 differentiation |
| Fig. S3I |  | | Ordinary one-way ANOVA followed by Tukey’s multiple comparisons test.  CCP1^WT#1^ vs CCP1^MUT#1^: *, p=0.0368  CCP1^WT#2^ vs CCP1^MUT#1^: ***, p=0.0003  CCP1^WT#3^ vs CCP1^MUT#1^: **,  p=0.0019  CCP1^WT#1^ vs CCP1^MUT#2^: **, p=0.0013  CCP1^WT#2^ vs CCP1^MUT#2^: ****,  p<0.0001  CCP1^WT#3^ vs CCP1^MUT#2^: ***,  p=0.0001  CCP1^WT#1^ vs CCP1^MUT#3^: ns  CCP1^WT#2^ vs CCP1^MUT#3^: **,  P=0.0016  CCP1^WT#3^ vs CCP1^MUT#3^: **,  p=0.0085 | CCP1^WT#1^ is 21.27 ± 1.06 µm  CCP1^WT#2^ is 27.02 ± 2.03 µm  CCP1^WT#3^ is 25.54 ± 2.28 µm  CCP1^MUT#1^ is 14.61 ± 1.05 µm  CCP1^MUT#2^ is 14.06 ± 0.38 µm  CCP1^MUT#3^ is 15.38 ± 1.83 µm | 27/28 clINs from 3-4 hvFOs CCP1^WT#1^ from 3 differentiation  7 clINs from 5 hvFOs CCP1^WT#2^ from 3 differentiation  6/7 clINs from 4-7 hvFOs CCP1^WT#3^ from 3 differentiation  6 clINs from 3-4 hvFOs CCP1^MUT#1^ from 3 differentiation  11/12 clINs from 4-5 hvFOs CCP1^MUT#2^ from 3 differentiation  5 clINs from 3 hvFOs CCP1^MUT#3^ from 3 differentiation |
| Fig. S3J |  | | Ordinary one-way ANOVA followed by Tukey’s multiple comparisons test.  CCP1^WT#1^ vs CCP1^WT#2^: *, p=0.143  CCP1^WT#1^ vs CCP1^WT#3^: ****,  p<0.0001  CCP1^WT#2^ vs CCP1^MUT#1^: ***, p=0.0001  CCP1^WT#3^ vs CCP1^MUT#1^: ****,  p<0.0001  CCP1^WT#1^ vs CCP1^MUT#2^: *, p=0.0136  CCP1^WT#2^ vs CCP1^MUT#2^: ****,  p<0.0001  CCP1^WT#3^ vs CCP1^MUT#2^: ****,  p<0.0001  CCP1^WT#2^ vs CCP1^MUT#3^: **, p=0.0016  CCP1^WT#3^ vs CCP1^MUT#3^: ****,  p<0.0001 | CCP1^WT#1^ is 4.97 ± 0.33 h  CCP1^WT#2^ is 7.04 ± 0.41 h  CCP1^WT#3^ is 8.14 ± 0.85 h  CCP1^MUT#1^ is 3.18 ± 0.28 h  CCP1^MUT#2^ is 3.27 ± 0.26 h  CCP1^MUT#3^ is 3.60 ± 0.30 h | 27/28 clINs from 3-4 hvFOs CCP1^WT#1^ from 3 differentiation  7 clINs from 5 hvFOs CCP1^WT#2^ from 3 differentiation  6/7 clINs from 4-7 hvFOs CCP1^WT#3^ from 3 differentiation  6 clINs from 3-4 hvFOs CCP1^MUT#1^ from 3 differentiation  11/12 clINs from 4-5 hvFOs CCP1^MUT#2^ from 3 differentiation  5 clINs from 3 hvFOs CCP1^MUT#3^ from 3 differentiation |
| Fig. S3K |  | | Unpaired T-test: ***, p=0.0003 | CCP1^WT#1&WT#2^ is 1.4 ± 0.14 CCP1^MUT#1&MUT#2^ is 0.73 ± 0.09 | 15 hvFOs for at least 3 independent differentiations. |
| Fig. S3L |  | | Unpaired T-test= ns | CCP1^WT#1&WT#2^ is 111.67 ± 13  CCP1^MUT#1&MUT#2^ is 94.3 ± 8.84 | 8 hvFOs for at least 3 independent differentiations |
| Fig. S3M | Secondary neurite  Tertiary neurite | | Ordinary one-way ANOVA:  ****, p<0.0001  Ns | CCP1^WT#1&WT#2^ is 4.93 ± 0.28  CCP1^MUT#1&MUT#2^ is 2.53 ± 0.24  CCP1^WT#1&WT#2^ is 2.07 ± 0.23  CCP1^MUT#1&MUT#2^ is 1.53 ± 0.22 | 15 hvFOs for at least 3 independent differentiations |
| Fig. S4D |  | | Kruskal-Wallis test  Followed by Dunn’s multiple comparision test.  Ns., no significative.  CCP1^WT#2^ vs. CCP1^WT#1^ p=0.8635  CCP1^WT#1^vs  CCP1^Mut#1-#3^ p>0.9999 | CCP1^WT#2^ is 1.001 ± 0.0029 (a.u.)  CCP1^WT#3^ is 0.6091 ± 0.07718 (a.u.)  CCP1^MUT#1^ is 1.174 ± 0.128 (a.u.)  CCP1^MUT#2^ is 2.596± 0.75(a.u.)  CCP1^MUT#3^ is 0.7905 ± 0.05 (a.u.) | 8 to 12 hdFOs for each of the 5 differentiations  Technical triplicates were additionally performed with its CT averaged |
| Fig. S4G |  | | Unpaired t test  Ns, p=0.7564 | CCP1 WT mouse is 1.476± 0.1826  CCP1 cKO mouse is 1.307 ± 0.1626 | 4 embryos of E13.5 of each genotype of different females. |
| Fig. S4I |  | | Unpaired t test  Ns, p=0.2087 | CCP1 WT mouse is 2.154 ± 0.2254  CCP1 cKO mouse is 1.952 ± 0.4184 | 4 embryos of E13.5 of each genotype of different females. |
| Fig. S4J |  | | Ordinary one-way ANOVA followed by Tukey’s multiple comparisons test.  shCTR vs shDCX#1=*** p=0.0002  shCTR vs shDCX#2***  p=0.0006 | shCTR is 100%±0  shDCX#1 is 7.667 ±1.453  shDCX#2 is 20.33±5.840 | 3 experiments independents of 3 differents HEK co transfections. |
| Fig. S5B | Resting potential  MAX OXPHOS  MAX ETC | | Ordinary one-way ANOVA  .  Ns  Ns  Ns | CCP1^WT#2^ is 5.83 ± 0.93 pmols O2/mL / million cells  CCP1^MUT#2^ is 5.50 ± 1.02 pmols O2/mL / million cells  CCP1^Iso#2^ is 6.89 ± 0.91 pmols O2/mL / million cells  CCP1^WT#2^ is 17.22 ± 1.94 pmols O2/mL / million cells  CCP1^MUT#2^ is 20.01 ± 5.91 pmols O2/mL / million cells  CCP1^Iso#2^ is 22.61 ± 4.15 pmols O2/mL / million cells  CCP1^WT#2^ is 25.46 ± 1.48 pmols O2/mL / million cells  CCP1^MUT#2^ is 27.03 ± 6.62 pmols O2/mL / million cells  CCP1^Iso#2^ is 33.62 ± 6.76 pmols O2/mL / million cells | For 2 biological replicates 30 hvFOs were combined per differentiation, the 3^rd^ replicate was computed with 35 hvFOs from the 3^rd^ differentiation |
| Fig. S5D |  | | One way ANOVA  Ns | CCP1^WT#2^ is 1.00 ± 0.13  CCP1^MUT#2^ is 0.45 ± 0.17  CCP1^Iso#2^ is 0.73 ± 0.40 | For all conditions, 5 to 8 hvFOs were pooled for each of the 3 differentiations  (until S5G) |
| Fig. S5E |  | | One way ANOVA  Ns | CCP1^WT#2^ is 1.00 ± 0.30  CCP1^MUT#2^ is 0.18 ± 0.01  CCP1^Iso#2^ is 0.64 ± 0.45 |  |
| Fig. S5F |  | | One way ANOVA  Ns | CCP1^WT#2^ is 1.00 ± 0.33  CCP1^MUT#2^ is 0.36 ± 0.22  CCP1^Iso#2^ is 0.57 ± 0.27 |  |
| Fig. S5G |  | | One way ANOVA  Ns | CCP1^WT#2^ is 1.00 ± 0.32  CCP1^MUT#2^ is 0.40 ± 0.05  CCP1^Iso#2^ is 0.52 ± 0.08 |  |
| Fig. S5J |  | | Shapiro-Wilk, Unpaired t-test  *, p=0.04 | CCP1^WT^ is 42.2 ± 3.5  CCP1^MUT#2^ is 31.8 ± 3.2 | CCP1^WT#2^: 18 cells from explants of 4/5 hvFOs from 3 differentiation  CCP1^MUT#2^: 12 cells from explants of 4/5 hvFOs from 3 differentiation |
| Fig. S5K |  | | Shapiro-Wilk, Unpaired t-test  Ns, p=0.08 | CCP1^WT^ is 1.1 ± 0.1 (μm^2^)  CCP1^MUT#2^ is 1.4 ± 0.1 (μm^2^) | CCP1^WT#2^: 18 cells from explants of 4/5 hvFOs from 3 differentiation  CCP1^MUT#2^: 12 cells from explants of 4/5 hvFOs from 3 differentiation |
| Fig. S5L |  | | Shapiro-Wilk, Mann-Whitney test  Ns, p=0.7 | CCP1^WT^ is 79.6 ± 8.4 (event/frame)  CCP1^MUT#2^ is 70.5 ± 6.2 (event/frame) | CCP1^WT#2^: 18 cells from explants of 4/5 hvFOs from 3 differentiation  CCP1^MUT#2^: 12 cells from explants of 4/5 hvFOs from 3 differentiation |
| Fig. S5M |  | | Shapiro-Wilk,  Unpaired Mann-Whitney test  Ns, p=0.9 | CCP1^WT^ is 80.4 ± 7.4 (event/frame)  CCP1^MUT#2^ is 75.1 ± 6.7 (event/frame) | CCP1^WT#2^: 18 cells from explants of 4/5 hvFOs from 3 differentiation  CCP1^MUT#2^: 12 cells from explants of 4/5 hvFOs from 3 differentiation |
| Fig. S5N | %TMA/%cell area soma : Pre-nucleokinesis  %TMA/%cell area soma : Pre-nucleokinesis  %TMA/%cell area soma : Pre-nucleokinesis  %TMA/%cell area soma : Pre-nucleokinesis  %TMA/%cell area soma : Pre-nucleokinesis  %TMA/%cell area soma : Pre-nucleokinesis | | Two way ANOVA : Ns  Ns  Ns  Two way ANOVA : Ns  Ns  Ns | CCP1^WT#2^ is 1.09 ± 0.20  CCP1^MUT#2^ is 1.20 ± 0.02  CCP1^WT#2^ is 1.46 ± 0.11  CCP1^MUT#2^ is 1.44 ± 0.09  CCP1^WT#2^ is 1.17 ± 0.13  CCP1^MUT#2^ is 1.16 ± 0.027  CCP1^WT#2^ is 0.89 ± 0.11  CCP1^MUT#2^ is 0.73 ± 0.07  CCP1^WT#2^ is 0.72 ± 0.05  CCP1^MUT#2^ is 0.49 ± 0.03  CCP1^WT#2^ is 0.82 ± 0.13  CCP1^MUT#2^ is 1.02 ± 0.64 | CCP1^WT#2^: 4 cells from explants of 4 hvFOs from 4 differentiation  CCP1^MUT#2^: 3 cells from explants of 3 hvFOs from 3 differentiation |
| Fig. S6C |  | | Unpaired t testNs  Ns, p=0.6872 | CCP1^dead^ is 21.21± 1.818 µm/h  CCP1^wt^ is 20.09± 2.074µm/h | 21 cINs from 5-6 embryos for CCP1^wt^  25 cINs from 5-6 embryos for CCP1^dead^ |
| Fig. S6D |  | | Unpaired t test  Ns, p=0.2742Ns | CCP1^dead^ is 1.061± 0.135 events/h  CCP1^wt^ is 0.8741 ± 0.08354 events/h | 21 cINs from 5-6 embryos for CCP1^wt^  25 cINs from 5-6 embryos for CCP1^dead^ |
| Fig. S6E |  | | Unpaired t test  ***p<0.0001 | CCP1^dead^ is 3.2± 0.5270 h  CCP1^wt^ is 7.438 ± 0.72027 h | 20 cINs from 5-6 embryos for CCP1^wt^  25 cINs from 5-6 embryos for CCP1^dead^ |
| Fig. S6F | |  | Unpaired t test  ** p<0.01**  p=0.0015 | CCP1^dead^ is 62.23± 3.514 µm  CCP1^wt^ is 80.80 ± ± 3.703 µm | 12 cINs from 5-6 embryos for CCP1^wt^  18 cINs from 5-6 embryos for CCP1^dead^ |
| Fig. S6G | |  | Unpaired t test  *** p<0.0001*** | CCP1^dead^ is 26.18± 1.768 µm  CCP1^wt^ is 49.47 ± 4.066 µm | 12 cINs from 5-6 embryos for CCP1^wt^  18 cINs from 5-6 embryos for CCP1^dead^ |
| Fig. S6H | |  | Unpaired t test  *** p<0.0001*** | CCP1^dead^ is 2.11 ± 0.2271  CCP1^wt^ is 5.5 ± 0.3794 | 12 cINs from 5-6 embryos for CCP1^wt^  18 cINs from 5-6 embryos for CCP1^dead^  \ |
| Fig. S6M | |  | Ordinary two-way ANOVA followed by Tukey’s multiple comparisons test.  Two-way ANOVA:  Interaction (bin between: LoxStoLox-CCP1^wt^ and LoxStoLox-CCP1^dead^): Ns, not significative.  (p=0.3980) | bin 1: CCP1 wt= 2,64  ±1,24; CCP1 dead=1,84± 1  bin2: CCP1 wt= 7,23 ± 3,19; CCP1 dead =7,09 ±1,98  bin 3: CCP1 wt= 7,56 ± 1,55; CCP1 dead=10,88±3,72  bin 4: CCP1 wt = 5,33 ± 0,96; CCP1 dead=4,5  ± 0,71  bin 5: CCP1wt=5,53 ±1,97; CCP1 dead =0,0 ±0,0  bin 6: CCP1wt=5,13 ±0,48; CCP1 dead =4,5 ±1,4  bin 7: CCP1wt=4 ±0,13; CCP1 dead =3,99 ±0,45  bin 8: CCP1wt=4,77 ±0,71; CCP1 dead =4,27±1,37  bin 9: CCP1wt=5,58 ±1,25; CCP1 dead =5,87 ±2,4  bin 10: CCP1wt=7,96 ±1,43; CCP1 dead =6,97 ±0,43  bin 11: CCP1wt=8,21 ±1,30; CCP1 dead =7,45 ±0,79  bin12: CCP1wt=8,26 ±1,80; CCP1 dead =7,13 ±1,14  bin 13: CCP1wt=8,43 ±1,80; CCP1 dead =7,16 ±1,48  bin14: CCP1wt=12,46 ±3,19; CCP1 dead =12,45 ±4,99  bin15: CCP1wt=6,93 ±2,89; CCP1 dead =10,17 ± 2,55 | 5 E17.5 embryos Dlx5/6:cre-GFP/ LoxStoLox-CCP1^wt^  3 E17.5 embryos Dlx5/6:cre-GFP/ LoxStoLox-CCP1^dead^ |
| Fig. S8B | |  | Ordinary One-way ANOVA (***, p<0.0001) followed by Tukey’s multiple comparisons test.  cINs Mouse vs cINs ferret: ***(p<0.0001)  cINS Mouse vs cINS human:  *** (p<0.0001)  cINs Human vs cINs ferret: ns,  p=0.069 | Mouse is 44.11 ± 3.129 µm/h  Human is 16.82± 2.497 µm/h  Ferret is 26.71 ± 1.792 µm/h | 29 mouse cINs from 3 independent pool of 3-4 E13.5 embryos  36 ferret E.36 cINs from 3 embryos  12 human cINs from 3 GW12 embryos |
| Fig. S8C | |  | Ordinary One-way ANOVA (***, p<0.0001) followed by Tukey’s multiple comparisons test.  cINs Mouse vs cINs ferret: ***(p<0.0001)  cINS Mouse vs cINS human:  *** (p<0.0001)  cINs Human vs cINs ferret: ns,  p=0.5965 | Mouse is 1.911± 0.1186 events/h  Human is 0.7125 ± 0.1262 events/h  Ferret is 0.8850 ± 0.007 events/h | 29 mouse cINs from 3 independent pool of 3-4 E13.5 embryos  36 ferret E.36 cINs from 3 embryos  12 human cINs from 3 GW12 embryos |
| Fig. S8D | |  | Ordinary One-way ANOVA (***, p<0.0001) followed by Tukey’s multiple comparisons test.  cINs Mouse vs cINs ferret: ***(p<0.0001)  cINS Mouse vs cINS human:  *** (p<0.0001)  cINs Human vs cINs ferret: *** (p<0.0001) | Mouse is 1.379 ± 0.1972 h  Human is 8.865 ± 0.4587 h  Ferret is 4.966 ± 0.3647 h | 29 mouse cINs from 3 independent pool of 3-4 E13.5 embryos  36 ferret E.36 cINs from 3 embryos  13 human cINs from 3 GW12 embryos |
| Fig. S8E | |  | Ordinary One-way ANOVA (***, p<0.0001) followed by Tukey’s multiple comparisons test.  cINs Mouse vs cINs ferret: **(p=0.0014)  cINS Mouse vs cINS human:  *** (p<0.0001)  cINs Human vs cINs ferret: ns,  p=0.1307 | Mouse is 1.506± 0.1199 events /h  Human is 3.14 ± 0.3610 events /h  Ferret is 2.481 ± 0.2164 events /h | 29 mouse cINs from 3 independent pool of 3-4 E13.5 embryos  28ferret E.36 cINs from 3 embryos  13 human cINs from 3 GW12 embryos |
| Fig. S8F | |  | Ordinary two-way ANOVA followed by Tukey’s multiple comparisons test.  Ferret vs mouse  10: **** p<0.001  20: **** p<0.001  30: ** p = 0.0011  40: ns  50: *, p=0.0276  60: ns  70: ns  80: ns  90: ns  100: ns  110: ns | 10: Ferret 3.048±0.234  Mouse is 0±0.0116; human is 2.714±0.$244  20: Ferret 3.857±0.354  mouse is 2.102±0.323, human is 4.5±0.359  30: Ferret 4.748±0.393  mouse is 2.158±0.289, human is 4.429 ±0.359  40: Ferret 3.095±0.308  Mouse is 1.684±0.217; human is 4.429±0.343  50: Ferret 2.714±0.269  Mouse is 1.421±0.207; human is 4.571±0.359  60: Ferret 1.619±0.253  Mouse is 1.316±0.203; human is 3.929±0.486  70: Ferret 1.19±0.245  Mouse is 0.737±0.185; human is 4.071±0.569  80: Ferret 0.905±0.194  Mouse is 0.526±0.160; human is 3±0.469  90: Ferret 0.8±0.192  Mouse is 0.368±0.191; human is 2.429±0.488  100: Ferret 0.619±0.176  Mouse is 0.020±0.011; human is 2.022±0.144  110: Ferret 0.429±0.019  Mouse is 0±0.0; human is 1.429±0.402 | 19 mouse cINs from 3 independent pool of 3-4 E13.5 embryos  21ferret E.36 cINs from 3 embryos  14 human cINs from 3 GW12 embryos |
| Fig. S8G | |  | Ordinary One-way ANOVA (***, p<0.0001) followed by Tukey’s multiple comparisons test.  cINs Mouse vs cINs ferret: ***(p<0.0001)  cINS Mouse vs cINS human:  *** (p<0.0001)  cINs Human vs cINs ferret:  *** (p<0.0001) | Mouse is 11.89 ± 0.4830  Human is 37.71 ± 2.794  Ferret is 22.29 ± 1.491 | 19 mouse cINs from 3 independent pool of 3-4 E13.5 embryos  21ferret E.36 cINs from 3 embryos  14 human cINs from 3 GW12 embryos |
| Fig. S8H | |  | Ordinary One-way ANOVA (**, p=0.0018) followed by Tukey’s multiple comparisons test.  cINs Mouse vs cINs ferret: ns, p=0.7142  cINS Mouse vs cINS human:  *(p=0.0134)  cINs Human vs cINs ferret: ns, p=0.1828 | Mouse is 0.5655 ± 0.006 events /h  Human is 0.9842 ± 0.09521 events /h  Ferret is 0.7090 ± 0.089 events /h | 27 mouse cINs from 3 independent pool of 3-4 E13.5 embryos  27 ferret E.36 cINs from 3 embryos  27 human cINs from 3 GW11-12 embryos |
| Fig. S8K | |  | Ordinary two-way ANOVA followed by Tukey’s multiple comparisons test.  Two-way ANOVA:  Interaction: ***p<0.001  p = 0.0002  bin 1: ns  bin 2: ns  bin 3: * ferret vs mouse (p=0.0421)  bin 4: ns  bin5: * ferret vs mouse (p=0.0137)  bin6: ns  bin 7: * human vs mouse (p = 0.0187)  bin 8: ns,  bin 9: ns,  bin 10: *human vs mouse (p = 0.0148)  * ferret vs mouse (p=0.0144) | bin 1: human= 2.14 ± 0.68; mouse=0.17± 0.17  ferret=5.632±0.61  bin2: human= 8.20± 2.46; mouse=2.522 ±1.537; ferret= 9.314±1.433  bin 3: human= 9.96 ± 2.86; mouse  =5.887±0.742;  Ferret=14.014±0.661  bin 4: human = 6.38 ±1.35; mouse=9.783  ± 0.832; ferret=10.018±1.293  bin 5: human=11.64 ±2.25; mouse=18.323  ±0.470 ; ferret= 8.7891±4.04  bin 6: human=8.33 ±1.12; mouse=12.694  ±1.622; ferret=9.193± 3.953  bin 7: human= 15.24  ±1.98; mouse= 6.756  ± 2.033; ferret= 4.651±0.776  bin 8: human= 13.23  ± 1.57; mouse= 7.044  ±1.237; ferret=12.760±4.14  bin 9: human=.11.79  ± 1.55; mouse=14.976  ± 5.326; ferret=13.274±1.380  bin 10: human= 13.09  ± 1.94; mouse= 21.841 ± 3.549; ferret=12.364 ±4.0484 | 4 E13,5 embryos of different Swiss female/  4 GW11/13 fetuses |
| Fig. S8M | |  | Unpaired t test  *<0.05 (p=0.0230) | Mouse is 1± 0.1188  Ferret is 2.147 ± 0.3592 | 4 pools of 2-3 mice E13.5 MGE;  4 MGE extracts coming from 4 embryos ferret E36. |
| Fig. S8O | |  | Unpaired t test  **<0.001 (p=0.0033) | Mouse is 1± 0.03715  Ferret is 0.4577 ± 0.167 | 4 pools of 2-3 mice E13.5 MGE;  4 MGE extracts coming from 4 embryos ferret E36. |

**Supplemental figures**

***Figure S1. Migratory dynamics of human and mouse cINs in situ***

**A.** UMAP of DLX5/6+ post-mitotic cells integrating mice and human GEs and subpallial datasets and showing expression of representative GE gene markers : CGE (*NR2F1, NR2F2*), LGE (*MEIS2, FOXP1, FOXP2*), MGE (*LHX6, MAF, NKX2-1*). **B**, **C**. Representative images of the distribution of migrating Dlx5/6-GFP expressing cINs in organotypic forebrain slices from GW12 human fetuses (B), and E13.5 mouse embryos (C), CP is cortical plate and t the time computed in minutes. Slices were transduced with a Dlx5/6-GFP lentiviral vector and recorded by time-lapse microscopy for 24 h. Scale bars, 50 μm**. D.** Branching complexity of cINs migrating from MGE explant cultures of **E13.5** mouse **or GW12 human forebrain.** **E, F**. Violin plots of amplitude of nucleokinesis (E) and normalized maximal secondary (II) and tertiary (III) neurite length (F) of mouse and human cINs migrating from MGE explants. Data represent 15 to 30 cells from distinct mouse embryos and human fetuses (4 to 5) ; *** p<0.001, ****p< 0.0001, unpaired t test.

***Figure S2. CCP1 and polyglutamylated tubulin expression in developing cINs across species***

**A**. Over-representation analysis (ORA) of genes contributing to the IPA function associated with cell migration pathways. The biological process (BP) GO term cytoskeleton-related pathways are shown as a dot plot, with dot size indicating the number of contributing genes and color representing the adjusted p-value (q value). **B.** Representative images of human cINs (GW12) co-expressing CCP1 (red) and βIII-tubulin (gray). Scale bar, 10 μm. **C, D.** Representative images of migrating cINs from E13.5 mouse (C) or GW11-12 human (D) MGE explant cultures. Explants were transduced with Dlx5/6-eGFP lentivirus (green) and immunolabeled for polyglutamylated tubulin (PolyE, magenta) and βIII-tubulin (gray). Scale bars, 20 μm. **E,** **F**. qRT-PCR analysis of *TTL1* (E) *or TTL7* (F) expression in human (GW11-12; n=5) and mouse (E13.5; n=5) MGE samples, normalized to mouse expression (set to 1). **p<0.001, Mann Whitney test, **G.** Experimental workflow for lentiviral transduction and live-cell imaging. Migrating cINs in human MGE explant cultures were transduced with Dlx5/6-eGFP lentiviral vectors expressing either control shRNA (shCTR) or CCP1-targeting shRNAs (shCCP1#1 and shCCP1#2). **H, I**. Quantitative analysis of cIN migration parameters. Violin plots show nucleokinesis frequency (H), and maximum number of secondary neurites per cell (I) (n=15-30 cells from 3-4 GW11-12 human embryos; one-way ANOVA with Tukey's multiple-comparison test; *p < 0.05, **p< 0.01).

***Figure S3. Human forebrain assembloid as a model to study the role of CCP1 in cortical interneuron migration***

**A-B**. RT-qPCR analyses of marker gene expression level in 30 days old human dorsal forebrain organoids (hdFOs) (*EMX1*, *EMX2*, *PAX6*, *TBR2*) (A), and human ventral forebrain organoids (hvFOS) (*DLX2*, *DLX5*, *NKX2.1*) (B) forebrain identities (n=8-12 organoids per extraction, 3 independent differentiations per cell line; unpaired t-test or Mann-Whitney). *p<0.05, **p<0.01, ***p<0.001; ns, not significant. **C.** UMAP of integrated CCP1^WT#2^, CCP1^Mut#2^, and CCP1^Iso#2^ scRNAseq datasets, showing expression of representative marker genes used for the identification of the following cell populations: dividing progenitors (*MKI67*), progenitors (*HES1, NKX6-2*), presumptive OPCs (*OLIG1*), migrating clINs (*IGFBP5, HES6*), clINs (*LHX6, DLX5, GAD1*), hyINs (*LHX9*), hiINs (*FEV, GATA3*) and mature clINs (CNTNAP2). clIN, cortical-like interneurons; hyIN, hypothalamic-like interneurons; hiIN, hindbrain-like interneurons; OPC, oligodendrocyte precursor cell; hvFO, human ventral forebrain organoid; MGE/CGE/LGE, medial/caudal/lateral ganglionic eminence. **D.** Dot plot of marker gene expression across the identified populations as mentioned in C. Dot size indicates the percentage of cells per cluster expressing each marker. **E.** Heatmap of DLX5/6^+^ cINs and clINs expressing selected genes associated across mouse, human, and hvFO datasets used on Figure 1. Gray indicates expression below the 5% detection threshold. **F.** Label transfer of DLX5/6^+^ hvFO cells onto a human GE reference atlas with annotation for MGE-, CGE-, and LGE-enriched regions. **G.** Western blot analysis of extracts from 30 days hvFOs derived from three control (CCP1^WT#1^, CCP1^WT#2^, and CCP1^WT#3^) and three CCP1 loss of function “mutant” (CCP1^Mut#1^, CCP1^Mut#2^, and CCP1^Mut#3^) hiPSC lines. CCP1 and polyglutamylated tubulin (PolyE) expression were analyzed, and β-actin was used as a loading control. **H-J**. Quantitative analysis of nucleokinesis in migrating clINs within 90 days old human forebrain assembloids (hFAs). hvFOs were transduced with Lv-Dlx5/6-eGFP before fusion. Violin plots show mean migration speed (H), nucleokinesis amplitude (I), and total nuclear pause duration (J) (n=5-15 cells from 3-4 independent differentiation experiments; one-way ANOVA with Tukey’s multiple-comparison test; ns, not significant; *p< 0.05). **K**-**M**. Quantitative analysis of branching dynamics in clINs from 90-day hFAs. Violin plots show growth cone splitting frequency (K), maximum leading process length (L), and maximum number of secondary neurites (M). Data from all CCP1^WT^ lines and all CCP1^Mut^ (loss-of-function) lines were pooled according to CCP1 status (n=4 independent cell cultures; unpaired t test ; ns, not significant ; **p< 0.001, ***p< 0.001).

***Figure S4. Proteomic analysis identifies pathways altered upon CCP1 loss in human cortical-like interneurons***

**A*.*** Dot plots of the top 10 dysregulated proteins across all categories. Dot size indicates statistical significance (-log₁₀ P), and color represents the magnitude and direction of differential expression (log₂ fold change). **B.** Ingenuity Pathway Analysis (IPA) of significantly enriched canonical pathways identified from the differentially expressed proteins. Bar colors show the corresponding biological processes, as indicated. **C.** IPA of significantly enriched canonical pathways identified from the differentially expressed proteins. Bar colours show the corresponding biological processes, as indicated. **D**. RT-qPCR analysis showing *DCX* mRNAs from extracts of 30 days old ventral forebrain organoids (hvFOs) derived from hiPSC lines, as indicated. Kruskal-Wallis test followed by Dunn’s multiple comparison post-hoc test. Ns, no significative. **E-F.** Western blot analyses of protein extracts from 30 days old ventral forebrain organoids (hvFOs) derived from two control hiPSC lines (CCP1^WT#2^ and CCP1^WT#3^) and three CCP1 loss-of-function hiPSC lines (CCP1^Mut#1^, CCP1^Mut#2^, CCP1^Mut#3^). DCX protein levels were assessed, and β-actin was used as loading control (D). Western blot analysis comparing DCX expression level (β-actin as loading control) between 30 days old hvFOs derived from CCP1^WT#2^, CCP1^Mut#2^, and CCP1^Iso#2^(E). **G**. RT-qPCR analysis showing *DCX* mRNAs from E13.5 mouse MGE extracts, genotype as indicated. **H.** Western blot analysis of DCX expression in E13.5 mouse MGE extract, genoytype as indicated. **I**. Histogram showing average ratios between DCX and β-actin, as reported in H. **J.** Graph bar showing the percentage of eGFP-hDCX after co-transfection of HEK 293 cells with either E1Fa-tdTomato-shCTR or E1Fa-tdTomato-shDCX#1/shDCX#2 (One-Way ANOVA, followed by Tukey’s multiple comparisons post-hoc test. n=3 independent cell cultures, ***p< 0.001).

***Figure S5. Mitochondrial and bioenergetic analyses in CCP1-deficient ventral forebrain organoids***

**A.** Dot plot of the top differentially expressed proteins associated with the IPA mitochondrial dysfunction category upon loss of CCP1. Dot size corresponds to statistical significance (-log₁₀ P), whereas their color indicates the magnitude and direction of differential expression (log₂ fold change). **B.** Oxygen consumption rate (OCR) measured in 90 days old human ventral forebrain organoids (hvFOs). Data are presented as mean ± SEM from three biological replicates (Ordinary one-way ANOVA followed by Tukey’s multiple comparisons test, Ns no significative). **C.** Representative Western blot showing expression level of oxidative phosphorylation (OXPHOS) complex proteins in extracts from 60 days old hvFOs. Cell line identity, as indicated. NDUFB8, complex I; SDHB, complex II ; UQCRC2, complex III ; COXII, complex IV ; ATP5A, ATP synthase ; VDAC1, voltage-dependent anion channel 1. **D**-**G**, quantification of OXPHOS protein levels from 3 independent Western blots (a representative one is shown in C). Ordinary one-way ANOVA followed by Tukey’s multiple comparisons post-hoc test; ns, no significant **H.** Experimental workflow for live labeling of mitochondria in cortical like-interneurons (clINs) migrating out of hvFOs. At 60 days, hvFOs were co-transduced with Lv-Dlx5/6-eGFP and Lv-mito-mCherry lentivirus prior to live imaging. **I.** Representative images of migrating clINs derived from CCP1^WT#2^ and CCP1^mut#2^ hvFOs expressing mito-mCherry. A fire lookup table (ImageJ) was applied to visualize mitochondrial fluorescence intensity during their movement. Scale bar, 20 μm. **J-M.** Quantitative analysis of mitochondrial dynamics in migrating clINs. Violin plots show number of mitochondria per cell (J), total mitochondrial area per cell (TMA) (K), mitochondrial fusion rate (fusion events per minute) (L), and mitochondrial fission rate (fission events per minute) (M). Data represent n=2-5 cells per hvFO from three independent differentiations (unpaired t test; ns, not significant; *p < 0.05). **N.** Quantification of the proportion of total mitochondrial area (TMA) localized within the soma (left) or neurites (right) over the course of nucleokinesis. Data represent n=2-5 cells per hvFO, from three independent differentiations (two-way ANOVA with Tukey's multiple-comparison post-hoc test; ns, not significant).

***Figure S6. CCP1 overexpression alters the migratory behavior of mouse cortical interneurons***

**A.** Immunolabeling of PolyE (magenta) in mouse cortical interneurons (cINs) overexpressing CCP1 (Dlx5/6-CCP1^wt^) or control ones (Dlx5/6-CCP1^dead^). Scale bar, 20 μm. **B.** Representative time-lapse sequence of a migrating cINs from E13.5 mouse MGE explants transduced with Lv-Dlx5/6-eGFP together with either Lv-CAG-mCCP1-Cherry (CCP1^wt^) or Lv-CAG-mCCP1dead-Cherry (CCP1^dead^). Scale bars, 20 μm. **C-E.** Quantitative analysis of nucleokinesis dynamics in CCP1^wt^ and CCP1^dead^ mouse cINs. Violin plots show mean migration speed (C), nucleokinesis frequency (D), and nuclear pause duration (E). Data represent n=15–30 cells from 4-5 embryos across three independent experiments (unpaired t test; ns, not significant; *p<0.05; ***p<0.001). **F-H.** Quantitative analysis of branching dynamics in CCP1^wt^ and CCP1^dead^ mouse cINs. Violin plots show maximal leading process length (F), maximal secondary neurite length (G), and maximum number of neurites per cell (H). Data represent n=15-30 cells from 4-5 embryos across three independent experiments (unpaired t test; ns, not significant; *p<0.05; ***p<0.001). **I**. Computational model showing cortical invasion by interneuron surrogates for human-like, CCP1^wt^-mouse like and Mouse-like ones, numbers as indicated on panels. **J**-**K**. Immunolabeling of HEK293 cells co-transduced with Lv-pCMV-Cre and Lv-loxStoploxCCP1^wt^-Cherry or Lv-loxStoploxCCP1^Dead^-Cherry showing CCP1 (grey), mCherry (red) and nuclei counterstained with Dapi (blue) (J). Immunolabeling of HEK293 cells co-transduced with pCMV-Cre and loxStoploxCCP1^WT^-Cherry or loxStoploxCCP1^Dead^-Cherry showing polyglutamylation (PolyE, grey), mCherry (red) and nuclei counterstained with Dapi (blue) (K). **L**. Immunolabeling of CCP1 (grey) in transfected cIN (mCherry positives, red) by in IUE of mouse forebrain at E17.5. Scale bars, (J) 50 µm, (K) 50 µm, and (L) 10µm. **M**. Apico-basal distribution of endogenous non IUE cINs that express Dlx5/6-eGFP in E17.5 mice subjected to IUE with CCP1^wt^ or CCP1^dead^ constructs. **N**, **O**. Representative immunolabeling images showing Cxcl12/CXC12 expression (purple) as gradients across the cortical wall and as hotspots around blood vessels, together with Lhx6 expression (yellow) in cINs, in human GW12 and mouse E13.5 cortical walls. Scale bars, 100µm.

***Figure S7. Predicting the Molecular Drivers of Homotypic repulsion Among Cortical Interneurons Across Species***

**A**-**B.** UMAP of integrated mouse (A) or human (B) single-cell RNA sequencing data from 19 independent datasets (*6-13*) and spanning mouse embryonic days 13, 13.5 and 14 (36,467 cells) (A) or human fetuses at gestational week 13 (21,823 cells) annotated by cell populations, as reported in the figure. Apical progenitor, AP; intermediate progenitor, IP; projection neuron, PN; cajal retzius cell, CR; interneuron, IN; oligodendrocyte precursor cell, OPC; outer radial glia, oRGC. Circles higlight relevant cell clusters. **C.** Scheme illustrating molecular homotypic contact between migrating cortical interneurons (cINs). **D.** Venn diagram of cell-cell interaction drivers (p < 0.01) inferred from cIN (MGE + CGE INs) signaling in mouse and human datasets, identifying mouse-specific, human-specific, and shared molecular interactions, using CellChat. **E.** Dot plot of all shared cell-cell interaction drivers shared between mouse and human cINs. **F.** Dot plot showing the top 10 human-specific cell-cell interactions, ranked by communication probability. Dot colour indicates communication probability, and dot size indicates statistical significance (p < 0.01).

***Figure S8. Ferret cortical interneurons exhibit migratory dynamics intermediate between mouse and human***

**A.** Representative time-lapse sequence of a migrating cIN derived from E36 ferret MGE explants and transduced with Dlx5/6-eGFP lentivirus. Scale bar, 20 μm. **B-E.** Comparative analysis of the migration dynamics of E13.5 mouse, E36 ferret, and GW12 human cINs in MGE explants. Violin plots show mean migration speed (B), nucleokinesis frequency (C), nuclear pause duration (D), and growth cone splitting frequency (E). Data represent n=15-30 cells from 4-5 embryos or fetuses across three independent experiments (one-way ANOVA with Tukey's multiple-comparison post-hoc test; ns, not significant; *p<0.05; ***p<0.001). **F.** Representative Sholl analysis of a migrating cIN at the stage of maximum branching complexity (n=15-30 cells from 4-5 embryos or fetuses; comparison ferret to mouse, two-way ANOVA with Tukey's multiple-comparison post-hoc test; *p<0.05, **p<0.01, ****p<0.0001). **G.** Quantification of total neurite intersections at maximum branching complexity in migrating cINs out of MGE explants from mouse (E13.5), ferret (E36), and human (GW12) (n=15-30 cells from 4-5 embryos or fetuses; one-way ANOVA with Tukey's multiple-comparison post-hoc test; ***p<0.001). **H.** Quantification of homotypic contact frequency between migrating cINs in mouse (E13.5), ferret (E36), and human (GW11-12) migrating out of MGE explants. Violin plot shows the number of contacts per hour (n=25-30 cells from three independent experiments; one-way ANOVA with Tukey's multiple-comparison test; ns, not significant; *p<0.05). **I.** Pie chart showing the frequency of repulsive and neutral (non-repulsive) homotypic contacts between migrating ferret cINs (n=27 cells from three independent experiments). **J.** Representative immunolabeling showing the distribution of LHX6⁺ cINs in embryonic E36 ferret cortical sections. Scale bar, 100µm **K.** Quantification of the apico-basal cIN distribution in embryonic mouse (gray), ferret (purple), and human (green) cortical sections (two-way ANOVA with Tukey's multiple-comparison test ; p*<0.05; n=3-4 embryos or fetuses per group) **L.** Immunoblot analysis of MGE lysates from E36 ferret embryos showing CCP1 expression. α-tubulin is the loading control. **M.** Quantification of CCP1 protein levels normalized to α-tubulin and expressed relative to mouse levels (n=4-5 samples per group ; unpaired t test ; p*<0.05). **N.** Immunoblot analysis of MGE lysates from E36 ferret embryos showing DCX expression. βIII-tubulin was used as a loading control. **O.** Quantification of DCX protein levels normalized to βIII-tubulin and expressed relative to mouse levels (n=4-5 samples per group; unpaired t test ; p*<0.05).

**Supplemental movies**

***Movie S1. Related to Fig. 1.*** Time lapse video microscopy performed with confocal microscope (Airyscan, Zeiss, 10X) on organotypic slices from E13.5 mouse forebrain transduced with Lv-DLX5/6-eGFP showing the migration of cINs in the developing cortical wall. Duration of recording is 8h.

***Movie S2. Related to Fig. 1.*** Time lapse video microscopy performed with confocal microscope (Airyscan, Zeiss, 10X and 1.6 zoom) on organotypic slice from GW12 human forebrain transduced with Lv-DLX5/6-eGFP showing the migration of cINs in the developing cortical wall. Duration of recording is 8h.

***Movie S3. Related to Fig. 1.*** Time lapse video microscopy performed with confocal microscope (Airyscan, Zeiss, 20X) on cINs migrating from MGE explant of E13.5 mouse forebrain transduced with Lv-DLX5/6-eGFP showing the migration of cINs. Duration of recording is 9h.

***Movie S4. Related to Fig. 1.*** Time lapse video microscopy performed with confocal microscope (Airyscan, Zeiss, 20X) on cINs migrating from MGE explant of GW12 human forebrain transduced with Lv-DLX5/6-eGFP showing the migration of cINs. Duration of recording is 17h.

***Movie S5. Related to Fig. 2.*** Time lapse video microscopy performed with confocal microscope (Airyscan, Zeiss, 20X) on shCTR-targeted cINs migrating from MGE explant of GW12 human forebrain transduced with Lv-DLX5/6-eGFP showing the migration of cINs. Duration of recording is 16.

***Movie S6. Related to Fig. 2.*** Time lapse video microscopy performed with confocal microscope (Airyscan, Zeiss, 20X) on shCCP1#1-targeted cINs migrating from MGE explant of GW12 human forebrain transduced with Lv-DLX5/6-eGFP showing the migration of cINs. Duration of recording is 16h.

***Movie S7. Related to Fig. S8.*** Computational modelling of cIN surrogate migration into the cortical wall of human (left), mouse (middle), and mouse with increase leading process branhing dynamcis and mutual repulsive contacts (mGoF).

***Movie S8. Related to Fig. S8.*** Time lapse video microscopy performed with confocal microscope (Andor Dragonfly Spinning Disc, 20X) on cINs migrating from MGE explant of E36 ferret forebrain transduced with Lv-DLX5/6-eGFP showing the migration of cINs. Duration of recording is 14h.
